# *Pseudomonas aeruginosa* R-bodies deliver an ADP deaminase toxin

**DOI:** 10.64898/2026.10.21.760852

**Authors:** Devin T. King-Roberts, Kiavash Garakani, Thomas Y. Tao, Hsin-Cheng Lin, Joshua Powell, Tongxin Wu, Eugenia J. Lee, Tiffany M. Yang, Mariia Serebriakova, Alexa Price-Whelan, Virginia W. Cornish, Liang Tong, Lars E.P. Dietrich

**Affiliations:** Department of Biological Sciences, Columbia University; New York, NY USA; Integrated Program in Cellular, Molecular, and Biomedical Studies, Columbia University Medical Center; New York, NY USA; Department of Chemistry, Barnard College; New York, NY USA; Department of Chemistry, Columbia University; New York, NY USA; Department of Systems Biology, Columbia University; New York, NY USA

## Abstract

Diverse bacteria produce proteinaceous polymeric coils called R-bodies (“refractile bodies”), which extend into ∼10 µm-long needles in response to environmental triggers. It has been proposed that R-bodies deliver toxins intracellularly after such bacteria are ingested by eukaryotes, but intra-host extension of R-bodies has not been directly observed and these toxins remain unidentified. We use *Pseudomonas aeruginosa* PA14 and *Dictyostelium discoideum* to show that R-bodies extend within amoebae. We also identify a novel toxin, DeaR (<u>de</u>aminase <u>a</u>ssociated with the <u>R</u>-body), which contributes to predation resistance and forms a near-megadalton dodecamer with ADP deaminase activity that alters purine metabolism. Our findings elucidate the molecular basis of R-body-mediated toxicity during PA14 infection and provide a paradigm for understanding the potential of <u>c</u>ytotoxic <u>e</u>ffectors <u>d</u>elivered by <u>R</u>-bodies (CEDRs) in interkingdom interactions.

## Main text

Bacteria in ecosystems face a variety of biological threats including phage infection and eukaryotic predation. Common tools in bacterial phage defense systems are deaminases that modify nucleotides to disable viral propagation (*1–5*), while escape from the eukaryotic phagolysosomephagolysosome is achieved by disrupting the host’s intracellular membranes (*6*, *7*). Here, we describe a defense mechanism that combines deaminase toxicity with phagolysosomal membrane damage to confer bacterial resistance against predation. Phylogenetic analyses suggest that this mechanism operates in diverse bacterial species to protect their populations from phagocytes, whether free-living or present within larger multicellular eukaryotes.

Our group recently discovered that *Pseudomonas aeruginosa* strain PA14 (*8*) has the capacity to produce large hyperstable protein polymers called R-bodies (for “refractile bodies”) (*9*). R-bodies were first identified in *Caedibacter taeniospiralis*, a bacterial endosymbiont of some paramecia (*10–12*). *Paramecium* strains that harbor *Caedibacter* shed the bacteria into the environment, where they can be phagocytosed by and are toxic to other, “sensitive” *Paramecium* strains. Although R-body production has been linked to host toxicity in studies of *Caedibacter* spp., *Azorhizobium caulinodans*, and PA14 (*9*, *13*, *14*), observations have also suggested that R-bodies alone are insufficient to effect this toxicity. These include the findings that either (i) purified *C. taeniospiralis* R-bodies, or (ii) an *E. coli* strain engineered to produce these R-bodies, are not toxic to sensitive paramecia (*14–16*). Purified R-bodies undergo a rapid (as fast as 100 ms) and drastic morphological change from a coiled roll to an extended needle at the transition from neutral to acidic pH (*9*, *17*, *18*); a popular model for the mode of R-body-dependent toxicity has therefore been that R-bodies serve to deliver yet unidentified bacterial toxins by disrupting the membrane of the maturing phagolysosome (*19*). However, this model has not been tested since its first proposition several decades ago (*20*).

*C. taeniospiralis* contains a plasmid with four structural genes sufficient for R-body production in *E. coli*, but the plasmid does not code for any compelling toxin candidates (*14*, *21*, *22*). By contrast, the R-body structural genes in PA14 are situated in a chromosomal locus called the *reb* cluster (**Figure 1A**), which includes several uncharacterized genes. In this study, we sought to probe the model that PA14 R-bodies extend within the host cell after phagocytosis and test the hypothesis that a *reb*-cluster-encoded protein confers host toxicity. We used the amoeba *Dictyostelium discoideum*, a model host that cohabits with pseudomonads in soil (*23*, *24*), to investigate the roles of *reb* cluster genes in predation resistance. We observed that R-bodies and other products of the *reb* cluster help to protect PA14 from *D. discoideum* and that R-body extension is triggered after *D. discoideum* phagocytosis. Importantly, we found that a protein encoded by the *reb* cluster gene *PA14_27675* is toxic to mammalian cells, thereby identifying PA14_27675 as a long-sought toxin associated with R-bodies. Structural and biochemical characterization of PA14_27675 revealed that it forms a near-megadalton-sized dodecamer with ADP deaminase activity, and we have therefore named the protein DeaR (<u>dea</u>minase associated with the <u>R</u>-body). Our phylogenetic analyses indicate that deaminases are among a diverse set of putative toxins encoded by *reb*-cluster-associated or -adjacent genes in bacterial species capable of R-body production. Together, these observations imply that R-body-facilitated delivery of toxins, including deaminases, may be a widespread strategy employed by bacterial populations to resist phagocytosis

**Figure 1.**
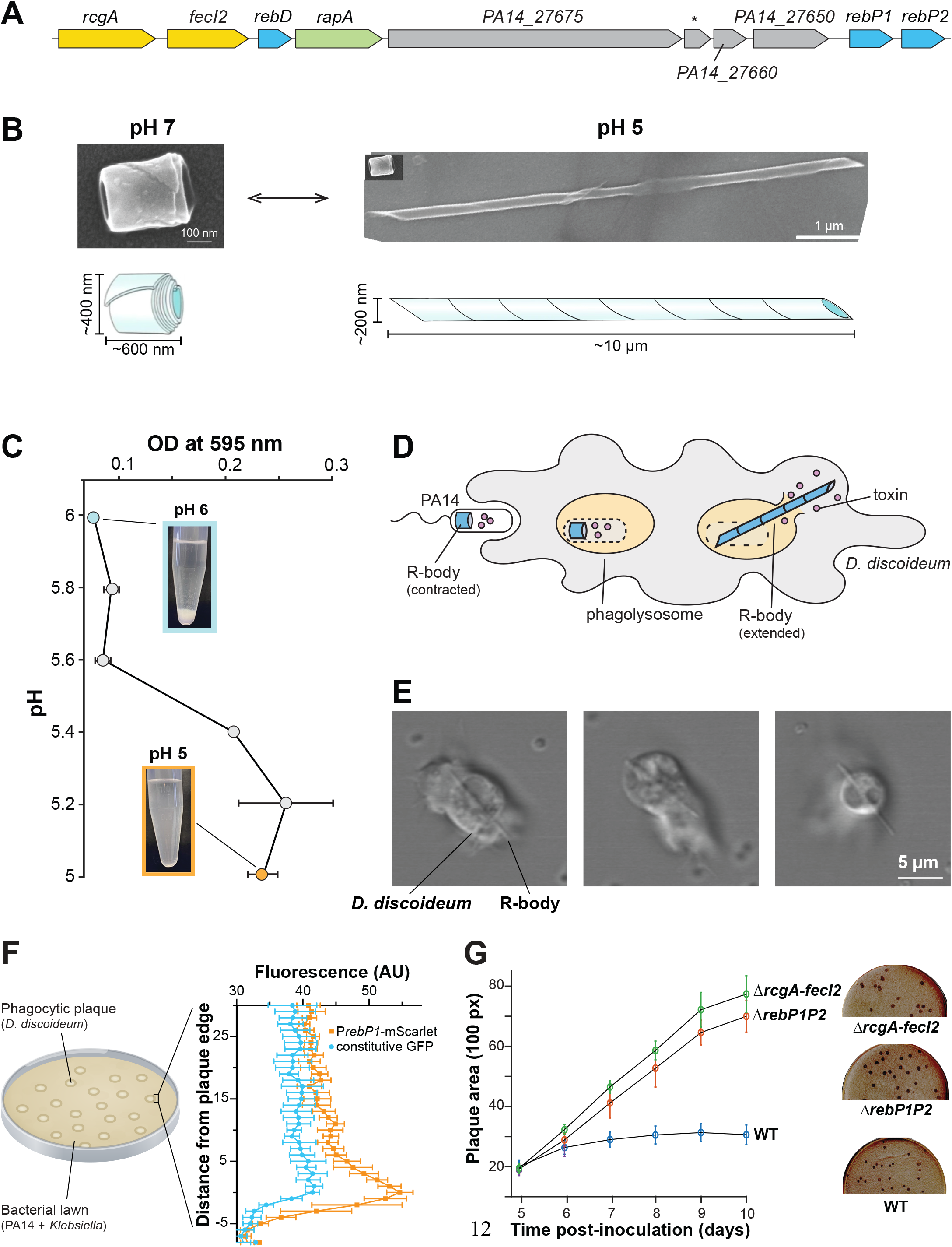
R-bodies are hypothesized to deliver toxins to phagocytic cells. **(A)** The PA14 *reb* cluster. Yellow, blue, and gray indicate regulatory, structural, and uncharacterized genes, respectively. Genes between *fecI2* and *rapA,* and *PA14_27675* and *PA14_27660* are annotated in BioCyc (*1*) but not the Pseudomonas Genome DB. **(B)** SEM images of coiled and extended R-bodies and schematics showing their dimensions. **(C)** OD_595_ of supernatant when R-body suspensions are adjusted to the indicated pH. Points represent averages and error bars represent SD. **(D)** Putative role of R-bodies in bacterial toxin delivery. After engulfment, lysosomal maturation promotes dissolution of the bacterial cell wall and decreases the pH of the phagosome. Low pH triggers R-body extension, releasing phagolysosomal contents (including bacterial toxins) into the eukaryotic cytoplasm. **(E)** Representative images showing *D. discoideum* incubated with PA14, displaying protruding R-bodies. **(F) Left:** Toxicity for *D. discoideum* is assessed by quantifying the clearings in a bacterial lawn caused by amoebal predation. **Right:** Average fluorescence and SD for traces taken across the plaque boundary around n≥12 plaques on 3 biological replicates of PA14. **(G)** Plaque area over time in lawns containing PA14 WT or mutants lacking the indicated *reb* cluster genes. Data represent the average of n≥8 plaques measured from 3 biological replicates with SD. Representative images of plates are shown on the right.

## Results

### PA14 R-bodies undergo pH-responsive extension

The PA14 *reb* cluster contains two regulatory genes (*rcgA* and *fecI2*), three genes homologous to *C. taeniospiralis* R-body structural genes (*rebD*, *rebP1*, and *rebP2*), one gene implicated in R-body assembly (*rapA*), and four uncharacterized genes (**Figure 1A**) (*9*). Using fluorescent constructs that report transcription, we have observed that genes in this cluster are stochastically expressed such that a small proportion (0.3-5%, depending on the growth condition) of cells produce R-bodies (*9*). We have also used fluorescent protein fusions to show that Reb proteins form an intracellular cylindrical structure. To further characterize the structure and activity of PA14 R-bodies, we created a strain that expresses the positive regulator of the *reb* cluster under an inducible promoter and used it to prepare enriched R-bodies at neutral pH. We buffer-exchanged an aliquot to acidic pH, then imaged both preparations by scanning electron microscopy (SEM). While the contracted R-bodies from the neutral-pH preparation resembled tightly coiled ribbons (**Figure 1B, figure S1A**), the acidic-pH preparation contained extended R-bodies, most of which were likely deformed by the fixation and/or dehydration steps required to prepare samples for SEM (**Figure 1B, figure S1B**). The intact, extended R-bodies that were visible resembled long (8.1 ± 0.9 µm) needles with acute ends, reminiscent of the so-called type 51 R-bodies such as those produced by *C. taeniospiralis* (*16*, *19*, *25*, *26*). We further analyzed R-body extension by exploiting the fact that contracted R-bodies sediment, whereas extended R-bodies remain in suspension. In this assay, we resuspended contracted R-bodies in buffers across a range of pH values and allowed them to sediment before measuring the optical density of the unsedimented fraction. This assay revealed that PA14 R-bodies extend at pH values below 5.6 (**Figure 1C**), similar to *C. taeniospiralis* R-bodies, which extend at pH values below 5.7 (though this value can shift depending on the ionic strength of the assay buffer system (*19*)).

### PA14 reb cluster products respond to and hinder phagocytic predation

For decades, the prevailing hypothesis about R-body biology has been that these coiled intrabacterial structures extend when bacteria are exposed to the conditions of the maturing host phagolysosome. This process destroys the R-body-containing bacterium and theoretically disables the phagocyte by releasing lysosomal contents and potential toxins present in the bacterium into the host cytoplasm (**Figure 1D**). In *Caedibacter* spp., this mechanism is thought to confer a competitive advantage on host paramecia that harbor the endosymbiotic bacteria and shed them for sensitive competitors to phagocytose (*11*). In contrast, in free-living bacteria, a small, stochastically differentiated subpopulation produces R-bodies that extend upon ingestion and damage the phagocytic predator. Because such a predator can consume up to 200 bacteria per hour (*27*), disabling one predatory individual lowers grazing pressure on bacterial clonemates, disproportionately to the number of bacterial cells lost.

The fact that R-bodies extend at low pH is consistent with the model that they are triggered to extend in the phagolysosome (*28*), and an early micrograph that appeared to show extension inside a sensitive paramecial host also supported this idea (*29*). In a more recent study, Vasquez-Rifo *et al.* examined the role of the PA14 *reb* cluster in host toxicity using *Caenorhabditis elegans*, a nematode that preys on pseudomonads in soil (*30*). They found that *C. elegans* subjected to a dynamin knockdown, which disables bacterial uptake in the intestine, is protected from toxic effects mediated by the *reb* cluster. This observation further indicated that engulfment of R-body-containing bacteria is required for the R-body and/or its associated genes to elicit host damage. Together, these observations represent the limited evidence to date supporting the model that R-body extension occurs in the phagolysosome and serves as a delivery mechanism for bacterial toxins.

To directly investigate the roles of *reb* cluster products during phagocytosis of *P. aeruginosa*, we turned to the model amoebal host *Dictyostelium discoideum*. First, we examined whether R-body extension occurs after live cells engulf bacteria. We combined axenic cultures of *D. discoideum* Ax4 with a nonpathogenic strain of *Klebsiella* (which serves as food for the amoebae) and PA14 expressing the positive regulator of the *reb* cluster under a constitutive promoter (RcgA+) in chamber slides, and imaged them using confocal microscopy at 1.07 frames/second. After 30 minutes of incubation, we observed needle-like structures protruding from live *D. discoideum* (**Figure 1E and movie S1**).

Next, to investigate the expression and effects of the *reb* cluster during interactions between PA14 populations and *D. discoideum*, we adapted a “plate killing” assay developed by Pukatzki *et al.* (*31*). In this assay, *P. aeruginosa* PA14, *D. discoideum*, and the *Klebsiella* food bacteria are mixed and plated on an agar-solidified medium. The bacteria form a lawn, and amoebae consume them, forming plaques (clearings) in this lawn. To examine whether PA14 produces R-bodies under these conditions, we used a strain in which expression of the fluorescent protein mScarlet is driven by a copy of the *rebP1* promoter (*9*). We grew this strain and a strain that constitutively produces GFP in the plate killing assay and imaged using fluorescence scanning. Interestingly, we observed enhanced mScarlet fluorescence at the edges of the clearings, specifically in lawns containing the P*_rebP1_* construct (**Figure 1F**), whereas constitutive GFP expression did not increase significantly. This result suggests that proximity to *D. discoideum* induces *reb* gene expression in PA14. Finally, to test whether the *reb* cluster and R-body production affect bacterial predation by *D. discoideum*, we used the plate killing assay and two mutants: Δ*rebP1P2*, which is unable to produce the major structural components of the R-body, and Δ*rcgAfecI2*, which lacks regulatory proteins required for *reb* cluster expression. Consistent with the observation in *C. elegans* that the *reb* cluster and R-bodies contribute to PA14 virulence (*9*), we found that lawns containing mutants lacking *rcgAfecI2* or *rebP1P2* formed larger plaques than lawns containing WT PA14 (**Figure 1G**). This indicates that the Δ*rcgAfecI2* and Δ*rebP1P2* mutants are more readily consumed by *D. discoideum* and, therefore, that the *reb* cluster and R-bodies contribute to PA14 predation resistance.

### PA14_27675 encodes a putative nucleotide deaminase that contributes to phagocyte resistance

The *reb* cluster contains four genes between *rapA* and *rebP1* (**Figure 1A**). The two genes upstream of *rebP1*, *PA14_27660* and *PA14_27650*, encode proteins of 8 kDa and 20 kDa, respectively. Upstream of *PA14_27660* is an open reading frame that is not universally detected by annotation software, but that we refer to as *PA14_27670*; its product is 7 kDa. None of these uncharacterized gene products share similarity (based on AlphaFold 3 (AF3) prediction (*32*)) with any characterized proteins in sequence and structure databases. Most notable, however, is the fourth uncharacterized product of the *reb* cluster, PA14_27675 (“DeaR”), an ∼80-kDa protein. The AF3-modeled structure of DeaR contains a TIM β-barrel domain, a motif commonly associated with enzyme activity (*33*, *34*) (**Figure 2A and file S1**). Using Foldseek (*35*), we found that DeaR shows the highest confidence overlap with solved structures of adenosine deaminases. These enzymes catalyze the conversion of adenosine’s primary amine group to a carbonyl. Specifically, Zn²⁺ in the deaminase active site coordinates and activates a water molecule, generating a nucleophilic base that attacks the adenine ring and promotes hydrolytic deamination (*36–39*). Alignment of the active site residues of a known adenosine deaminase from *Mus musculus* (PDB: 1UIP) (*40*) to the putative active site of DeaR further revealed an identical configuration of residues (one aspartate and three histidine residues) implicated in metal-cofactor binding (**Figure 2A and file S1**).

**Figure 2.**
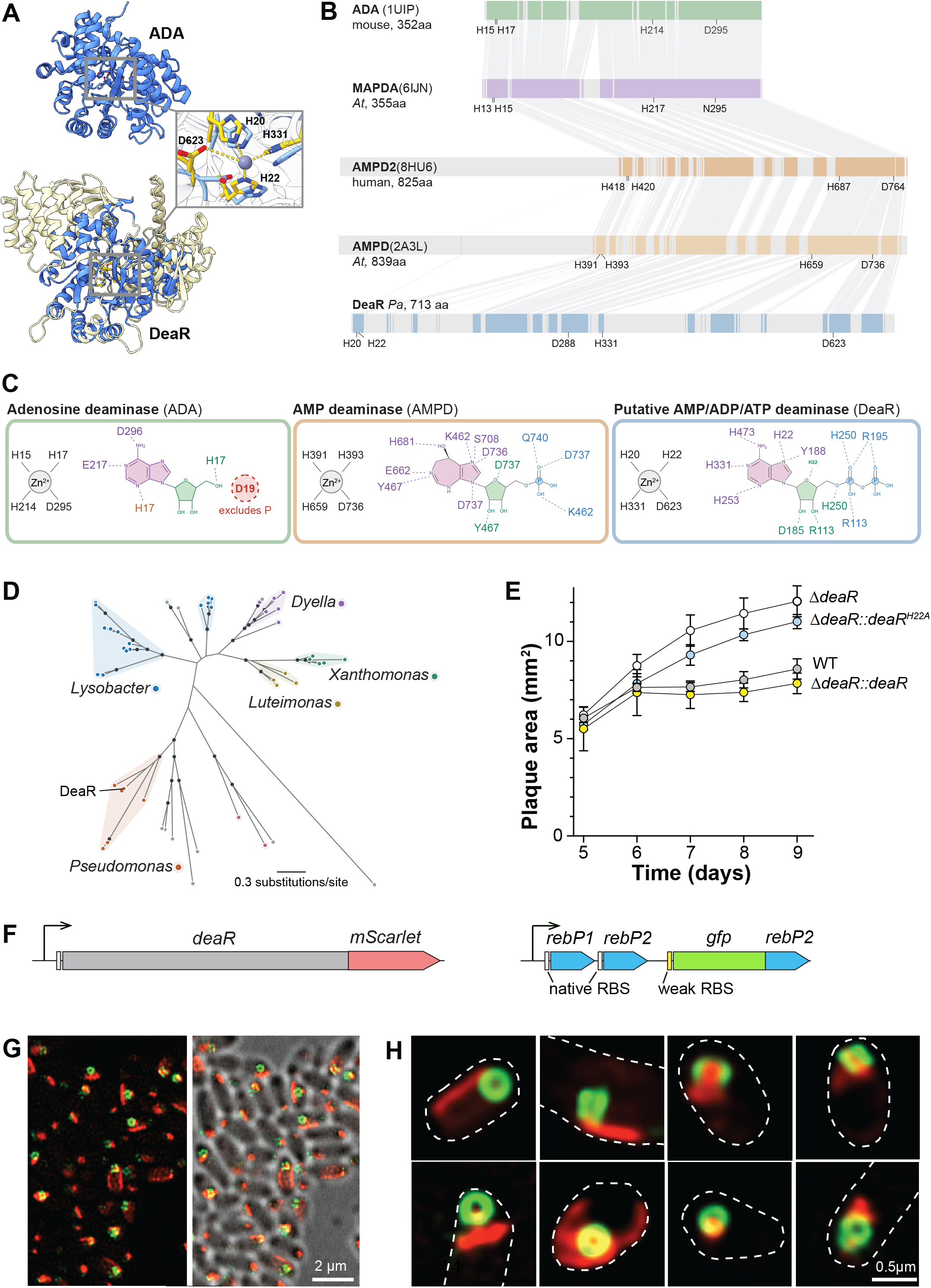
*PA14_27675* (*deaR*) codes for a putative deaminase that contributes to predation resistance and associates with the R-body. **(A)** Alignment of an *M. musculus* adenine deaminase (**top**) and a model of PA14_27675 (DeaR) (**bottom**). Blue indicates the DeaR domain homologous to known deaminases. Inset shows an overlay of active sites with highlighted key residues (numbering from DeaR). Residues in blue and gold are from 1UIP and DeaR, respectively. **(B)** Linear protein diagram showing regions (shaded) of DeaR that are homologous to characterized deaminases. Key active-site and conserved residues are indicated. *Pa*, *Pseudomonas aeruginosa*. *At*, *Arabidopsis thaliana*. MAPDA, N6-methyl-AMP deaminase. **(C)** Schematics of the deaminase types shown in (B) with Zn^2+^ and potential substrates (**left panel:** adenosine; **right panel:** ADP) or the AMP deaminase inhibitor coformycin 5-phosphate (**middle panel**). Active-site residues associated with Zn^2+^ or the substrate are depicted. Red indicates that D19 prevents mouse ADA from binding adenosine nucleotides. **(D)** Phylogenetic tree of PA14_27675 homologs identified using NCBI BLASTP. Colored shading indicates grouping by genus. **(E)** Average plaque size produced by *D. dictyostelium* predation in bacterial lawns containing PA14 WT, the Δ*deaR* mutant, and Δ*deaR* strains complemented with either WT DeaR or DeaR H22A. Data represent the average of n≥4 plaques measured from each of 3 biological replicates with SD. **(F)** Fusion constructs used to visualize intracellular distribution of DeaR and RebP2. RBS, ribosome binding site. **(G)** Low-magnification fluorescence (**left**), fluorescence + DIC overlay (**right**), and high-magnification fluorescence **(H)** images of cells expressing DeaR and RebP2 fluorescent-protein fusions.

The TIM barrel domain and the active site of DeaR are formed by sequence segments conserved in shorter deaminase proteins, interspersed with unrelated sequence stretches (**Figure 2B**). Expanding our analysis to include additional proteins identified by Foldseek, we find plant and human proteins that are similar in size to DeaR but only show significant homology to the short deaminases over about one-half of the sequence. Like DeaR, these proteins also harbor conserved regions implicated in metal-cofactor and substrate binding, interspersed with unrelated sequences. These proteins are annotated as adenosine monophosphate deaminases. Looking in more detail at the active sites of both the short and long deaminases, we can see the high conservation of the metal cofactor-binding residues and parallels between the proteins in the interactions of active-site-oriented side chains with moieties on adenosine and AMP (**Figure 2C**). A crucial difference lies in the presence of an aspartate residue (Asp19) in adenosine deaminase, which excludes phosphorylated derivatives and is missing from both the characterized AMP deaminases (8HU6 and 2A3L), thereby allowing the active site to accommodate the phosphorylated form. Examining the AF3-generated structure of DeaR, we find that the putative active site also lacks this residue and shows the potential to accommodate additional phosphates, raising the possibility that DeaR could act on other adenosine derivatives.

Transformations of nucleosides and their derivatives, and nucleic acid editing, are common mechanisms by which bacteria damage other cells, both during interbacterial “warfare” and interkingdom interactions (*41–45*). Searching for DeaR homologs in other organisms using NCBI Blast, we identified similar proteins that are present in diverse genera of bacteria and found that many of the bacterial homologs are situated within 10 kb of homologs of *rebP1* from PA14 (**Figure 2D and figure S2**). These observations and the revelations from our structural modeling suggest that deaminases contribute to *reb* cluster-mediated toxicity. We therefore used the plate killing assay to test whether DeaR contributes to PA14 protection from *D. discoideum* predation and indeed found that the Δ*deaR* mutant exhibited larger plaque formation than WT PA14 and that chromosomal complementation of *deaR* restored the WT phenotype (**Figure 2E**). We also created a strain in which the gene coding for an H22A variant of DeaR was inserted at the complementation site, and found that this strain did not recapitulate the WT phenotype. This result indicates that the predicted Zn^2+^-coordinating residue H22 is required for DeaR function, consistent with our structural modeling (**Figure 2A,C**).

### R-bodies and DeaR colocalize within PA14 cells

The presence of DeaR-like deaminases in R-body-producing bacteria, the proximity of *deaR* to R-body structural genes, and the contribution of DeaR to defense against *D. discoideum* all supported the idea that the R-body and DeaR function as part of the same mechanism to confer toxicity in phagocytic host cells. To further probe the associations between DeaR and the R-body, we engineered a strain that produces fluorescently tagged DeaR and RebP2 in the RcgA+ background. Tagged RebP2 is produced using a construct in which an extra copy of *rebP2* is fused to *gfp* and translation is driven by a weak ribosomal binding site, so that only a portion of the RebP2 monomers in the R-body are labeled (decreasing potential structural perturbation by the label) (**Figure 2F**) (*9*, *19*). Liquid cultures of this strain were spotted onto an agar pad and imaged using a confocal microscope. We observed that DeaR and RebP2 tend to exist as discrete but proximal structures (**Figure 2G,H**). While RebP2 has a rigid ring morphology, DeaR forms a supramolecular assembly that shows variable morphology, including a common oblong shape, that is consistently in contact with the R-body. To probe this colocalization, we performed automated segmentation of images and, for each cell double-positive for RebP2-GFP and DeaR-mScarlet, computed the minimum distance between the two thresholded fluorescence signals. RebP2-GFP and DeaR-mScarlet fluorescence signals directly overlapped in 89.3% of instances in 1,041 analyzed cells, and were within 1 pixel of each other in 93.9% of instances. The formation of a supramolecular assembly by DeaR might serve to confer self-protection by modulating its enzymatic activity (*46–50*).

### Heterologous DeaR expression modulates nucleotide pools within mammalian cells

Having observed that DeaR contributes to predation resistance and is spatially associated with the R-body, we hypothesized that this protein is delivered by the R-body into the eukaryotic cytosol, where its deaminase activity damages the host. We sought to test the effects of DeaR activity on host physiology, independent of delivery, by expressing the enzyme directly in the cytosol of a eukaryote. We therefore generated constructs in pINDUCER21, a doxycycline-inducible lentiviral vector carrying a constitutively expressed eGFP reporter (*51*), for expression of DeaR and the DeaR H22A variant. Inducible expression of DeaR was driven by the TRE2 promoter, consisting of seven *tet* operator elements upstream of a minimal CMV promoter. Given the propensity of the CMV promoter to undergo silencing in hematopoietic-lineage cells (*52*), we elected to evaluate DeaR expression in the non-hematopoietic HeLa cell line. We verified the incorporation and doxycycline-inducible expression in HeLa cells using flow cytometry, RT-qPCR, and RNA sequencing (**figure S3A,S3B**). Using a commercial assay kit to assess viability, we found that across a range of doxycycline doses, induction of WT DeaR reduced HeLa viability at 48 h by ∼80% (**Figure 3A**), whereas DeaR H22A, expressed at comparable levels (**figure S3B**), showed a minor effect on viability (∼15%). These results indicate that DeaR catalytic activity is toxic to mammalian cells.

**Figure 3.**
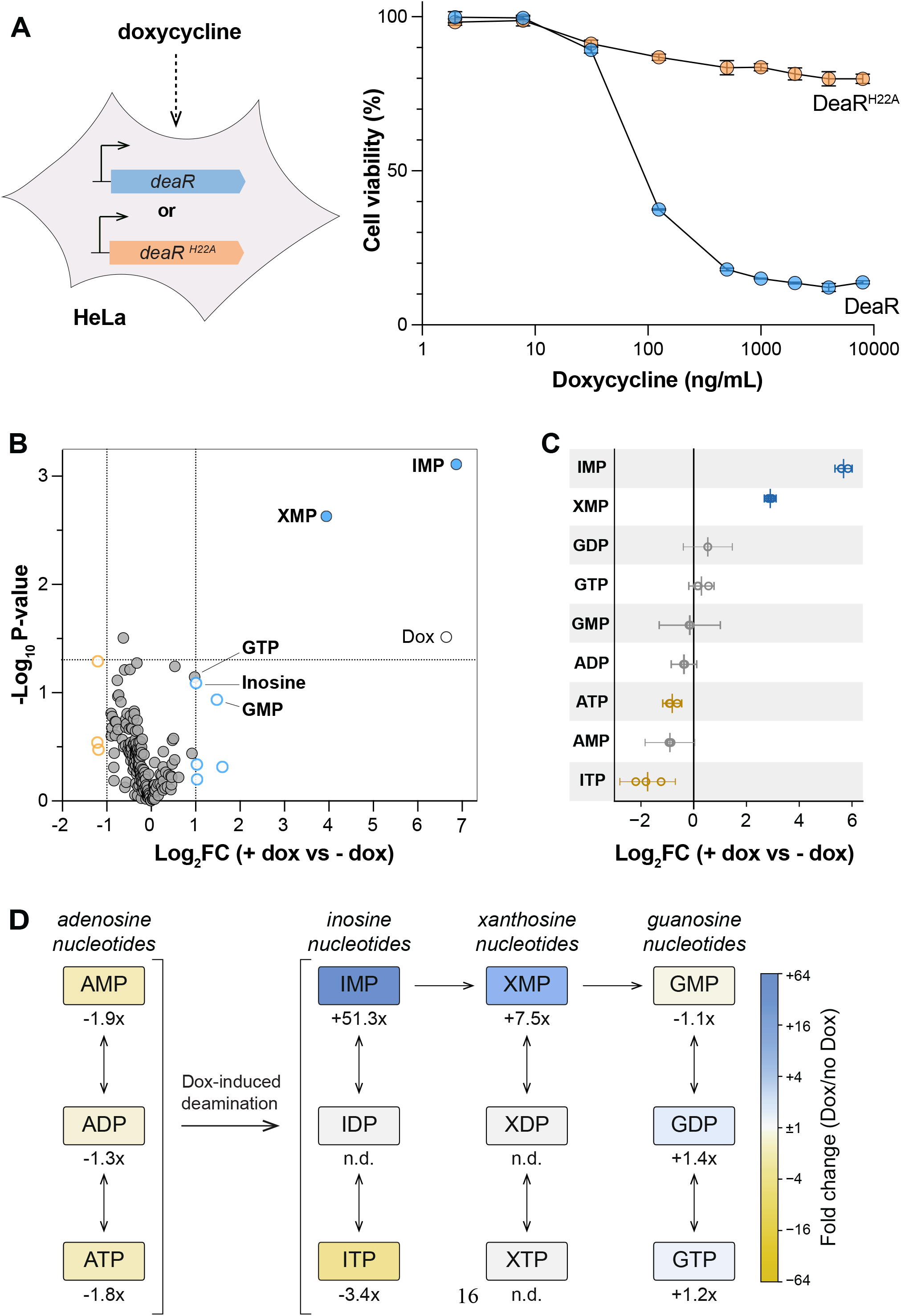
DeaR expression reduces mammalian cell viability and alters purine nucleotide metabolism. **(A) Left**: Schematic of constructs and experimental approach used to evaluate the toxicity of DeaR to HeLa cells. **Right**: Effect of DeaR or DeaR-H22A induction on HeLa cell viability as a function of doxycycline concentration. **(B)** Extent and significance of changes in metabolite amounts after DeaR induction in HeLa cells, detected via untargeted analysis. Metabolites that showed a fold change ≥2 or ≤-2 are indicated by colored circles. Metabolites that met this criterion and had a p-value <0.05 are indicated by filled, colored circles. Metabolites with fold changes near or >2 that are related to purine metabolism are labeled. Doxycycline (Dox) was also detected. **(C)** Fold changes in purine nucleotides detected in HeLa cells producing DeaR, detected by targeted metabolomic analysis. Each circle represents a biological replicate (n = 3), and squares indicate the mean. Error bars represent 95% confidence intervals (CI). Colored circles represent significant up- or downregulation. **(D)** Schematic showing the pathway relationships between nucleotide variants and pools within HeLa cells. The change in amount of each nucleotide after DeaR induction is indicated by colored shading. The proposed conversion of adenine to hypoxanthine nucleotides by deamination is indicated. n.d., not detectable.

To characterize the specific effect of DeaR expression on HeLa cell metabolism, we prepared 24-hour doxycycline-induced (i.e., DeaR-expressing) and uninduced samples for metabolomic profiling via untargeted ion-pairing LC-QTOF-MS. Samples from induced cells showed pronounced increases in the nucleotides inosine monophosphate (IMP) and xanthosine monophosphate (XMP) (**Figure 3B and file S2**)–in addition to moderate increases in GTP, inosine, and GMP–indicating that DeaR induces a shift in cellular nucleotide pools with an effect on purine metabolism in particular. To further elucidate this effect, we used pHILIC LC-MS and standards to examine targeted purine metabolites. This approach validated the results of the untargeted analysis with respect to accumulation of IMP and XMP and allowed us to assay ITP and IDP (**Figure 3C**). We did not detect IDP but observed depletion of adenosine phosphates and ITP. We note that DeaR elicited this pronounced effect on the HeLa-cell metabolome even though its expression level was relatively low: based on sequencing reads and comparison with housekeeping genes (*53–55*), we estimate DeaR abundance at 0.3 transcripts per cell (**figure S3B**). The marked accumulation of IMP and XMP, along with additional effects on related compounds, therefore underscores the potent influence of DeaR on HeLa cell physiology. These changes are consistent with DeaR deaminating adenine nucleotides in the host cytosol, redirecting the adenosine pool toward the inosine branch of purine metabolism (*56–58*) (**Figure 3D**).

### Purification of DeaR yields homomeric complexes

The two nucleotides that accumulated in DeaR-expressing HeLa cells (IMP and XMP) might point to an AMP deaminase activity for DeaR: IMP is the product of AMP deaminases, and IMPDH, which is present in HeLa cells, can theoretically convert the enriched IMP to XMP (*59*, *60*). However, because the pools of nucleoside derivatives in cells are interrelated through shared metabolic pathways and cross-regulation (*61*, *62*), the changes we observed may not directly indicate DeaR’s substrate. The mammalian enzymes ITPA and NUDT16, for example, convert ITP and IDP (which are produced by promiscuous kinases or under stressful conditions) to IDP and IMP, respectively, thereby serving as alternative sources of these compounds (*63*, *64*). To enable further characterization of the structure and activity of the DeaR toxin, we expressed N-terminally His6-tagged DeaR and its H22A mutant in *E. coli* and purified the proteins through a multi-step process beginning with Ni-NTA immobilized metal affinity chromatography. After subsequent ion-exchange chromatography, we obtained preparations with >95% purity, showing DeaR present in two peaks at different salt concentrations (**Figure 4A** and **figure S4A**). The higher-salt peak was purified further using size exclusion chromatography (SEC) (**Figure 4A**) and showed primarily high-molecular weight species and some low-molecular weight species. In comparison, the lower-salt peak was primarily low-molecular weight species. Sedimentation velocity analytical ultracentrifugation (AUC) experiments, performed on WT DeaR, indicated that the low-molecular-weight fraction contained 74% dimer and minor amounts of monomer and trimer, while the high-molecular-weight fraction contained 81% dodecamer with 13% dimer (**Figure 4B**). These results were validated by size-exclusion chromatography with multi-angle light scattering (**figure S4C**). Moreover, a thermal shift assay performed on the lower- and higher-molecular weight peaks yielded distinct melting temperatures of 47.6°C and 60.9°C, respectively, consistent with greater stability of the dodecamer (**Figure 4C**). The similar peaks formed by WT DeaR and its H22A mutant in SEC indicate that the mutant exhibits similar oligomerization behavior.

**Figure 4.**
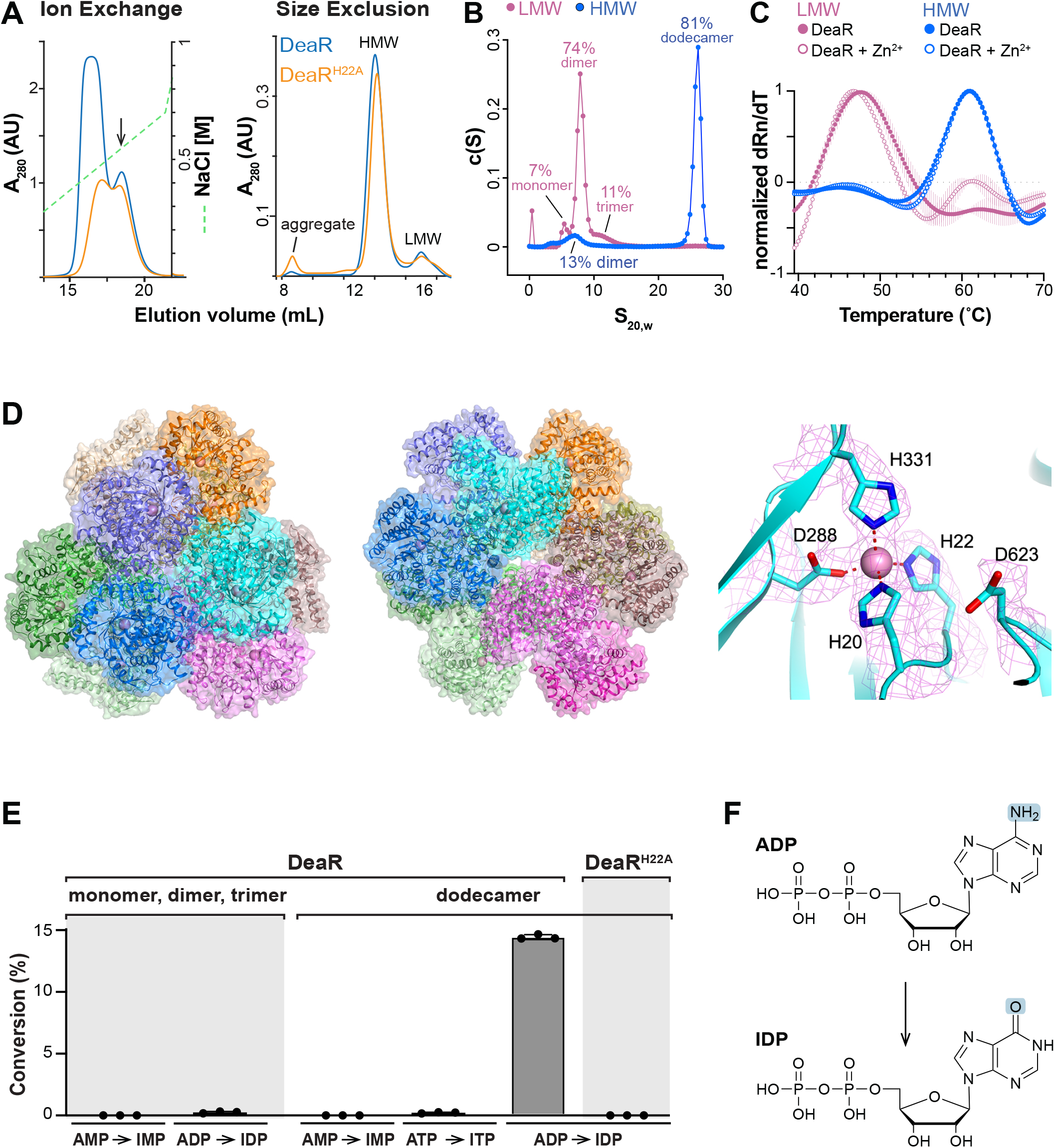
DeaR forms a dodecamer with ADP deaminase activity. **(A) Left:** Elution profile of DeaR and the H22A mutant prepared from an *E. coli* expression strain and separated by ion-exchange chromatography. The arrow indicates the fraction that was sampled and used for size exclusion chromatography. The green dotted line indicates concentration along the NaCl gradient. **Right:** Elution profiles of the higher-charge fractions of WT and H22A DeaR (obtained from ion exchange chromatography) separated by size exclusion chromatography. **(B)** Results of analytical ultracentrifugation analysis. **(C)** Results of thermal shift assay. **(D)** Cryo-EM structure of DeaR, viewed down the three-fold (**left**) and two-fold (**center**) symmetry axes of the dodecamer. Each monomer is given a different color, with monomers in the same trimer sharing similar colors. The zinc ion is shown as a sphere (pink), indicating the locations of the twelve active sites. The molecular surface is shown with semi transparency. The active site of DeaR is shown at **right**. Magenta mesh represents cryo-EM density. **(E)** Percent conversion of the nucleotides AMP, ADP, and ATP to inosine nucleotides by the DeaR dodecamer and LMW fractions obtained in (A,B). The HMW fraction of the DeaR H22A mutant did not show activity. **(G)** Deaminase reaction catalyzed by DeaR. HMW, high molecular weight; LMW, low molecular weight.

### Structure of DeaR reveals a dodecamer with tetrahedral symmetry

We were intrigued by indications that purified DeaR forms a dodecamer; because DeaR is already a large bacterial protein at ∼80 kDa, such a complex would approach one megadalton in size and the order of magnitude of the largest protein complexes found in bacteria, such as the ∼2.5-MDa 70S ribosome and the ∼1.8-7.5 MDa encapsulin (*65*, *66*). To elucidate the structure and oligomerization state of DeaR, we prepared our dodecamer-enriched SEC fraction (**Figure 4A**) for single-particle analysis using cryo-electron microscopy (cryo-EM), and determined the structure at 2.3 Å resolution (**figure S5, table S2**). The structure reveals a dodecamer with tetrahedral symmetry, with four trimers located at the corners of a tetrahedron (**Figure 4D**). The three monomers of each trimer have close contacts at one end, but the three TIM barrel domains are splayed apart. Three segments of each monomer in the trimer interface are disordered, residues 1-13, 496-573, 681-713. Some of these residues form a β-sheet that packs against another monomer of the trimer (**figure S6A**). There are two large cavities, one at the center of the trimer and the other in the dodecamer.

The active site of DeaR is located at the C-terminal end of the TIM barrel, where a zinc ion is coordinated by His20, His22, Asp288, and His331 (**Figure 4D**). This coordination is distinct from that in other adenosine deaminases and the AF3-modeled DeaR monomer (**Figure 2A, 2C**), although His22, the residue mutated in the predation-susceptible H22A mutant (**Figure 2E**), is confirmed as a primary metal ligand. A more detailed comparison suggests that DeaR is a distinct type of deaminase, with the ligands to the metal ion coming from different regions of the TIM barrel (**figure S6B, S6C**). The zinc ion is coordinated tetrahedrally by its four ligands in the current structure of DeaR (**Figure 4D**), while it is coordinated by five ligands arranged in a trigonal bipyramid in mouse adenosine deaminase (PDB 1A4M) (*67*), with one of the ligands being a hydroxyl from a substrate-analog inhibitor.

### DeaR catalyzes ADP deamination

While cryo-EM analysis indicated the presence of a metal ion in the DeaR active site, this approach cannot unequivocally determine the identity of the metal. The results of our Foldseek and AF3 analyses, combined with the metabolic shifts we observed in HeLa cells induced to make DeaR, strongly suggested that DeaR is a deaminase that acts on nucleotides (*35*). Zn^2+^-dependence is a common feature across major families of deaminases (*36*). We performed ICP-MS and indeed found that DeaR exhibits near-stoichiometric Zn^2+^-loading (**figure S4B**). Zn^2+^ occupancy was significantly lower for the H22A mutant, consistent with a role for this residue in metal binding (**figure S4B**).

Based on the accumulation of IMP and XMP in DeaR-producing HeLa cells, we hypothesized that DeaR acts on one or more adenosine phosphates to produce inosine phosphates. We therefore tested purified DeaR and its H22A mutant for deamination of ATP, ADP, and AMP using LC-MS. We observed productive substrate conversion in the dodecameric fraction of WT DeaR but not the low-molecular-weight fraction or the high-molecular-weight fraction of the H22A mutant (**Figure 4E, 4F**). DeaR deaminase activity was specific to ADP and, consistent with the role of His-22 in metal binding, required this residue. We attempted to determine the structure of DeaR in complex with its substrate by including 10 mM ADP or IDP in the cryo-EM sample, but did not observe any EM density for the nucleotide.

### Diverse toxin families are associated with bacterial *reb* clusters

Examining DeaR homologs in bacteria, we found that most were encoded by strains that also contained *rebP1* homologs, suggesting a role for R-bodies in delivering these deaminase toxins (**Figure 2D and figure S2**). However, *rebP1* homologs are found in a much broader range of bacterial genomes than those encoding DeaR-like proteins, prompting us to investigate whether R-bodies might also deliver other classes of toxins (*9*). We identified bacterial strains encoding RebP1 homologs (BLASTP, E ≤ 10⁻¹⁰) and examined a 20-kb chromosomal region centered on each homolog for genes encoding putative toxins and effectors **(figure S7)**. These include nucleic acid- and protein-modifying enzymes (deaminases, a CheD-like deamidase, an HNH endonuclease, and a nucleotidyltransferase), cell wall hydrolases (muramidases, a GH24 lysozyme, and an NlpC/P60 amidase), and holins. We also identified components of extracellular contractile injection systems (eCISs) and defense or addiction modules, several of which have been implicated in bacterial toxicity during characterized interspecies interactions: the *Photorhabdus* virulence cassette (PVC), an eCIS that delivers toxins into insects such as the wax moth (*68–70*); *Neisseria* and *Bordetella* transglycosylases that produce peptidoglycan toxic to ciliated human cells (*71–73*); and vgrGs, part of the *Vibrio cholerae* T6SS machinery, which delivers cytoskeleton-modifying toxins into *D. discoideum* and macrophages (*74*). The widespread co-localization of putative toxin genes with *rebP1* homologs across phylogenetically diverse bacteria (**Figures 5A and 5B**) suggests that such <u>c</u>ytotoxic <u>e</u>ffectors <u>d</u>elivered by <u>R</u>-bodies (CEDRs) may represent a broadly conserved mechanism of interaction between bacteria and other organisms.

**Figure 5.**
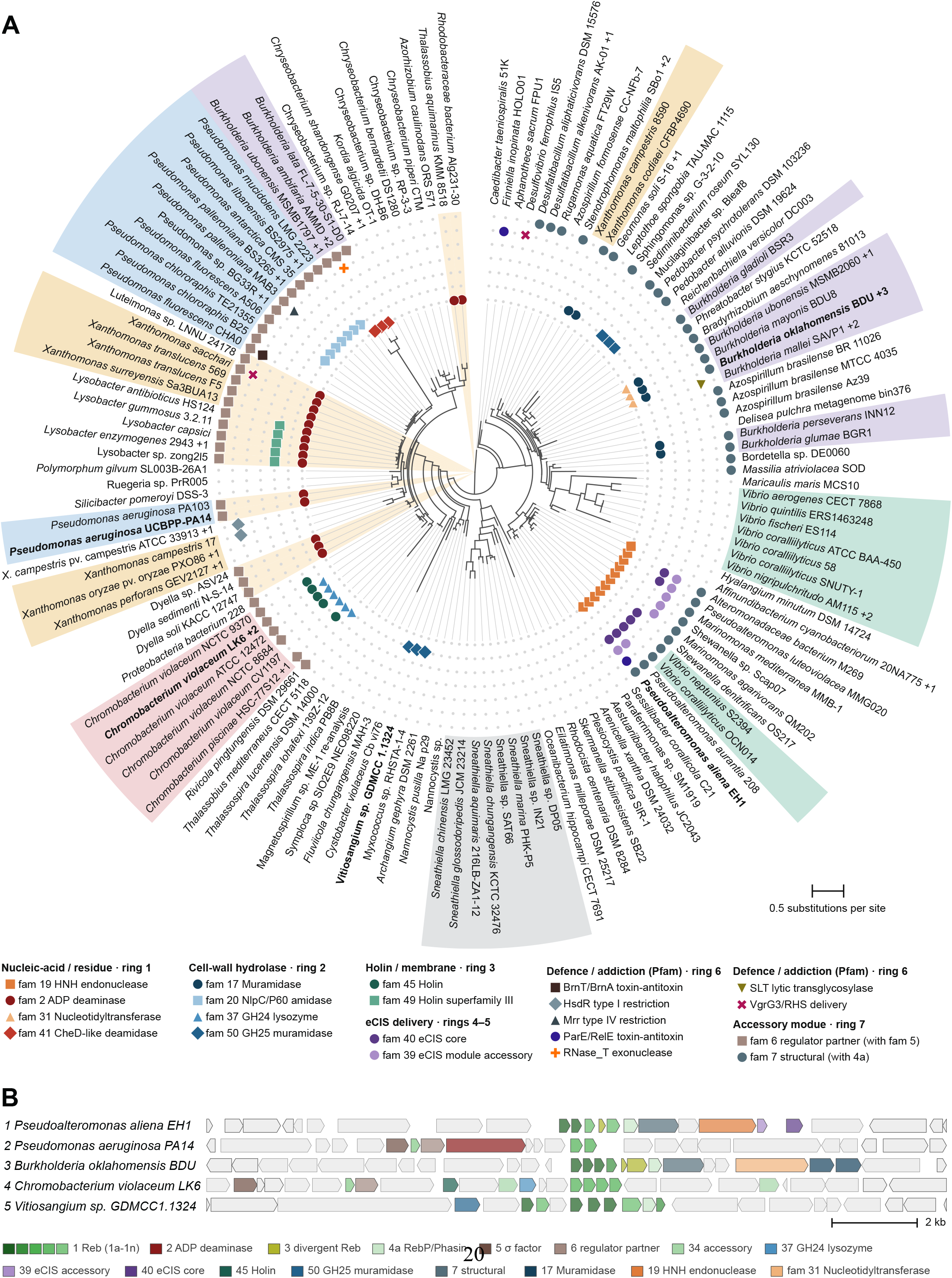
Genes for diverse toxins are associated with bacterial *reb* clusters. **(A)** Phylogenetic tree showing relatedness of bacterial strains containing *rebP1* homologs. Strain names are color-coded if they belong to the most-represented genera. The tree branches of strains that contain ADP deaminases homologous to DeaR are indicated with a maroon dot and shaded beige. The presence of genes for other toxins are denoted by symbols on each tree branch as indicated by the figure legend. **(B)** Chromosomal arrangement of putative toxin genes in selected genomes, indicated in bold on the tree in (A). The genes are shaded according to their putative function as indicated in the legend.

## Discussion

A role for R-bodies in host toxicity was first proposed almost 70 years ago, when, without knowledge of their ultrastructure or dynamics, Tracy Sonneborn suggested that they deliver toxins (*75*). Two key observations–that R-bodies turn into long needles in response to changes in environmental conditions, such as pH (*11*, *19*) and that they are insufficient to confer toxicity in other bacteria (*14*)–further supported an intracellular toxin delivery mechanism (**Figure 1D**). However, this possibility remained unexamined for decades. Having discovered that R-bodies are produced by *P. aeruginosa* strain PA14 (*9*), we set out to test the R-body toxin-delivery model in this globally important pathogen.

In the present work, we observed that PA14 R-bodies are produced during co-culture with the model phagocyte *D. discoideum* and that they extend after R-body-containing bacteria are engulfed (**Figure 1F, 1E**). We also identified a putative toxin gene, *PA14_27675*, in the R-body-encoding region of the PA14 genome. We demonstrated that *PA14_27675* codes for an ADP deaminase that forms a large, subcellular structure that is associated with the R-body and toxic to eukaryotic cells (**Figures 2G, 2H**, **3A**, and **4E** and **F**). HeLa cells exposed to PA14_27675, which we have named DeaR, contain altered pools of nucleoside derivatives (**Figure 3B and C**) consistent with deamination of one or more adenosine phosphates and detoxification of inosine tri- or diphosphate (**Figure 3D**). The toxicity of DeaR to mammalian cells could be attributed to its depletion of nucleotide pools crucial for nucleic acid synthesis, signaling, and the function of motor and transport proteins. Another possibility is that, because DeaR activity promotes the accumulation of inosine derivatives, toxicity arises from substitution of these compounds for those natively incorporated into RNA or those that modulate transcription factor activity. We favor the hypothesis that DeaR toxicity is primarily caused by accumulation of harmful IDP, which is detoxified to IMP. Consistent with this, we detected accumulation of IMP, but not IDP or ITP, during heterologous DeaR expression (*76–79*) (**Figure 3C,D**).

To our knowledge, DeaR is the first characterized deaminase that acts specifically on ADP. Purified DeaR showed activity solely with ADP and converted 14.3% of the provided substrate. DeaR efficiency may depend on an unidentified cellular cofactor (*80*). Nevertheless, the results of our cryo-EM analyses and biochemical characterization together support an assembly-gated catalytic mechanism in which only the higher-order DeaR oligomer brings the catalytic residues into a productive Zn²⁺-coordinated geometry (*36*, *81*) (**Figure 4D**). Cryo-EM and analytical ultracentrifugation (AUC) analyses revealed that DeaR forms a 966-kDa dodecamer complex (**Figure 4B and D**). Imaging in PA14, however, suggests a much larger complex, ranging from 200 to 700 nm in length compared to an approximately 16-nm diameter of the dodecamer (**Figure 2G and H**). Further oligomerization of DeaR within PA14 may serve to reversibly inactivate the protein and protect the producing bacterium from DeaR-mediated toxicity.

Looking at the distribution of DeaR across bacterial phylogeny, we identified homologs in dozens of strains that also contain R-body structural genes (**Figure 2D**). Many of these bacteria are found associated with plants and/or soil, indicating that the PA14 model of R-body-dependent toxicity is common in these niches. Examining bacteria with R-body structural genes more broadly, we identified several additional types of toxins encoded near these genes, indicating distinct mechanisms for combating phagocytic predation and paralleling the variety of strategies used by bacteria in defense against viruses (phages) (**Figure 5A**) (*82*, *83*). The diversity of R-body-proximal candidate toxin genes (i.e., candidate CEDRs) suggests that R-bodies can serve as delivery vehicles for proteins that disable hosts through distinct mechanisms. We note that because our search for candidate toxin genes was limited to chromosomal regions surrounding R-body structural genes, the toxin families in **Figure 5A** are a conservative representation of potential R-body-associated toxins; co-regulation may also allow R-bodies to deliver products of distal genes that contribute to host toxicity (*84*, *85*). Moreover, the niches occupied by these R-body- and toxin-producing bacteria range from freshwater environments to sites on and within marine and terrestrial animals, indicating that R-bodies have widespread potential to modulate ecosystem dynamics and pathogenicity.

Our findings have implications for the relevance of R-bodies in various interkingdom interactions, including human infections, where R-body-dependent toxin delivery might disable macrophages that would otherwise consume multiple bacteria in a population (*86*). Modulating the process of R-body-mediated toxicity may facilitate treatment of *P. aeruginosa* infections. Moreover, the observation that R-body genes are proximal to genes for diverse putative toxins indicates that R-body activity remains robust in the presence of various “cargoes” and might therefore be co-opted in applications delivering small-molecule-based or gene-, RNA-, or protein-editing therapies (*87*).

## Acknowledgments

We thank Jegg Yoon for their artistic contributions, Maurice Chakour for his experimental contributions, and Christian Sieben and Nicolas Oswaldo Gomez for helpful discussions. During the preparation of this work, the authors used Claude Opus 5.0 and Gemini Pro 3.1 for data analysis. After using these tools and services, the authors reviewed and edited the content as needed and take full responsibility for the content of the manuscript.

We acknowledge the use of facilities and instrumentation supported by NSF through the Columbia University Nano Initiative, and the Materials Research Science and Engineering Center DMR-2011738 under the leadership of Keren Bergman.

We thank the Confocal and Specialized Microscopy Shared Resource at the Herbert Irving Comprehensive Cancer Center at Columbia University under the leadership of Theresa Swayne; Stem Cell Initiative Flow Cytometry Core Facility at Columbia University Irving Medical Center, under the leadership of Michael Kissner; METALab (Multi-Element Trace Analysis Laboratory) at Columbia University Irving Medical Center of Environmental Health Sciences under the leadership of Kathrin Schilling for the analysis of metal content with ICP-MS; Johnson Foundation Structural Biology and Biophysics Core at the Perelman School of Medicine under the leadership of Kushol Gupta; BRCF Metabolomics Core Facility (RRID:SCR_026721) at the University of Michigan Medical School under the leadership of Maureen Kachman; and Proteomics and Metabolomics Core Facility (RRID:SCR_027811) at Memorial Sloan Kettering Cancer Center under the leadership of Mara Monetti.

## Funding

Hypothesis Fund (LEPD).

National Institutes of Health grant (LEPD) R01AI189733.

National Institutes of Health grant (LT) R35GM118093.

National Institutes of Health grant (Confocal and Specialized Microscopy Shared Resource of the Herbert Irving Comprehensive Cancer Center at Columbia University) P30CA013696.

NSF GRFP grant (DKR) DGE 2437839. Any opinions, findings, and conclusions or recommendations expressed in this material are those of the authors and do not necessarily reflect the views of the National Science Foundation.

National Center for CryoEM Access and Training (NCCAT) and the Simons Electron Microscopy Center located at the New York Structural Biology Center:

The NIH Common Fund Transformative High Resolution Cryo-Electron Microscopy program (U24 GM129539 and R24 GM154192).

The Simons Foundation (SF349247).

The NY State Assembly.

## Author contributions

Conceptualization: DTKR, KG, TYT, APW, LEPD

Methodology: DTKR, KG, TYT, JP, HL, TW, TMY, MS, LEPD

Investigation: DTKR, KG, TYT, JP, HL, TW, TMY, MS, LEPD

Visualization: DTKR, KG, TYT, APW, LT, LEPD

Funding acquisition: APW, VWC, LT, LEPD

Project administration: APW, VWC, LT, LEPD

Supervision: VWC, LT, LEPD

Writing – original draft: DTKR, KG, TYT, APW, LEPD

Writing – review & editing: DTKR, KG, TYT, JP, HL, TW, TMY, MS, APW, VWC, LT, LEPD

## Data and materials availability

The genomes used for phylogenetic analyses were obtained from the JGI/IMG database and supplemented with genomes from NCBI. The cryo-EM structure was deposited into the PDB with ID 39OY and is undergoing evaluation. Protein structures used for structural and sequence comparison were acquired from the PDB (1UIP, 6IJN, 8HU6, and 2A3L). All other data needed to evaluate the conclusions in the manuscript are present in the manuscript and/or the Supplementary Materials. All strains created for use in this study are available upon reasonable request.

## Supplementary Materials

**Materials and Methods**

**Figs. S1 to S7**

**Tables S1 and S2**

**References** (**88–112**)

**Data S1 to S3**

**Movie S1**

