## Supplementary material for "*Pseudomonas aeruginosa* R-bodies deliver an ADP deaminase toxin": Materials and Methods

Bacterial strains and growth conditions

Bacterial strains used in this study are listed in **table S1**. *Escherichia coli*, *Pseudomonas aeruginosa*, and *Klebsiella pneumoniae* strains were routinely grown at 37°C, unless otherwise noted, in lysogeny broth (LB). Liquid cultures were shaken at 250 rpm. LB plates contained 1.5% agar.

Creation of *P. aeruginosa* RcgA+ and $P_{rha}$-*rcgA*

All plasmids used and created in this study are listed in **data S3**; primer sequences are available upon request. To create the RcgA+ strain, WT *rcgA* (*PA14_27700*) was amplified from the PA14 genome and inserted into pAKN69 (*89*) via restriction ligation. To create $P_{rha}$-*rcgA*, a rhamnose-inducible promoter was amplified from pLD4358 (*90*) and used to replace the $P_{PA1/04/03}$ promoter of pAKN69-*rcgA* using restriction ligation. The resulting plasmid was transformed into UQ950 for amplification, then moved into PA14 using a triparental conjugation via the donor strain BW29427 and the helper strain *E. coli* β2155 containing pUXβF-13 (*91*). Successful integration was selected for on LB plates supplemented with 70 µg/mL gentamicin.

R-body purification

The *P. aeruginosa* $P_{rha}$-*rcgA* strain was grown for 14-16 h in LB supplemented with 30 µg/mL gentamicin. These precultures were diluted 1:100 into 50 mL fresh LB with 30 µg/mL gentamicin and grown to an optical density at 500 nm ($OD_{500}$) of 0.6-0.8. Cultures were placed at 4°C for 30 minutes, then R-body production was induced by adding L-rhamnose to 4 mM and cultures were incubated at 25°C with shaking for 20 hours. Cells were pelleted via centrifugation at 4696 x g for 1 hour and the supernatant was removed. Cell pellets were frozen, then thawed in a 9 mL lysis buffer (PBS, pH 7, 1.5 mg/mL lysozyme). Once thawed, lysates were treated with 250 U Pierce Universal Nuclease for Cell Lysis (ThermoFisher), on ice for 1 hour. Lysates were then treated with sodium dodecyl sulfate at a final concentration of 1% and nutated at 4°C overnight. R-bodies were pelleted via centrifugation at 4696 x g for 1 hour and the supernatant, containing all SDS-soluble components, was removed. R-body pellets were then washed 3X by resuspension in PBS (pH 7) and pelleted as above. Purified R-bodies were stored as pellets at 4 °C.

Sample preparation for SEM

Purified R-bodies were suspended in 1 mL neutral buffer (PBS pH 7.0) or acidic buffer (0.1 M sodium acetate pH 5.0), pelleted at 16,873 × g for 1 minute, then resuspended again to ensure complete buffer exchange. R-bodies were again pelleted, then resuspended in 1mL pre-fixative solution (2% paraformaldehyde, 2.5% glutaraldehyde, 0.0075% L-lysine in either PBS at pH 7 or 0.1 M sodium acetate at pH 5) and nutated at 21°C for 30 min in the dark. R-bodies were pelleted, then washed in 1 mL sterile buffer and fixed in 2.5% glutaraldehyde in PBS at pH 7 or 0.1 M sodium acetate at pH 5 at 21°C for 30 min in the dark. Fixed pellets were washed twice in buffer and dehydrated through a series of 1 mL ethanol dilutions (25%, 50%, 75%, 95%, 2 × 100% ethanol) in buffer. R-bodies were then resuspended in 20 µL 100% ethanol, loaded onto silicon glass chips, and dried in a vacuum desiccator over desiccant beads. Chips were adhered using PELCO Tabs (Ted Pella) onto 12.7 x 6mm, grooved-edge aluminum pins. Samples were then sputter-coated 2X for 20s with Au/Pd and imaged using a Zeiss Sigma VP SEM.

R-body extension assay

R-body pellets were resuspended in 1 mL citrate phosphate buffer (0.1 M sodium citrate, 0.2 M $Na_2HPO_4$ pH 5.0-6.0), then 3x 300 µL were transferred to a flat-bottom, 96-well clear plate (Greiner). Plates were covered and kept in the dark for 4 hours to allow R-bodies to sediment. The uppermost 200 µL of solution/suspension were removed and transferred to empty wells of the 96-well plate. OD at 595nm was measured using an Agilent BioTek Synergy H1 Microplate Reader.

*Dictyostelium discoideum* fruiting body propagation

Five milliliters of LB were inoculated using a frozen stock of *Klebsiella pneumoniae* DBS0351098 and incubated overnight at 37 °C with shaking at 250 rpm. Two hundred microliters of this overnight culture were spread with glass beads onto a dry SM agar plate (1% glucose, 1% proteose peptone, 0.1% yeast extract, 4.15 mM $MgSO_4$, 13.96 mM $KH_2PO_4$, 3.45 mM $K_2HPO_4$, 2% agar). The plate was incubated for 2 hours at 37 °C, then cooled to 4 °C for 10 minutes. The plate was allowed to return to room temperature, and a frozen stock of *Dictyostelium* was spread onto the middle using a wooden applicator stick. The plate was incubated at 23 °C; after 5 days, fruiting bodies began to appear in the *Dictyostelium* clearing.

*D. discoideum* live imaging

*D. discoideum* cells from the feeding front of a fruiting-body plate were used to inoculate 4mL HL5 medium (Formedium) supplemented with 300 µg/mL carbenicillin and 300 µg/mL streptomycin and incubated as standing liquid cultures for 2 days at 23°C. The overlying medium was gently removed using a micropipette, and cells were washed once and resuspended in 1 mL of Sörensen buffer ("SB", 14.7 mM $KH_2PO_4$, 2.04 mM $Na_2HPO_4$). *P. aeruginosa* RcgA+ and *K. pneumoniae* cultures were grown in LB at 25°C for 14-16 hours, then 40 µL and 60 µL of each respective culture were combined in a microcentrifuge tube. Ten microliters of this suspension were spotted, then spread with a micropipette tip into a Thermo Scientific Nunc Lab-Tek II Chambered Coverglass and allowed to dry for 20 minutes. One hundred microliters of the *D. discoideum* suspension were added to each well, followed by an additional 100 µL of fresh SB. After 20 min of incubation at 25°C, the chambered coverglass was transferred to an LSM880 confocal microscope (Zeiss) for DIC imaging.

*D. discoideum* plate imaging and plate killing assay

*D. discoideum* was grown with *Klebsiella pneumoniae* DBS0351098 and various *P. aeruginosa* strains to assess the effect of co-incubation on gene expression (using fluorescent strains) and the roles of *reb* cluster genes in predation resistance (using deletion mutants).

Thirty microliters each of PA14 and *K. pneumoniae* overnight cultures were diluted into 3 or 5 mL fresh LB, respectively. Subcultures were incubated at 37° C with shaking for approximately 2.5 hours until reaching an OD between 0.4 and 0.6 for PA14 and 0.7 and 0.9 for *K. pneumoniae*. Cultures were combined in an OD ratio of 3.5 : 96.5 PA14 to *K. pneumoniae* then diluted to a final OD of 0.57 in 200 µL (total) with sterile LB. Two *D. discoideum* fruiting bodies (which are estimated to contain 100,000 spores each (*92*) were suspended in development buffer (5 mM $Na_2HPO_4$, 5 mM $KH_2PO_4$, 1 mM $CaCl_2$, 2 mM $MgCl_2$ pH 6.5) to an approximate concentration of 200,000 spores/mL. Two thousand spores were added to the PA14-*Klebsiella* mixture, then spread onto dried SM plates using glass beads. Plates were incubated at 23°C for 10 days and imaged daily after day 3 with an Epson Expression 11000XL scanner. Fluorescently scanned plates were imaged on the 10th day of growth using a GE Typhoon FLA 7000 fluorescence imager. Images for plaque size quantification were obtained daily on incubation days 3-10, using an Epson Expression 11000XL scanner.

For plaque identification and analysis, whole-plate scans were subdivided into a 5 × 3 grid of non-overlapping sub-images using pixel-coordinate cropping. To correct for positional drift between days, each condition was aligned to day 5 using the Oriented FAST and Rotated BRIEF (ORB) feature detector in OpenCV. Plate images were then padded with a black border (15% of image width on each side) to preserve edge features during transformation. To prepare for rotational alignment, keypoints were computed for the reference and target images and matched using a brute-force Hamming distance matcher, retaining the top 25% of matches by distance. A homography transformation matrix was estimated using RANSAC and applied to warp each target image to align with the reference frame. In cases where ORB failed to yield sufficient keypoints (fewer than 10), alignment defaulted to phase correlation, which estimates a translational shift between images; shifts exceeding one-third of the image dimension in either axis were rejected, and the unaligned image was retained. Aligned grayscale images were binarized using a fixed intensity threshold applied independently per day (optimal binarization thresholds were determined visually). Pixels with intensity values above the threshold were classified as foreground, and the resulting binary image was inverted to yield white plaques on a black background. Binary images were processed prior to plaque detection by removing small noise objects (area < 1,200 pixels) and applying morphological closing with a 10 × 10 elliptical structuring element to fill gaps within plaque interiors resulting from variable contrast between empty agar and *D. discoideum* fruiting bodies. Plaque candidates were identified by contour detection (OpenCV findContours, RETR_LIST mode). Contours were retained as plaques if they satisfied both an area criterion (1200–80,000 pixels) and a circularity threshold ($4\pi \times \text{area} / \text{perimeter}^2 > 0.05$). Detected plaques were tracked across consecutive time points using a nearestneighbor greedy matching algorithm. Each plaque identified on day 5 was initialized as a track. For each subsequent day, active tracks were matched to the nearest detected plaque centroid (Euclidean distance < 200 pixels) and results were inspected visually. Only tracks with detections at every timepoint throughout the full observation period (days 5–11) were classified as surviving plaques and retained for downstream analysis. For each surviving track, the pixel area at each timepoint was recorded. Growth trajectories were plotted as plaque area versus growth day using MATLAB R2026a.

AlphaFold3 structural prediction

Structural predictions were generated locally using AlphaFold 3 (version 3.1.1) with DeepMind model parameters on NVIDIA A100 graphics processing units (GPUs). Precomputed multiple sequence alignments (MSAs) were supplied for each protein as unpaired MSAs, with paired MSAs left empty. UniRef100 ligands and metal ions were specified using Chemical Component Dictionary (CCD) identifiers: ADN (adenosine), AMP (adenosine monophosphate), ADP (adenosine diphosphate), ATP (adenosine triphosphate), and ZN (zinc ion). Predictions were generated using the default AF3 settings of five diffusion samples per random seed and ten recycling iterations. For each condition, 40 random seeds were specified across eight input files containing five seeds each, yielding a total of 200 structural predictions per condition. Prediction confidence was evaluated using the predicted local distance difference test (pLDDT), which measures local structural confidence; the predicted template modeling score (pTM), which measures confidence in the overall structure; the interface predicted template modeling score (ipTM), which assesses confidence in the relative positioning of interacting components; and the AF3 ranking score, a composite metric used to rank predicted structures. Confidence metrics were reported as distributions: the five diffusion samples drawn from a given seed were first reduced to a per-seed median and distributions were then reported across the 40 seeds (all protein Cα atoms (global), protein Cα atoms within 12 Å of the catalytic zinc (active site), and all heavy atoms of a given ligand (per-ligand confidence)).

DeaR homolog phylogeny

DeaR homologs were identified by BLASTP against NCBI ClusteredNR (E $\leq 10^{-5}$; maximum 1,000 hits). Partial proteins and hits covering <70% of DeaR were excluded. Family membership and homologous regions were assessed using a profile HMM derived from the curated DeaR-family alignment. Within-genus redundancy was reduced at ≥90% identity and ≥90% mutual overlap. The resulting 52 sequences were aligned using MAFFT v7.526 (L-INS-i), retaining columns with ≥70% residue occupancy. The 709-position alignment was analyzed using IQ-TREE v3.1.4 with the Q.PFAM+I+G4 model, selected by ModelFinder using BIC. Support was estimated using 1,000 ultrafast bootstrap and 1,000 SH-aLRT replicates. The tree was displayed unrooted.

Comparative genomics and phylogenetics of *reb*-cluster loci

*Genome set.* Candidate *reb*-carrying strains were identified in the JGI/IMG database (*93*) by searching for genes encoding homologs of the *C. taeniospiralis* R-body structural protein RebB (Pfam PF11747) (*94*). The search identified 229 genes, with one to eight genes per strain. Genome assemblies were downloaded from JGI, and the collection was expanded to include additional strains and species belonging to the same taxa, yielding 154 genomes. Nine candidate strains were excluded because they represented IMG-only metagenome bins (n = 6) or lacked strain-level NCBI assemblies (n = 3). An additional 11 genomes not included in the JGI download were retrieved from NCBI RefSeq/GenBank (*95*) using Entrez E-Search and E-Summary, yielding a final collection of 165 genomes representing 164 distinct strain names. Of these, 113 corresponded to the original candidates, 42 represented additional strains or species belonging to the same taxa, and nine were identified through other sources. Thus, the collection represents a selected set of *reb*-carrying genomes rather than an exhaustive survey of IMG. All genomes were annotated using Prokka v1.15.6 (*96*).

*Locus set.* RebP1 homologs were found by BLASTP (*97*) using the *Pseudomonas aeruginosa* PA14 RebP1 protein (PA14_27640) as the query ($E \leq 10^{-10}$, ≥25% sequence identity). The highest-scoring hit in each genome was retained, yielding one RebP1 homolog per genome. A 20-kb genomic region extending 10 kb upstream and 10 kb downstream of each *rebP1* homolog was extracted, yielding 165 loci. Seven genomes identified only by IMG accession numbers were assigned strain names based on their 16S rRNA gene sequences. To reduce redundancy, 28 loci were excluded: 23 from nearly identical genomes sharing gene order and ≥99.5% median amino acid identity across corresponding genes, and five from closely related strains sharing gene arrangement and 96.0–99.4% sequence identity without additional genes. This yielded 137 representative loci for visualization.

*Visualization.* Genomic loci were visualized using LoVis4u v0.2.0 *(98)*, with gene orientation standardized so that all family-1 *reb* genes were displayed from left to right. Genomic regions extending beyond contig boundaries (n = 17) were indicated by dashed bands. Phylogenetic trees were visualized using Matplotlib v3.11.1 (*99*).

<u>Construction of deletion and complementation strains and DeaR fluorescent protein fusion strain</u>
To create the *deaR* deletion, approximately 1 kb of flanking sequence from each side of *deaR* were amplified and inserted into pMQ30 via gap-repair cloning in *Saccharomyces cerevisiae* InvSc144. The resulting plasmid contained an insert of the two fused flanking regions and was transformed into *E. coli* UQ950 and verified via whole-plasmid sequencing (Plasmidsaurus). It was then moved into *P. aeruginosa* using biparental conjugation via the *E. coli* donor strain BW29427. *P. aeruginosa* single recombinants were selected on LB agar plates containing 70 µg/mL gentamicin. Double recombinants (markerless mutants) were selected on a modified LB medium (containing 10% sucrose and lacking NaCl) and genotypes were confirmed by PCR.

Strains containing more than one alteration (i.e., a reporter construct and a fluorescent protein fusion) were made by using singly altered strains as hosts for biparental conjugation. Complementation and point mutant strains were made via complementation of a full-length gene into a background strain lacking the WT copy of the gene.

*P. aeruginosa* Δ*deaR* was complemented by inserting either WT *deaR* or the sequence for the DeaR H22A variant on the chromosome. The WT DeaR complementation plasmid was constructed by amplifying the native *deaR* gene and 1kb flanking sequences on either side, which were then inserted into pMQ30. The DeaR H22A complementation plasmid was constructed by amplifying the WT copy of *deaR* and 1kb overhangs, with primers overlapping H22 to introduce the H22A mutation. Fragments were inserted into pMQ30, as above.

To create the DeaR-mScarlet GFP-RebP2 fluorescent-protein-fusion strain, 1kb of flanking sequence on either side of the 3' end of *deaR* were amplified from the PA14 genome, and *mScarlet-I* was amplified from LD3224. Fragments were joined and inserted into pMQ30 as described above.

*P. aeruginosa* confocal imaging

Single colonies of *P. aeruginosa* pAKN-*rcgA deaR-mScarlet gfp-rebP2* were used to inoculate 2 mL LB containing 30 µg/mL gentamicin and incubated for 16 hours at 25°C. Liquid cultures were subsequently diluted 10-fold, and 2 µL were mounted on a 2% agarose pad on a glass slide. Imaging was performed at 63x on a Zeiss LSM900 confocal microscope with Airyscan 2. Fiji was used for photobleach correction of live imaging trajectories and upscaling image size, for which bilinear interpolation was utilized (*100*).

For analysis of GFP/mScarlet signal spatial proximity, individual cells were segmented using Omnipose (*101*). GFP and mScarlet channels were separated and independently thresholded into binary masks using the Triangle auto-threshold method in Fiji (*102*). For every cell, the fraction of the cell positive for its signal was measured, and cells were classified as double positive when at least 2% of its area was positive for both channels. The minimum value distance between the two binary mask channels was then assessed using Fiji.

Mammalian cell line and growth conditions

HeLa CCL-2 and HEK-293T cells (ATCC) were maintained in Dulbecco's modified Eagle's medium (DMEM; Gibco, 10566024) supplemented with 10% (v/v) fetal bovine serum (FBS; Sigma-Aldrich, F0926) and 1% (v/v) penicillin–streptomycin (Gibco, 15070063) at 37 °C in a humidified atmosphere containing 5% $CO_2$. Cells were passaged at 70–80% confluency using 0.25% trypsin–EDTA (Gibco, 25200072). Cell cultures were routinely tested for mycoplasma contamination using a Mycoplasma Detection Kit (Southern Biotechnology, 103259-632).

Creation of lentivirus constructs, and plasmids for recombinant protein expression

*E. coli* DH10B was used to create constructs to express *P. aeruginosa* proteins (i) to test physiological effects on mammalian cells or (ii) for purification. To generate the lentiviral construct, we used LR Gateway Assembly (Gateway LR Clonase II Enzyme Mix, Invitrogen #11791020) to insert synthetic gene fragments (Twist Bioscience) into pINDUCER21 (Addgene #46948). Plasmids for expression and purification of DeaR and the DeaR H22A variant were designed by codon-optimizing for expression in *E. coli* and cloned via Golden Gate assembly (NEB #E1601S) into pKG001.

Construction of transfected HeLa cell lines

Lentivirus for HeLa cell line construction was produced in HEK293FT cells (ThermoFisher) using a second-generation lentiviral packaging system. Two days prior to transfection, 1 million HEK293FT cells were seeded per 6 $cm^2$ tissue culture dish in 4.5 mL of DMEM containing 10% FBS, glutamax, and penicillin/streptomycin. One day prior to transfection, the medium was replaced with DMEM containing 20% FBS. On the day of transfection, for each dish 2 µg of transfer plasmid, 2 µg of psPAX2 (Addgene #12260), 1 µg of pMD2.G (Addgene #12259), and 15 µL of TransIT-LT1 transfection reagent (Mirus Bio) was prepared in 600 µL Opti-MEM, mixed gently by pipetting, and incubated for 30 min before being added dropwise to the cells. After 8 hours, the medium was replaced with 3 mL of DMEM containing 20% FBS. Virus-containing medium was harvested at 24 and 48 hours post-transfection, pooled, clarified by centrifugation (500 × g for 5 min at 4 °C), concentrated using Lenti-X concentrator (Takara Bio), and stored at -80 °C until use.

For HeLa cell transduction, cells were grown in 6 well plates containing 2 mL of complete DMEM medium. Once the cells reached 40-60% confluence, medium was removed and replaced with 1.5 mL of complete DMEM containing lentiviral particles and 5 µg/mL polybrene. The medium was replaced after 24 hours. $eGFP^+$cells were FACS sorted using a Sony MA900 Cell Sorter to generate homogenous and stable HeLa lines expressing DeaR or the DeaR H22A variant.

Cell viability assay

HeLa CCL-2 cells transduced with DeaR or the DeaR H22A variant were plated at 10,000 cells per well in a 96-well TC-treated plate and induced with doxycycline hyclate at the indicated concentrations. Cell viability was assessed using the CellTiter-Glo 2.0 Assay (Promega) at 48-hours post-induction with doxycycline.

Preparation of samples for untargeted LC-MS/MS metabolomics and Quantitative pHILIC LC-MS metabolomics

HeLa CCL-2 cells expressing DeaR or the DeaR H22A variant were grown in 10 $cm^2$ dishes. At 60-70% confluency, they were treated with 1 µg/mL doxycycline hyclate and incubated for an additional 24 hours. The plates were then moved to ice and cells were washed with 5 mL each of chilled solutions of DPBS, then 150 mM ammonium acetate, and then quenched by pouring liquid nitrogen to cover the plate bottom. All washing and quenching was completed within 30s. Samples were wrapped in aluminum foil and stored at -80°C.

For extraction, cell plates were removed from -80 °C storage and maintained on dry ice while samples were processed individually. Each plate was transferred to a bed of wet ice, and 500 µL of 1:1:1:1 methanol:acetone:acetonitrile:water was added. Cells were scraped into a corner of the plate and quantitatively transferred to a microtube. Samples were incubated on wet ice for 10 min, then centrifuged at 4° C, 16,873 × g for 10 mins. Two hundred microliters of supernatant were transferred to an autosampler vial fully dried in a nitrogen drier. Samples were reconstituted with 100 µL of (8:2 v/v) water:methanol. A combined-sample control for drift correction was analyzed alongside individual samples.

Untargeted LC-MS/MS metabolomics

Reversed Phase LC-MS analysis was performed as previously described (*103*). Briefly, samples were analyzed on an Agilent 1290 Infinity II / 6545 qTOF MS system using the Acquity HSS T3 1.8 µm x 50 mm column (Waters Corporation, Milford, MA) and matched precolumn. Each sample was analyzed in positive and negative ion modes. The mobile phases were (A) 100% water, (B) 100% methanol, and (C) 2.5% formic acid. Metabolites were separated with a gradient from 100% A to 100% B in 10 minutes, holding at 100% B from 10-20 minutes. Mobile phase C was also ramped from 4% to 1% over the course of the run.

Central carbon metabolite detection was also performed as previously described. Briefly, samples were analyzed on an Agilent 1290 Infinity II / 6545 qTOF MS system (Agilent Technologies, Santa Clara, CA) using an Agilent ZORBAX RRHD Extend C18, 2.1 × 150 mm, 1.8 µm column with an Agilent ZORBAX SB-C8 guard column. The mobile phases were (A) 97:3 water:methanol and (B) 100% methanol; both A and B contained tributylamine and glacial acetic acid at concentrations of 10 mM and 15 mM, respectively. Metabolites were separated using a 30 minute gradient, and the column was back-flushed with mobile phase C (100% acetonitrile, no additives) between injections.

Data analysis was conducted using a hybrid targeted/non-targeted approach. Semi-quantitative data for known compounds was obtained by manual integration using Profinder v8.00 (Agilent Technologies, Santa Clara, CA.) Metabolites were identified by matching the retention time (+/- 0.1 min), mass (+/- 10 ppm) and isotope profile (peak height and spacing) to authentic standards. Non-targeted data analysis was performed using Agilent's MassHunter Find by Molecular Feature workflow (v7.0) with recursion using Agilent's Mass Profiler Pro (v8.0). A combined feature set was generated by merging untargeted features and named metabolites into a single feature list, and data were normalized using the SERRF (*104*) algorithm. The combined feature set underwent data reduction using Binner. Briefly, Binner first performs retention time-based binning, followed by clustering of features by Pearson's correlation coefficient, and the assignment of isotopes, adducts or in-source fragments by searching for known mass differences between highly correlated features. After Binner data reduction, in-house software was used to search Refmet (https://www.metabolomicsworkbench.org/databases/refmet/index.php) to provide MSI Level III identifications, or an in-house library of authentic standards was used to provide MSI Level I identifications.

Quantitative pHILIC LC-MS metabolomics

Nucleotides were analyzed using a Vanquish HPLC system (Thermo Fisher Scientific, MA, USA) coupled to a Q Exactive Plus Orbitrap mass spectrometer equipped with a HESI II probe (Thermo Fisher Scientific).

Chromatographic separation was achieved on an Atlantis Premier BEH Z-HILIC column (100 mm × 2.1 mm, 2.5 µm particle size; Waters). Mobile phase A consisted of 10 mM ammonium carbonate (Sigma Aldrich, CAS 506-87-6) in water, and mobile phase B consisted of acetonitrile:water (95:5). All solvents were LC–MS grade (Fisher Chemical, Thermo Scientific, CAS 75-05-8). The linear gradient was set as follows: 0 min, 95% B; 15 min, 50% B; 15.5 min, 30% B; 17.5 min, 30% B; 17.6 min, 95% B; 25 min, 95% B. The flow rate was set to 200 µL $min^{-1}$, the column temperature was maintained at 27 °C, and the autosampler was held at 4 °C. A volume of 5 µL was injected for each analysis.

The ionization source was operated with the following parameters: sheath gas, 50; auxiliary gas, 10; sweep gas, 1 (arbitrary units); spray voltage, 3.5 kV; capillary temperature, 315 °C; S-lens RF level, 55; and auxiliary gas heater temperature, 350 °C. Data were acquired in full-scan, polarity-switching mode over a mass range of 70–700 m/z, with a resolution of 70,000, an AGC target of $1 \times 10^6$, and a maximum injection time of 100 ms.
Spectra were processed using FreeStyle (Thermo Fisher Scientific) and Skyline-daily (v.26.1) with a mass tolerance of 5 ppm, using an in-house retention time library of chemical standards for metabolite identification.

RT-qPCR and RNA sequencing

The HeLa CCL-2 lines expressing DeaR or DeaR H22A and the untransfected control line were grown in 6-well plates to 60-70% confluency, then treated with 1 µg/mL doxycycline hyclate for 24 hours. To harvest, plates were moved to ice and cells were washed with 5 mL DPBS then mobilized with 0.25 mL TRIzol Reagent. Samples were stored at -80°C until further extraction.

Total RNA was extracted from homogenized tissues by the TRIzol Reagent procedure. RNA concentration and purity were assessed by spectrophotometric absorbance (NanoDrop One, ThermoFisher). First-strand cDNA was synthesized by reverse transcribing 100 ng of total RNA with a High-Capacity cDNA Reverse Transcription Kit (Life Technologies). Relative levels of mRNA were determined by quantitative real-time PCR (qRT-PCR) using the StepOnePlus real-time PCR system (Life Technologies) with reference gene Actin-beta and cycling parameters following the manufacturer's protocol of SYBR Green Universal Master Mix. Primers for *deaR* were designed using NCBI Primer-BLAST. Standard curves were prepared from serial dilutions of pooled concentrated cDNA with a cut-off of $R^2$>0.99 and efficiency between 90-110%.

Bulk RNA sequencing was performed following instructions from Plasmidsaurus. RNA was extracted from HeLa cells carrying the doxycycline-inducible DeaR construct that had been treated with 1 µg/mL doxycycline or vehicle for 48 h (n = 3 per group) and was submitted for RNA sequencing. Quality of the FASTQ files was assessed using FastQC v0.12.1. Reads were quality filtered using fastp v0.24.0 with poly-X tail trimming, 3′ quality-based tail trimming, a minimum Phred quality score of 15, and a minimum length requirement of 50 bp. Quality-filtered reads were aligned to the reference genome, supplemented with the *PA14_27675* transgene sequence as a custom contig, using STAR aligner v2.7.11 with non-canonical splice junction removal and output of unmapped reads, followed by coordinate sorting using samtools v1.22.1. PCR and optical duplicates were removed using UMI-based deduplication with UMIcollapse v1.1.0. Alignment quality metrics, strand specificity, and read distribution across genomic features were assessed using RSeQC v5.0.4 and Qualimap v2.3, with results aggregated into a comprehensive quality control report using MultiQC v1.32. Gene-level expression quantification was performed using featureCounts (subread package v2.1.1) with strand-specific counting, multi-mapping read fractional assignment, exons and three-prime UTRs as the feature identifiers, and grouping by gene_id. Final gene counts were annotated with gene biotype and other metadata extracted from the reference GTF. Sample–sample correlations for the correlation heatmap and PCA were calculated on TMM (trimmed mean of M-values) normalised counts using Pearson correlation. Induction of DeaR was confirmed from the transgene feature itself (0, 0.12 and 0 CPM without doxycycline versus 1.60, 1.68 and 1.21 CPM with doxycycline). Gene-level counts were analysed in Python 3.11.15.

<u>Recombinant protein expression and purification</u>

IPTG-inducible plasmids for expression of DeaR WT and the H22A variant were transformed into electrocompetent *E. coli* BL21 (DE3) and selected on LB plates containing 50 µg/mL kanamycin. Precultures were grown under standard conditions for 14-16 hours, then 5 mL were used to inoculate into 500 mL Overnight Express Instant LB Medium (Sigma-Aldrich). These overexpression cultures were incubated for 48 hours at 18 °C and 250 rpm. After harvesting by centrifugation at 4100x g for 30 minutes, the pellets were snap-frozen with liquid nitrogen and stored at -80°C until further purification.

Binding/wash buffer was prepared by adding imidazole to a final concentration of 40 mM in buffer (50 mM HEPES, 500 mM NaCl, and 5 mM TCEP pH 7.5). Elution buffer was prepared by adding imidazole to a final concentration of 500 mM into a HEPES buffer composed of 20 mM HEPES, 150 mM NaCl, 1 mM β-mercaptoethanol pH 7.5. Frozen cell pellets were thawed and resuspended in 30 mL of binding buffer, transferred to a 50 mL conical tube, and treated with one cOmplete EDTA-free protease inhibitor tablet (Roche) with gentle inversion. Lysozyme

(Sigma L-7651, 6 mg), DNase I (0.6 mg), and $MgCl_2$ (1 mM) were then added. The suspension was incubated at 4°C for 1 hour without agitation, followed by 30 min between 15-20k psi on a C3 Homogenizer (Avestin Emulsiflex). Lysates were transferred to 38 mL high-speed centrifuge tubes and clarified by centrifugation at 15,000 × g for 45 min in a pre-cooled (4 °C) centrifuge. The clarified supernatant was decanted into a fresh conical tube and carried forward directly into IMAC purification. With equilibrated His GraviTrap columns, the supernatant was loaded and washed 10 times with the column volume and eluted in 2 mL elution buffer. After overnight dialysis (Slide-A-Lyzer G3 Dialysis Cassettes, 20 KDa MWCO) in a buffer of 20 mM HEPES,150 mM NaCl, 1 mM β-Mercaptoethanol pH 7.5, the protein was purified with ion-exchange (Cytiva HiTrap Q High Performance 1 mL) on a gradient from 100-700 mM NaCl, then with size-exclusion chromatography (Cytiva Superose 6 Increase 10/300 GL 24 mL), and concentrated with Amicon Ultra centrifugal filter units (30 kDa MWCO to 1 mg/mL (high MW) or 2 mg/mL (low MW)). Protein concentration was measured by both PierceBCA assay (Thermo Scientific) and OD 280 NanoDrop. Protein identity was verified with SDS-PAGE and rapid instant Coomassie Blue Staining. Preparations were then aliquoted and snap-frozen with liquid nitrogen and stored at -80°C until further assay or analysis.

Sedimentation velocity analytical ultracentrifugation

Experiments were performed at 20°C using an Optima analytical ultracentrifuge (Beckman Coulter, Brea, CA) equipped with a TiAn50 rotor. Samples were loaded into two-channel, charcoal-filled Epon centerpieces with sapphire windows, and data were collected using both absorbance and interference optics. Loaded cells were allowed to equilibrate in the instrument under a temperature of 20 ºC and full vacuum for one hour before experiments were initiated. Sedimentation velocity centrifugation was performed at 40,000 rpm and 20 ºC. Refractive index and absorbance data (260, 280 nm) were collected simultaneously every 30 seconds to record the radial concentration as a function of time until all the sedimenting components cleared the optical path (~6 hours). Data were fit using the *c(S)* and *c(S,f/f₀)* implementations of the Lamm equation as implemented in the program SEDFIT version 17. The partial specific volume ($\bar{\upsilon}$), solvent density (ρ), and viscosity (η) were derived from chemical composition by SEDNTERP. Sedimentation coefficients were corrected for $S_{20,w}$. Measurements were performed in 20 mM HEPES-NaOH pH 7.5, 150 mM NaCl, and 1 mM DTT.

Thermal shift assay

Thermal shift assays of DeaR ion-exchange fractions (LMW and HMW) were performed with the Protein Thermal Shift Dye Kit (Thermo Fisher Scientific) following the manufacturer's protocol. Reactions (20 µL) contained 5 µM protein, 1× Protein Thermal Shift Buffer and 1× dye in 150 mM NaCl, 20 mM HEPES pH 7.4, 1 mM 2-mercaptoethanol (*105*), with 5 µM $ZnCl_2$ or no added metal in technical triplicate. Melting was monitored on a StepOnePlus Real-Time PCR System (Applied Biosystems) from 25 to 99 °C at 0.92 °C $min^{-1}$ with continuous acquisition in the ROX channel. Normalized reporter (R_n) traces were smoothed, baseline-corrected by subtracting an exponential fitted to the 27–41 °C and 82–96 °C flanks, and differentiated; T_m was taken as the maximum of d(corrected R_n)/dT. Metal conditions were paired by plate row and compared by two-tailed paired t-test with Bonferroni correction ($n = 3$).

Cryogenic electron microscopy (cryo-EM)

Specimens for cryo-EM analysis were prepared using a Vitrobot Mark IV plunge freezer (Thermo Fisher Scientific) at the Simons Electron Microscopy Center, New York Structural Biology Center (NYSBC). Purified DeaR was diluted to 0.3 mg/mL and supplemented with 5 mM AMP or 10 mM ADP. A 3 µL aliquot of this mixture was applied to a glow-discharged AuFlat 1.2/1.3 300-mesh grid (Electron Microscopy Sciences), blotted for 4 s at 25 °C and 95% humidity, and plunge-frozen in liquid ethane. Grid quality was screened on a Glacios microscope (Thermo Fisher Scientific) at the Columbia University Cryo-Electron Microscopy Center. A total of 5,351 image stacks were collected using Leginon (*106*) on a Titan Krios electron microscope (Thermo Fisher Scientific) equipped with a K3 direct electron detector (Gatan) at the National Center for CryoEM Access and Training (NCCAT), NYSBC. Movies were recorded at a calibrated pixel size of 0.828 Å, with a total dose of 54.25 $e^{-}/Å^2$ fractionated over 50 frames.

Cryo-EM image processing, model building and refinement

All image processing was performed in CryoSPARC (*107*), and a flowchart for the data processing is shown in **figure S5A**. Motion correction and dose weighting were applied to image stacks, and contrast transfer function (CTF) parameters were estimated using PatchCTF. A total of 1,205,209 particles were identified using the blob picker and subjected to multiple rounds of 2D classification. Particles from manually selected 2D classes (478,941 particles) were used to generate several ab initio 3D volumes, and they were used as the references for heterogeneous refinement. The resulting 330,345 particles for the good volume were used for Topaz training (*108*), which identified 785,497 particles from the full dataset. These particles were combined with the earlier ones, and after three additional rounds of 2D classification, 396,989 particles were subjected to 3D classification into five classes. Two classes showed density for the dodecamer (**figure S5B**) and were combined, and the resulting 217,948 particles were subjected to homogeneous refinement with tetrahedral (T) symmetry imposed, yielding a map at 2.31 Å resolution (**figure S5C,D**). The third class showed density for a nonamer (**figure S5E**), which probably arose from one of the trimers dissociating during vitrification. The remaining two classes had very few particles. An initial atomic model of the monomer of DeaR was generated using AlphaFold 3 prediction (*109*) and rigid-body fitted into the dodecamer cryo-EM density map using ChimeraX (*110*). Manual model adjustments were carried out in Coot (*111*). Iterative real-space refinement was performed using PHENIX (*112*) to optimize model–map agreement and stereochemistry. The cryo-EM data processing and structure refinement statistics are summarized in **table S2**.

Enzymatic assay

*In vitro* enzymatic assays with DeaR and the DeaR H22A variant were performed with protein at 5 µM with 1 mM nucleotide (ATP, ADP, AMP, GTP, or UTP) in the reaction buffer (20 mM HEPES pH 7.5, 150 mM NaCl, and 1 mM β-Mercaptoethanol). After one hour of incubation at 25°C and one hour of incubation at 37°C, reactions were quenched by adding extraction solvent (acetonitrile/methanol/water, 4:4:2, v/v/v) at a 1:100 ratio. Samples were centrifuged at 20,000 × g for 20 min at 4 °C, and the supernatant was transferred to clean microcentrifuge tubes. Extracts were dried under vacuum using a Genevac system for 2 h and stored at −80 °C until LC–MS analysis.

For LC-MS analysis, dried extracts were resuspended in 80 µL of a 6:4 acetonitrile:water mixture and 20 µL of 100% methanol, followed by vortexing, incubation at room temperature for 20 min, and centrifugation at 20,000 × g for 20 min at 4 °C. Nucleotides were analyzed using the same procedure described above in the section "Quantitative pHILIC LC-MS metabolomics".

Inductively Coupled Plasma-Mass Spectrometry

Metal concentrations in protein samples were quantified by inductively coupled plasma triple-quadrupole mass spectrometry at the METAL Laboratory, Department of Environmental Health Sciences, Columbia University. Sample aliquots were weighed into 15-mL metal-free polypropylene tubes (VWR), and 40 µL of an internal standard solution containing 500 µg/L Ga in 2% $HNO_3$ was added to each sample. Samples were diluted to a final volume of 2 mL with 10% $HNO_3$ and analyzed using an Agilent 8900 ICP-QQQ-MS. Mg, Mn, Fe, and Zn were measured in ammonia reaction-gas mode, whereas Ni was measured in helium collision-gas mode. Measured concentrations were corrected for the internal standard, calibration background, and a 10% $HNO_3$ procedural blank. Method detection limits were determined from six replicate blank measurements as 3.33 times the standard deviation of the blank and were adjusted for the sample dilution factor. Element concentrations were reported as both µg/L and µM and the occupancy was calculated by the molarity of metal over protein, measured by ICP-MS and BCA assay, respectively.

*Use of AI-assisted tools.* Initial exploratory analysis and draft visualizations were performed with the assistance of Claude (Opus 5.0, Anthropic) and Gemini (Pro 3.1, Google). The tools were used to suggest and generate analysis code and produce preliminary plots. All AI-generated code was reviewed by the authors, and every analysis and figure in this manuscript was regenerated and verified against the raw data. No figures or images were generated directly by AI tools. The authors take full responsibility for the accuracy and integrity of all reported results.
