## Supplementary material for "*Pseudomonas aeruginosa* R-bodies deliver an ADP deaminase toxin": figures S1-7

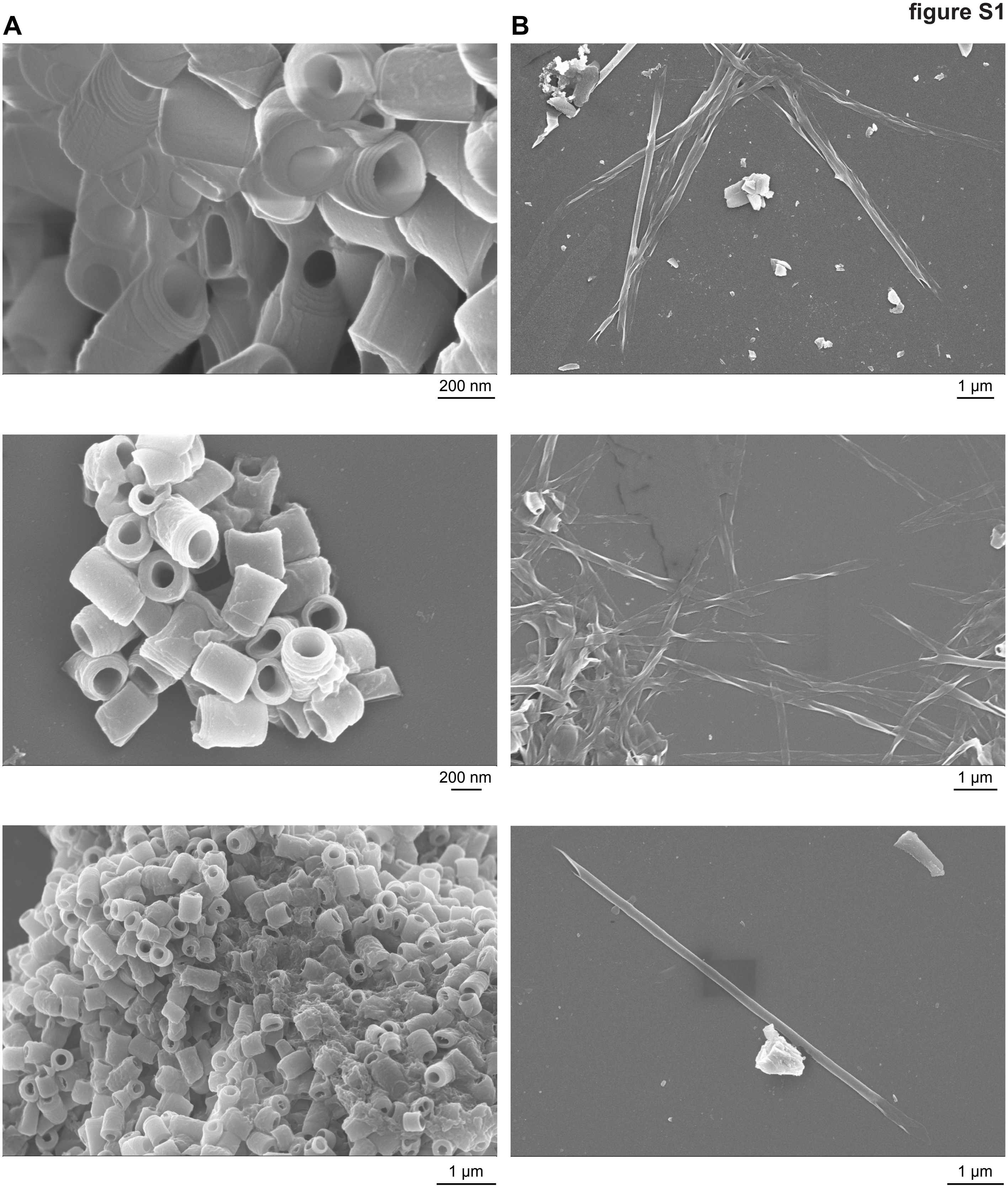


**figure S1. Scanning electron microscopy (SEM) of purified R-bodies. (A)** Contracted R-bodies purified from *P. aeruginosa* PA14, fixed, and dehydrated for SEM imaging. **(B)** R-bodies extended by buffer exchange to acidic pH and subsequently prepared for SEM imaging as described in (A). Some extended R-bodies show apparent unraveling, likely due to artifacts introduced during sample preparation or imaging.

### figure S2

**Maximum-likelihood tree**

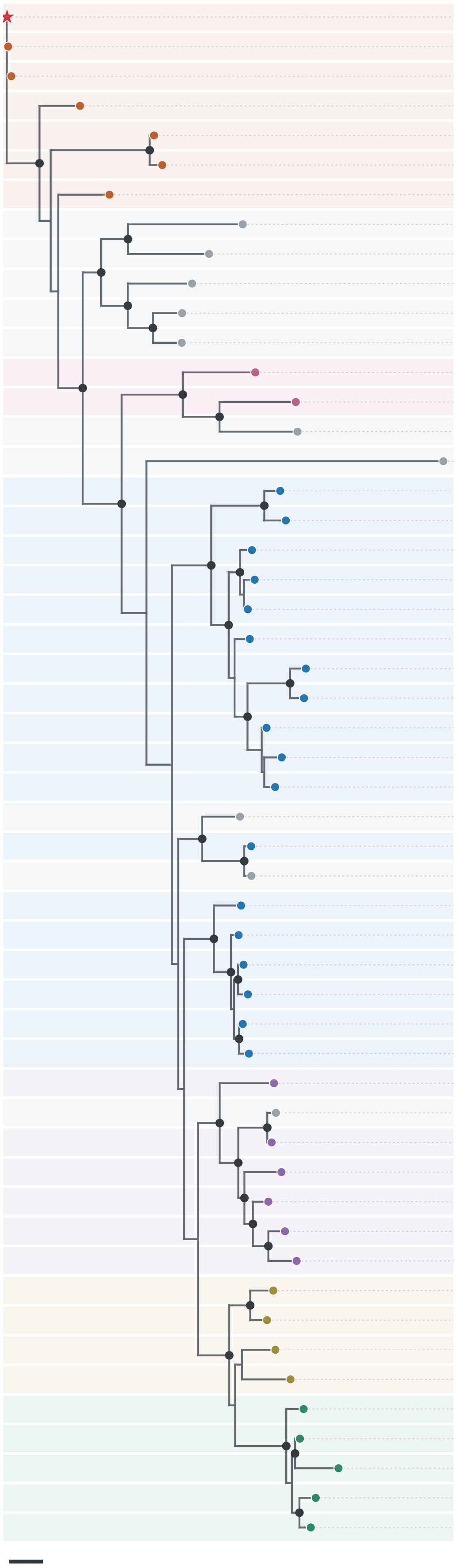

0.3 substitutions/site

| Organism | Accession / locus |
|---|---|
| ***Pseudomonas aeruginosa* PA14** | PA14_27675 (DeaR) |
| *Pseudomonas aeruginosa* | RCI74724.1 |
| *Pseudomonas aeruginosa* | WP_427123833.1 |
| *Pseudomonas indica* | WP_084336223.1 |
| *Pseudomonas aeruginosa* | WP_147318657.1 |
| *Pseudomonas* | WP_063861531.1 |
| *Pseudomonas mangiferae* | WP_143488276.1 |
| *Leminorella grimontii* | WP_347253499.1 |
| *Enterobacteriaceae bacterium* | MIL2803844.1 |
| *Zoogloeaceae bacterium* | MDR1648598.1 |
| *Betaproteobacteria bacterium* | GHU44898.1 |
| *Azoarcus sp.* | MDR1461752.1 |
| *Chromobacterium subtsugae* | WP_152680237.1 |
| *Chromobacterium violaceum* | WP_210562799.1 |
| *Zoogloea sp.* | WP_374489511.1 |
| *Pseudophaeobacter sp.* | WP_348342893.1 |
| *Lysobacter sp.* K5869 | WP_215839186.1 |
| *Lysobacter enzymogenes* | WP_096376580.1 |
| *Lysobacter capsici* | WP_046659987.1 |
| *Lysobacter capsici* | WP_096412691.1 |
| *Lysobacter sp.* Root690 | WP_056115883.1 |
| unclassified *Lysobacter* | WP_215056721.1 |
| *Lysobacter firmicutimachus* | WP_336132632.1 |
| *Lysobacter sp.* BMK333-48F3 | WP_221671195.1 |
| *Lysobacter antibioticus* | WP_031371089.1 |
| *Lysobacter sp.* 1R34A | WP_435397251.1 |
| *Lysobacter sp.* CA196 | WP_441329225.1 |
| Lysobacteraceae bacterium | TXH67356.1 |
| *Lysobacter sp.* | TXI50555.1 |
| Lysobacteraceae bacterium | MBP6749686.1 |
| *Lysobacter silvisoli* | WP_115861653.1 |
| *Lysobacter sp.* Root983 | WP_057160971.1 |
| unclassified *Lysobacter* | WP_056305964.1 |
| *Lysobacter sp.* cf310 | WP_091637811.1 |
| unclassified *Lysobacter* | WP_332939814.1 |
| *Lysobacter sp.* 5GHs7-4 | WP_232908956.1 |
| *Dyella terrae* | WP_131151070.1 |
| *Frateuria sp.* sgz306004 | WP_488384094.1 |
| *Dyella sp.* ASV21 | WP_199099733.1 |
| *Dyella sp.* AtDHG13 | WP_074547957.1 |
| *Dyella sedimenti* | WP_243041871.1 |
| *Dyella kyungheensis* | WP_426270943.1 |
| *Dyella terrae* | WP_239947954.1 |
| *Luteimonas sp.* | MGO1540546.1 |
| *Luteimonas sp.* R10 | WP_407353863.1 |
| *Luteimonas salinilitoris* | WP_370563838.1 |
| *Luteimonas sp.* | MGH8032559.1 |
| *Xanthomonas bonasiae* | WP_206230736.1 |
| *Xanthomonas bonasiae* | WP_191824300.1 |
| *Xanthomonas bundabergensis* | WP_369980036.1 |
| *Xanthomonas sacchari* | WP_267114490.1 |
| *Xanthomonas theicola* | WP_128420886.1 |

**figure S2. Phylogenetic relationships among DeaR homologs**. Rectangular representation of the unrooted maximum-likelihood tree (**Figure 2B**) of 52 proteins, with organism names and protein accessions indicated. Tip colors and row shading denote the same taxonomic groups as in the main figure. DeaR (PA14_27675) is highlighted in red. Black dots indicate SH-aLRT support ≥80 and ultrafast bootstrap support ≥95. Branch lengths represent amino acid substitutions per site; scale bar is 0.3. The root of the tree is arbitrary and does not imply evolutionary direction.

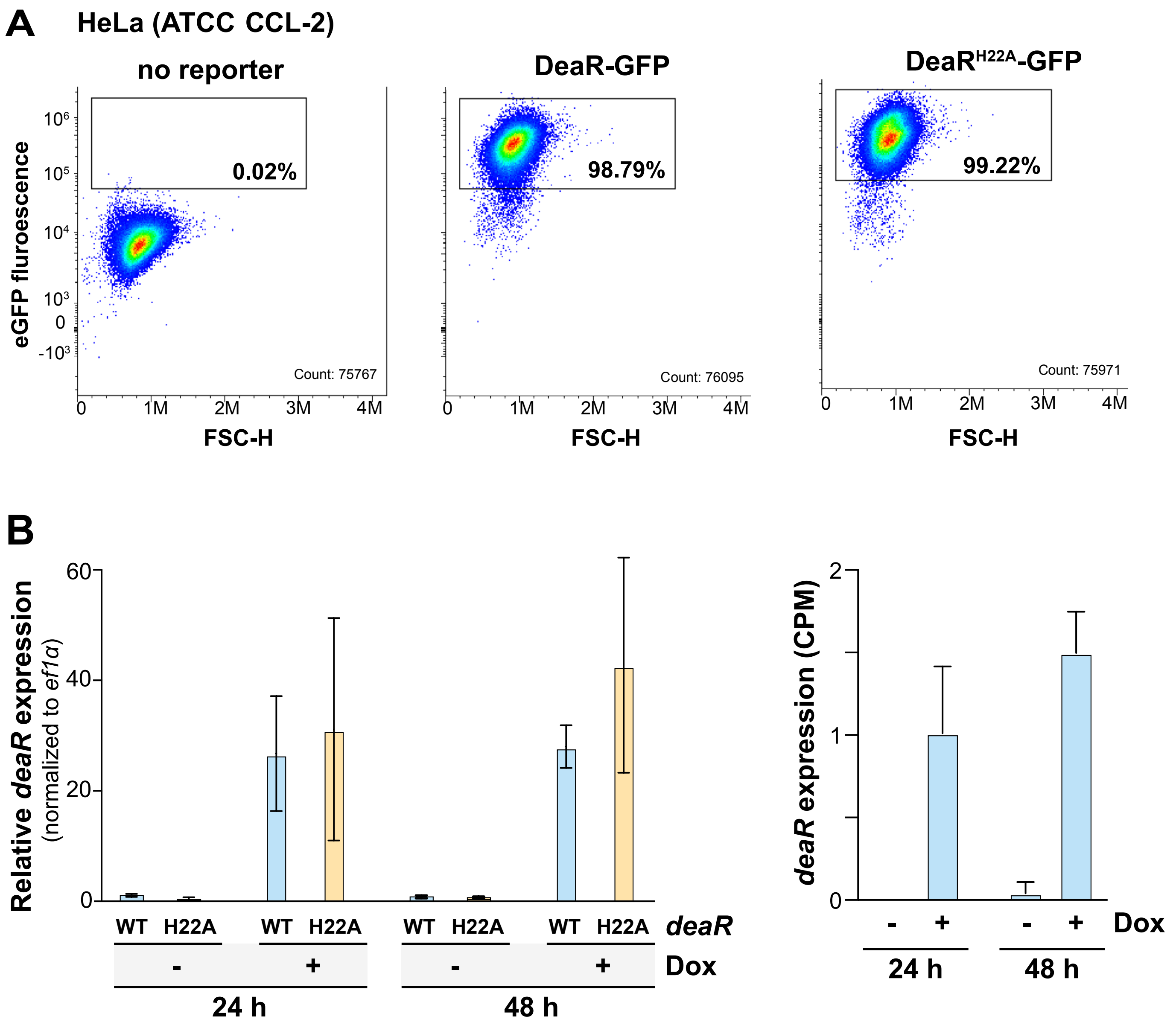


**figure S3. Doxycycline-inducible expression of WT DeaR and the H22A variant in HeLa cells. (A)** Fluorescence-activated cell sorting (FACS) analysis of eGFP fluorescence in HeLa cells carrying pInducer21 constructs for inducible expression of WT DeaR-eGFP or the H22A variant. eGFP-positive cells were sorted and collected for downstream assays. HeLa cells lacking the reporter served as a negative control. Percentages indicate the proportion of cells within the indicated fluorescence gate (0.02%, 98.79%, and 99.22%, respectively).

**(B)** Quantification of *deaR* transcript abundance in HeLa cells following doxycycline (Dox) induction. **Left:** Relative *deaR* transcript abundance (assessed by RT-qPCR) in cells expressing WT DeaR or the H22A variant, normalized to *eif1α*, after 24 or 48 h in the presence (+) or absence (−) of Dox. **Right:** *deaR* transcript abundance (assessed by RNAseq), expressed as transcripts per million (TPM), in cells expressing WT DeaR after 24 or 48 h with or without Dox. Error bars represent standard deviations.

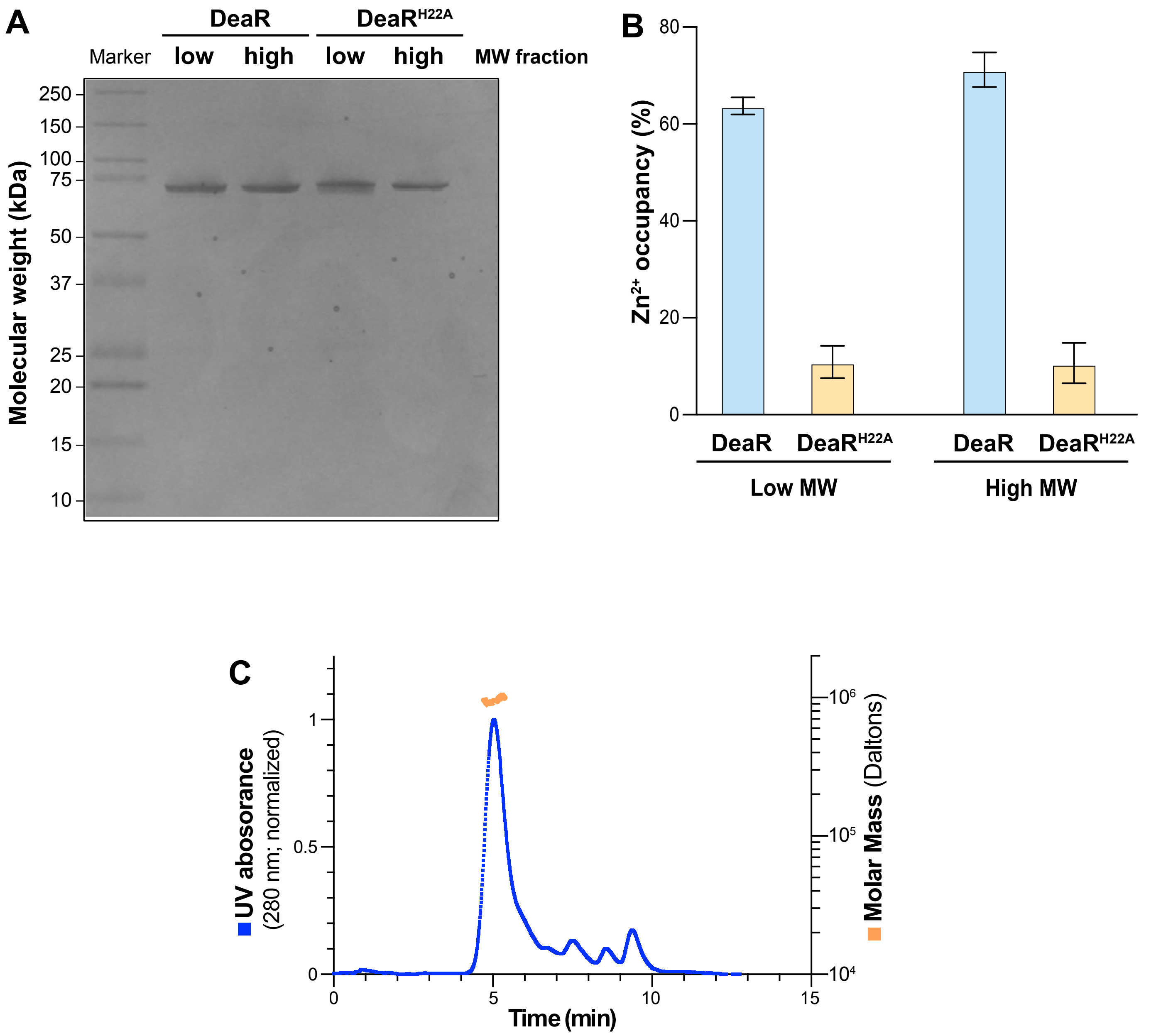


**figure S4. Biochemical characterization of WT DeaR and the H22A variant**.
**(A)** Coomassie-stained SDS-PAGE gel of low- and high-molecular-mass fractions of recombinant WT DeaR purified from *E. coli* by nickel-affinity chromatography and subsequently separated by ion-exchange chromatography. SDS-PAGE was performed under reducing conditions. **(B)** Inductively coupled plasma mass spectrometry (ICP-MS) analysis of metal content in the low- and high-molecular-mass fractions of WT and H22A DeaR. **(C)** Size-exclusion chromatography coupled to multi-angle light scattering (SEC-MALS) analysis of the oligomeric state and molecular mass of the high-molecular-mass fraction of WT DeaR.

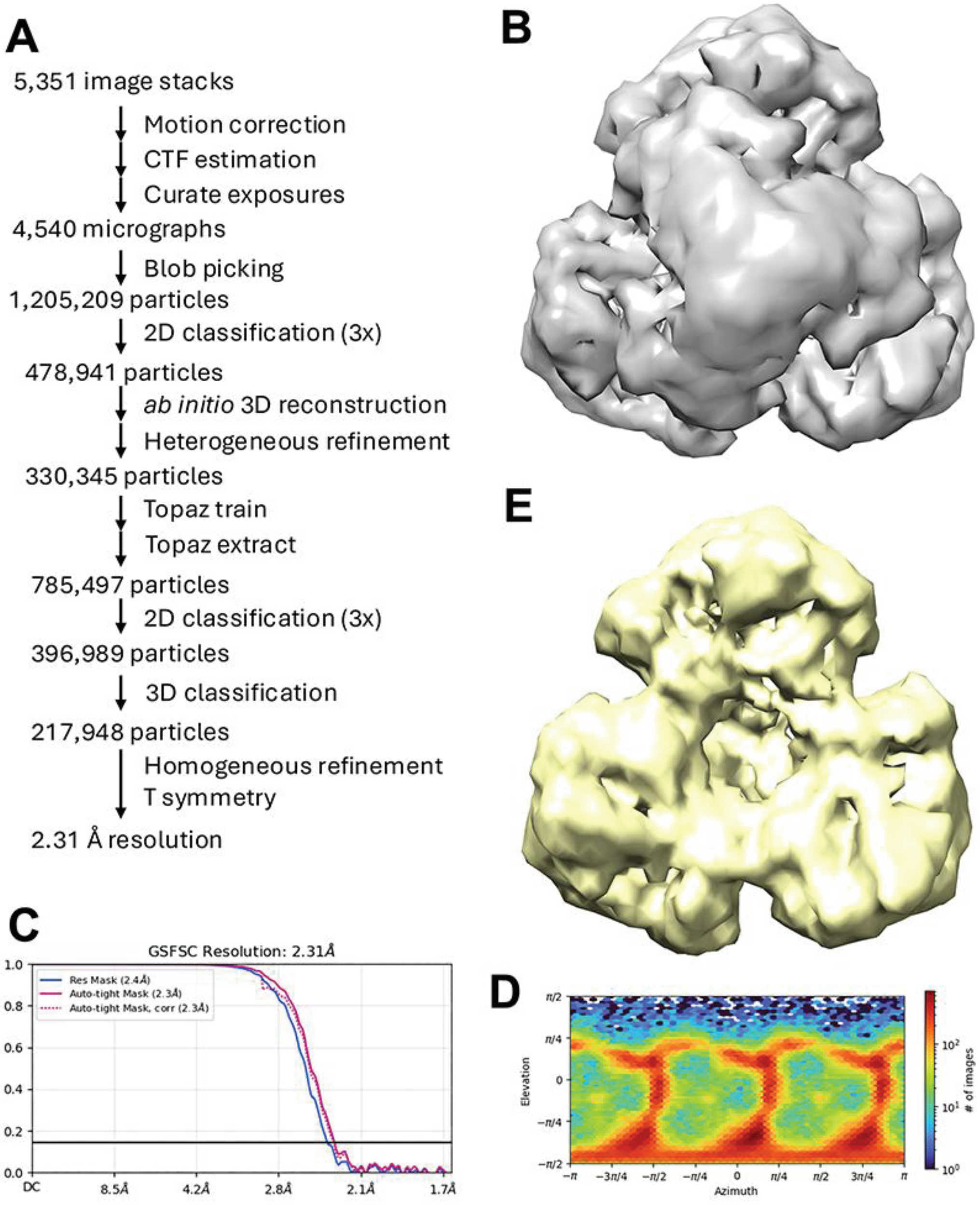


**figure S5. Cryo-EM data processing for Pseudomonas aeruginosa PA14 DeaR.** (A). Flowchart for EM data processing. (B). EM density for a dodecamer after 3D classification. (C). Fourier shell correlation curves for the final reconstruction. Tetrahedral symmetry was enforced. (D). The orientation distribution of particles used the reconstruction. (E). EM density for a nonamer after 3D classification. One trimer is missing in the density.

**figure S6**

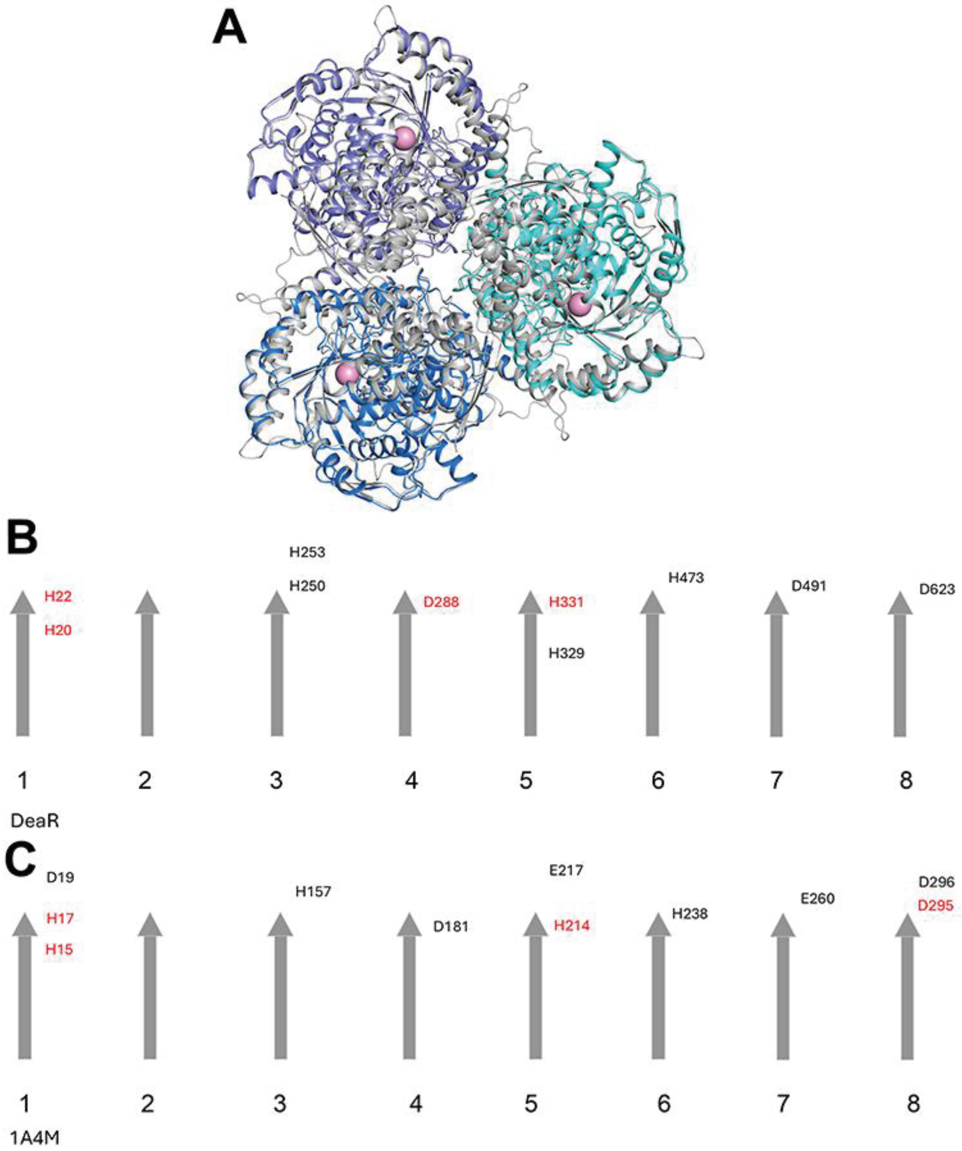


**figure S6.** (**A**) Overlay of the cryo-EM structure of DeaR trimer (in color) with that from AlphaFold 3 prediction (gray). The AlphaFold 3 prediction for the trimer is similar to the cryo-EM structure. (**B**) Topological drawing showing the locations of His and Asp/Glu residues near the active site of DeaR. The eight β strands of the TIM barrel are shown as arrows (gray). The ligands to the zinc ion are shown in red. The His and Asp/Glu residues are located near the end of the β strands, either in the strands or in the following loops. (**C**) Locations of His and Asp/Glu residues near the active site of mouse adenosine deaminase (PDB entry 1A4M). A large number of His and Asp/Glu residues are located in the active site region in these enzymes. The Asp288 ligand in DeaR is topologically equivalent to Asp181 in 1A4M, but the side chain of Asp181 is 5.8 Å away from zinc. On the other hand, the Asp295 ligand of 1A4M is equivalent to Asp623 of DeaR, originally identified as a zinc ligand based on AF3 predictions and structural comparisons (Fig. 2A). However, Asp623 is 7 Å away from the zinc ion and has weak EM density in our structure (Fig. 4E).

**Figure S7. Chromosomal alignment of *reb* clusters represented in Figure 5A.**

**Movie S1.** Movie showing a *D. discoideum* amoeba incubated with PA14 and *K. pneumoniae* bacteria. Following the collapse of an internal compartment, an R-body can be seen protruding from *D. discoideum* and persisting in time. Time is displayed in seconds and the movie is sped up 10.72x from real time.

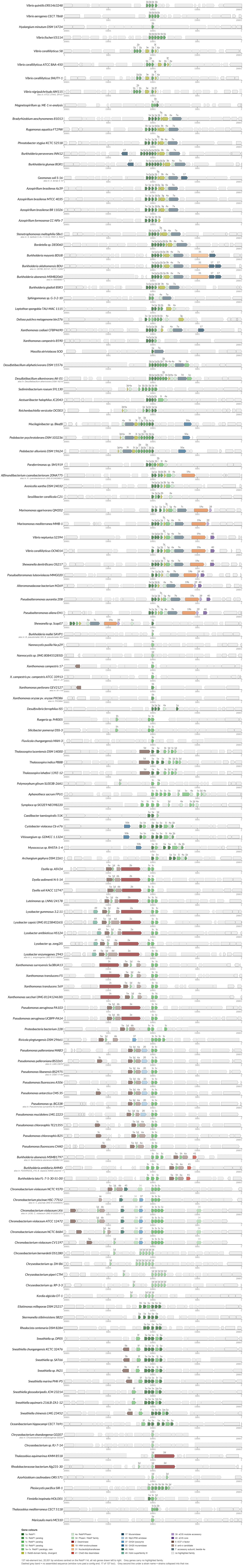


137 reb-element loci, 20,001 bp windows centred on the RebP1 hit, all reb genes drawn left to right. Grey genes carry no highlighted family.
Dashed grey band = no assembled sequence (window runs past a contig end; 17 of 137 loci). Grey second line under a strain name = strains collapsed into that row.
