## Supplementary material for "*Pseudomonas aeruginosa* R-bodies deliver an ADP deaminase toxin": table S1

**Table S1. Strains used in this study**

| Strain | Number | Description | Source | Appears in figure |
|---|---|---|---|---|
| *Escherichia coli* | | | | |
| *E. coli DH10B* | EC0113 | | Thermo Scientific | |
| *E. coli BL21(DE3)* | C600003 | | Invitrogen™ | |
| *E. coli UQ950* | LD44 | *E. coli* DH5α λ(*pir*) strain for cloning; FΔ(*argF-lac*) 169φ80*dlacZ*58(ΔM15) *glnV44*(AS) *rfbD1 gyrA96*(NalR) *recA1 endA1 spoT thi-1 hsdR17 deoR λpir+* | D. Lies, Caltech | |
| *ß2155* | LD69 | Helper strain. *thrB1004 pro thi strA hsdsS lacZ*ΔM15 (F' *lacZ*ΔM15 *lacIq traD36 proA+ proB+*) Δ*dapA::erm* (Ermr ) *pir::*RP4 [::kan (Kmr ) from SM10] | (91) | |
| *E. coli BW29427* | LD661 | Donor strain for biparental conjugation; t*hrB1004 pro thi rpsL hsdS lacZ* ΔM15RP4-1360 Δ(*araBAD*)567 *ΔdapA1341*::[*erm pir*(wt)] | W. Metcalf, University of Illinois | |
| *Saccharomyces cerevisiae* | | | | |
| *S. cerevisiae InvSc1* | LD676 | *MATa/MATα leu2/leu2 trp1-289/trp1-289 ura3-52/ ura3-52 his3-Δ1/his3-Δ1* | Invitrogen™ | |
| *Pseudomonas aeruginosa* | | | | |
| *UCBPP-PA14* | LD0 | NC_008463 | (8) | 1g, 2e |
| *rhaSR-PrhaBAD-rcgA* | LD5163 | PA14 with *pAKN69-rhaSR-P$_{rhaBAD}$-rcgA* inserted at the *glmS* site. Made by mating LD5667 into PA14 | This paper | 1b |
| *pAKN69-rcgA* | LD4613 | PA14 with P$_{PA1/04/03}$-*rcgA* inserted at the *glmS* site. Made by mating *pAKN69-rcgA* into PA14 | This paper | 1e |
| *PrebP-mScarlet* | LD3224 | PA14 with PrebP1-mScarlet inserted at the attB site. | (9) | 1f |
| *pAKN-gfp* | LD3801 | PA14 with *P$_{PA1/04/03}$-gfp* inserted at the glmS site | (9) | 1f |
| ∆*rebP1rebP2* | LD2935 | PA14 with deletion in *PA14_27640* (*rebP1*)-*27630* (*rebP2*). Made by mating pLD2934 into UCBPP-PA14. | (9) | 1g |
| *gfp-rebP2* | LD3199 | PA14 with insertion of *BBaB003-gfp-rebP2* adjacent to the native coding region of *rebP1* and *rebP2* | (9) | |
| ∆*rcgAfecI2* | LD3137 | PA14 with deletion in *PA14_27690* (*fecI2*)-*27700* (*rcgA*). Made by mating LD3136 into *UCBPP-PA14.* | This paper | 1g |
| ∆*deaR* | LD3699 | PA14 with deletion in *PA14_27675* (*deaR*). Made by mating LD3684 into *UCBPP-PA14.* | This paper | 2e |
| ∆*deaR::deaR* | LD5473 | PA14 Δ*PA14_27675* (*deaR*) strain with wild-type *PA14_27675* complemented back into the site of deletion. Made by mating LD5556 into LD3699. | This paper | 2e |
| ∆*deaR::H22A-deaR* | LD5488 | PA14 Δ*PA14_27675* (*deaR*) strain with *H22A-PA14_27675* complemented back into the site of deletion. Made by mating LD5557 into LD3699. | This paper | 2e |
| *pAKN-rcgA DeaR -mScarlet GFP-RebP2* | LD5558 | PA14 with *pAKN69-rcgA* inserted at the *glmS* site, insertion of *mScarlet* adjacent to the native locus of *PA14_27675*, and insertion of *BBaB003-gfp-rebP2* adjacent to the native coding region of *rebP1* and *rebP2* | This paper | 2f,g,h |
| *Klebsiella pneumoniae* | | | | |
| *DBS0351098* | LD3978 | *Klebsiella pneumoniae* from Dictybase | Dictybase | 1f,g,2e |
| *Homo sapien* | | | | |
| HeLa | Cat# CCL-2; RRID:CVCL_0 030 | | ATCC | 3a |
| HEK 293T | Cat# CRL-3216 ™ | | ATCC | |
| *Dictyostelium discoideum* | | | | |
| *Ax4* | LD3985b | *Dictyostelium discoideum Ax4* from Dictybase | Dictybase | 1e,f,g 2e |
