## Supplementary material for "*Pseudomonas aeruginosa* R-bodies deliver an ADP deaminase toxin": table S2

**Table S2. Summary of cryo-EM data processing and structure refinement statistics**

| **Structure** | ***P. aeruginosa* PA14 DeaR dodecamer** |
|---|---|
| **Data collection and processing** | |
| Voltage (kV) | 300 |
| Electron exposure ($e^{-}$/Å$^{2}$) | 54.25 |
| Movies | 5,351 |
| Initial particle images | 1,205,209 |
| Final particle images | 217,948 |
| Symmetry imposed | T |
| Map resolution (Å) | 2.31 |
| FSC threshold | 0.143 |
| Map sharpening *B* factor (Å$^{2}$) | –77.4 |
| **Structure refinement** | |
| No. of protein residues | 6,492 |
| No. of atoms | 51,564 |
| No. of metal ions | 12 |
| rms deviations | |
| Bond lengths (Å) | 0.005 |
| Bond angles (°) | 0.82 |
| Clash score | 6 |
| Poor rotamers (%) | 3 |
| Ramachandran plot | |
| Favored (%) | 97 |
| Allowed (%) | 3 |
| Disallowed (%) | 0 |
