## Supplementary material for "*Pseudomonas aeruginosa* R-bodies deliver an ADP deaminase toxin": data S3

### file S3. Annotated plasmid sequences

Plasmids used in this study. Each entry gives the plasmid name, a description, the backbone and source, an annotated feature list, and the complete double-stranded sequence (one strand shown, 5'->3', 60 nt per line). Coordinates are 1-based and inclusive; 'c(x..y)' denotes a feature on the complementary strand.

#### Contents


| | Plasmid | Length | Source |
|---|---|---|---|
| 1. | pINDUCER21 (ORF-EG) | 12,145 bp | Addgene plasmid #46948 |
| 2. | pINDUCER21 (ORF-EG)_27675_WT | 12,645 bp | This paper |
| 3. | pINDUCER21 (ORF-EG)_27675_H22A | 12,646 bp | This paper |
| 4. | pKG001 (pET-ccdB-GoldenGate) | 6,060 bp | This paper |
| 5. | LD5562 (pRTP001_IPTG_WT_27675) | 7,412 bp | This paper |
| 6. | LD5563 (pRTP010_27675_HA_IPTG) | 7,412 bp | This paper |
| 7. | pMQ30-Δ27675 | 9,178 bp | This paper |
| 8. | pMQ30-ΔrcgAfecI2 | 9,183 bp | This paper |
| 9. | pMQ30-H22A-27675 | 11,373 bp | This paper |
| 10. | pMQ30-Δ27675::27675 | 11,286 bp | This paper |
| 11. | pMQ30-27675-mScarlet | 9,880 bp | This paper |
| 12. | pAKN69-rhaSR-PrhaBAD-rcgA | 9,725 bp | This paper |
| 13. | pAKN69-rcgA | 7,798 bp | This paper |

#### 1. pINDUCER21 (ORF-EG)

**Description** Doxycycline-inducible lentiviral Gateway destination vector (TRE-attR1/CmR-ccdB/attR2, EF-1alpha-rtTA-Advanced, IRES-EGFP). Parent vector; sequence as deposited (Addgene-verified full sequence 392924).

**Length** 12,145 bp, circular

**Source** Addgene plasmid #46948

**Note** *Features are as deposited. The 5'-proximal LTR at 1..634 is shown here as "5' LTR"; the deposited annotation calls it "3' LTR" because in this vector design the 5' LTR is derived from the 3' LTR.*

##### Features

| Location | Type | Feature |
|---|---|---|
| 1..634 | LTR | 5' LTR |
| 681..806 | misc_feature | HIV-1 Psi |
| 1303..1536 | misc_feature | RRE |
| 1721..1765 | CDS | gp41 peptide |
| 2058..2175 | misc_feature | cPPT/CTS |
| 2230..2500 | protein_bind | tetracycline response element |
| 2533..2571 | promoter | minimal CMV promoter |
| 2701..2825 | protein_bind | attR1 |
| 2850..2880 | promoter | lac UV5 promoter |
| 2934..3593 | CDS | cat (chloramphenicol resistance) |
| 3935..4240 | CDS | ccdB |
| c(4281..4405) | protein_bind | attR2 |
| 4458..4484 | CDS | HA |
| 4704..5882 | promoter | EF-1-alpha promoter |
| 4935..5873 | intron | EF-1-alpha intron A |
| 5952..5985 | protein_bind | loxP |
| 5992..6738 | CDS | rtTA-Advanced |
| 6895..7357 | misc_feature | IRES |
| 7373..8092 | CDS | EGFP |
| 8535..9123 | misc_feature | WPRE |
| c(9006..9017) | CDS | Factor Xa site |
| 9198..9431 | LTR | 3' LTR (Delta-U3) |
| 9636..9740 | promoter | bla promoter |
| 9741..10601 | CDS | bla (ampicillin resistance) |
| 10772..11360 | rep_origin | ori |
| 11771..11792 | protein_bind | CAP binding site |
| 11807..11837 | promoter | lac promoter |
| 11845..11861 | protein_bind | lac operator |

##### Sequence (12,145 bp)

1 tggaagggct aattcactcc caaagaagac aagatatcct tgatctgtgg atctaccaca
61 cacaaggcta cttccctgat tagcagaact acacaccagg gccaggggtc agatatccac
121 tgacctttgg atggtgctac aagctagtac cagttgagcc agataaggta gaagaggcca
181 ataaaggaga gaacaccagc ttgttacacc ctgtgagcct gcatgggatg gatgacccgg
241 agagagaagt gttagagtgg aggtttgaca gccgcctagc atttcatcac gtggcccgag
301 agctgcatcc ggagtacttc aagaactgct gatatcgagc ttgctacaag ggactttccg
361 ctggggactt tccagggagg cgtggcctgg gcgggactgg ggagtggcga gccctcagat
421 cctgcatata agcagctgct ttttgcctgt actgggtctc tctggttaga ccagatctga
481 gcctgggagc tctctggcta actagggaac ccactgctta agcctcaata aagcttgcct
541 tgagtgcttc aagtagtgtg tgcccgtctg ttgtgtgact ctggtaacta gagatccctc
601 agaccctttt agtcagtgtg gaaaatctct agcagtggcg cccgaacagg gacttgaaag
661 cgaaagggaa accagaggag ctctctcgac gcaggactcg gcttgctgaa gcgcgcacgg
721 caagaggcga ggggcggcga ctggtgagta cgccaaaaat tttgactagc ggaggctaga
781 aggagagaga tgggtgcgag agcgtcagta ttaagcgggg gagaattaga tcgcgatggg
841 aaaaaattcg gttaaggcca gggggaaaga aaaaatataa attaaaacat atagtatggg
901 caagcaggga gctagaacga ttcgcagtta atcctggcct gttagaaaca tcagaaggct
961 gtagacaaat actgggacag ctacaaccat cccttcagac aggatcagaa gaacttagat
1021 cattatataa tacagtagca accctctatt gtgtgcatca aaggatagag ataaaagaca
1081 ccaaggaagc tttagacaag atagaggaag agcaaaacaa aagtaagacc accgcacagc
1141 aagcggccgg ccgctgatct tcagacctgg aggaggagat atgagggaca attggagaag
1201 tgaattatat aaatataaag tagtaaaaat tgaaccatta ggagtagcac ccaccaaggc
1261 aaagagaaga gtggtgcaga gagaaaaaag agcagtggga ataggagctt tgttccttgg
1321 gttcttggga gcagcaggaa gcactatggg cgcagcgtca atgacgctga cggtacaggc
1381 cagacaatta ttgtctggta tagtgcagca gcagaacaat ttgctgaggg ctattgaggc
1441 gcaacagcat ctgttgcaac tcacagtctg gggcatcaag cagctccagg caagaatcct
1501 ggctgtggaa agatacctaa aggatcaaca gctcctgggg atttggggtt gctctggaaa
1561 actcatttgc accactgctg tgccttggaa tgctagttgg agtaataaat ctctggaaca
1621 gatttggaat cacacgacct ggatggagtg ggacagagaa attaacaatt acacaagctt
1681 aatacactcc ttaattgaag aatcgcaaaa ccagcaagaa aagaatgaac aagaattatt
1741 ggaattagat aaatgggcaa gtttgtggaa ttggtttaac ataacaaatt ggctgtggta
1801 tataaaatta ttcataatga tagtaggagg cttggtaggt ttaagaatag tttttgctgt
1861 actttctata gtgaatagag ttaggcaggg atattcacca ttatcgtttc agacccacct
1921 cccaaccccg aggggacccg acaggcccga aggaatagaa gaagaaggtg gagagagaga
1981 cagagacaga tccattcgat tagtgaacgg atctcgacgg tatcgccgaa ttcacaaatg
2041 gcagtattca tccacaattt taaaagaaaa ggggggattg gggggtacag tgcaggggaa
2101 agaatagtag acataatagc aacagacata caaactaaag aattacaaaa acaaattaca
2161 aaaattcaaa attttcgggt ttattacagg gacagcagag atccagtttg gactaggatc
2221 ctttaccact ccctatcagt gatagagaaa agtgaaagtc gagtttacca ctccctatca
2281 gtgatagaga aaagtgaaag tcgagtttac cactccctat cagtgataga gaaaagtgaa
2341 agtcgagttt accactccct atcagtgata gagaaaagtg aaagtcgagt ttaccactcc
2401 ctatcagtga tagagaaaag tgaaagtcga gtttaccact ccctatcagt gatagagaaa
2461 agtgaaagtc gagtttacca ctccctatca gtgatagaga aaagtgaaag tcgagctcgg
2521 tacccgggtc gaggtaggcg tgtacggtgg gaggcctata taagcagagc tcgtttagtg
2581 aaccgtcaga tcgcctggag acgccatcca cgctgttttg acctccatag aagacaccgg
2641 gaccgatcca gcctccgcgg ccccgaacta gtccagtgtg gtggaattct gcagatatca
2701 acaagtttgt acaaaaaagc tgaacgagaa acgtaaaatg atataaatat caatatatta
2761 aattagattt tgcataaaaa acagactaca taatactgta aaacacaaca tatccagtca
2821 ctatggcggc cgcattaggc accccaggct ttacacttta tgcttccggc tcgtataatg
2881 tgtggatttt gagttaggat ccgtcgagat tttcaggagc taaggaagct aaaatggaga
2941 aaaaaatcac tggatatacc accgttgata tatcccaatg gcatcgtaaa gaacattttg
3001 aggcatttca gtcagttgct caatgtacct ataaccagac cgttcagctg gatattacgg
3061 cctttttaaa gaccgtaaag aaaaataagc acaagtttta tccggccttt attcacattc
3121 ttgcccgcct gatgaatgct catccggaat tccgtatggc aatgaaagac ggtgagctgg
3181 tgatatggga tagtgttcac ccttgttaca ccgttttcca tgagcaaact gaaacgtttt
3241 catcgctctg gagtgaatac cacgacgatt tccggcagtt tctacacata tattcgcaag
3301 atgtggcgtg ttacggtgaa aacctggcct atttccctaa agggtttatt gagaatatgt
3361 ttttcgtctc agccaatccc tgggtgagtt tcaccagttt tgatttaaac gtggccaata
3421 tggacaactt cttcgccccc gttttcacca tgggcaaata ttatacgcaa ggcgacaagg
3481 tgctgatgcc gctggcgatt caggttcatc atgccgtttg tgatggcttc catgtcggca
3541 gaatgcttaa tgaattacaa cagtactgcg atgagtggca gggcggggcg taaagatctg
3601 gatccggctt actaaaagcc agataacagt atgcgtattt gcgcgctgat ttttgcggta
3661 taagaatata tactgatatg tatacccgaa gtatgtcaaa aagaggtatg ctatgaagca
3721 gcgtattaca gtgacagttg acagcgacag ctatcagttg ctcaaggcat atatgatgtc
3781 aatatctccg gtctggtaag cacaaccatg cagaatgaag cccgtcgtct gcgtgccgaa
3841 cgctggaaag cggaaaatca ggaagggatg gctgaggtcg cccggtttat tgaaatgaac
3901 ggctcttttg ctgacgagaa caggggctgg tgaaatgcag tttaaggttt acacctataa
3961 aagagagagc cgttatcgtc tgtttgtgga tgtacagagt gatattattg acacgcccgg

4021 gcgacggatg gtgatccccc tggccagtgc acgtctgctg tcagataaag tctcccgtga
4081 actttacccg gtggtgcata tcggggatga aagctggcgc atgatgacca ccgatatggc
4141 cagtgtgccg gtctccgtta tcggggaaga agtggctgat ctcagccacc gcgaaaatga
4201 catcaaaaac gccattaacc tgatgttctg gggaatataa atgtcaggct cccttataca
4261 cagccagtct gcaggtcgac catagtgact ggatatgttg tgttttacag tattatgtag
4321 tctgtttttt atgcaaaatc taatttaata tattgatatt tatatcattt tacgtttctc
4381 gttcagcttt cttgtacaaa gtggttgata tccagcacag tggcggccgc tcgagtctag
4441 agggcccgcg gttcgaatac ccatacgacg tcccagacta cgcttagtaa tgattaatta
4501 aactagaaat tctaccgggt aggggaggcg cttttcccaa ggcagtctgg agtaacccac
4561 ccaagatctg gcctccgcgc cgggttttgg cgcctcccgc gggcgccccc ctcctcacgg
4621 cgagccgcgt aagatctggc ctccgcgccg ggttttggcg cctcccgcgg gcgcccccct
4681 cctcacggcg agcctagtcg tgaggctccg gtgcccgtca gtgggcagag cgcacatcgc
4741 ccacagtccc cgagaagttg gggggagggg tcggcaattg aaccggtgcc tagagaaggt
4801 ggcgcggggt aaactgggaa agtgatgtcg tgtactggct ccgccttttt cccgagggtg
4861 ggggagaacc gtatataagt gcagtagtcg ccgtgaacgt tctttttcgc aacgggtttg
4921 ccgccagaac acaggtaagt gccgtgtgtg gttcccgcgg gcctggcctc tttacgggtt
4981 atggcccttg cgtgccttga attacttcca cctggctgca gtacgtgatt cttgatcccg
5041 agcttcgggt tggaagtggg tgggagagtt cgaggccttg cgcttaagga gccccttcgc
5101 ctcgtgcttg agttgaggcc tggcctgggc gctggggccg ccgcgtgcga atctggtggc
5161 accttcgcgc ctgtctcgct gctttcgata agtctctagc catttaaaat ttttgatgac
5221 ctgctgcgac gctttttttc tggcaagata gtcttgtaaa tgcgggccaa gatctgcaca
5281 ctggtatttc ggtttttggg gccgcgggcg gcgacggggc ccgtgcgtcc cagcgcacat
5341 gttcggcgag gcggggcctg cgagcgcggc caccgagaat cggacggggg tagtctcaag
5401 ctggccggcc tgctctggtg cctggcctcg cgccgccgtg tatcgccccg ccctgggcgg
5461 caaggctggc ccggtcggca ccagttgcgt gagcggaaag atggccgctt cccggccctg
5521 ctgcagggag ctcaaaatgg aggacgcggc gctcgggaga gcgggcgggt gagtcaccca
5581 cacaaaggaa aagggccttt ccgtcctcag ccgtcgcttc atgtgactcc acggagtacc
5641 gggcgccgtc caggcacctc gattagttct cgagcttttg gagtacgtcg tctttaggtt
5701 ggggggaggg gttttatgcg atggagtttc cccacactga gtgggtggag actgaagtta
5761 ggccagcttg gcacttgatg taattctcct tggaatttgc cctttttgag tttggatctt
5821 ggttcattct caagcctcag acagtggttc aaagtttttt tcttccattt caggtgtcgt
5881 gaagcggccg ccatgctagt aaattgtccg ctaaattctg gccgtttttg gcttttttgt
5941 tagacggatc cataacttcg tataatgtat gctatacgaa gttatgccac catgtctaga
6001 ctggacaaga gcaaagtcat aaacggagct ctggaattac tcaatggtgt cggtatcgaa
6061 ggcctgacga caaggaaact cgctcaaaag ctgggagttg agcagcctac cctgtactgg
6121 cacgtgaaga acaagcgggc cctgctcgat gccctgccaa tcgagatgct ggacaggcat
6181 catacccact tctgccccct ggaaggcgag tcatggcaag actttctgcg gaacaacgcc
6241 aagtcatacc gctgtgctct cctctcacat cgcgacgggg ctaaagtgca tctcggcacc
6301 cgcccaacag agaaacagta cgaaaccctg gaaaatcagc tcgcgttcct gtgtcagcaa
6361 ggcttctccc tggagaacgc actgtacgct ctgtccgccg tgggccactt tacactgggc
6421 tgcgtattgg aggaacagga gcatcaagta gcaaaagagg aaagagagac acctaccacc
6481 gattctatgc ccccacttct gagacaagca attgagctgt tcgaccggca gggagccgaa
6541 cctgccttcc ttttcggcct ggaactaatc atatgtggcc tggagaaaca gctaaagtgc
6601 gaaagcggcg ggccgaccga cgcccttgac gattttgact tagacatgct cccagccgat
6661 gcccttgacg actttgacct tgatatgctg cctgctgacg ctcttgacga ttttgacctt
6721 gacatgctcc ccgggtaact aagtaaggat ccgcggccgc actagaggaa ttccgcccct
6781 ctccccccccc ccctcccccc cccctaacgt tactggccga agccgcttgg aataaggccg
6841 gtgtgcgttt gtctatatgt tattttccac catattgccg tcttttggca atgtgagggc
6901 ccggaaacct ggccctgtct tcttgacgag cattcctagg ggtctttccc ctctcgccaa
6961 aggaatgcaa ggtctgttga atgtcgtgaa ggaagcagtt cctctggaag cttcttgaag
7021 acaaacaacg tctgtagcga ccctttgcag gcagcggaac cccccacctg gcgacaggtg
7081 cctctgcggc caaaagccac gtgtataaga tacacctgca aaggcggcac aaccccagtg
7141 ccacgttgtg agttggatag ttgtggaaag agtcaaatgg ctctcctcaa gcgtattcaa
7201 caaggggctg aaggatgccc agaaggtacc ccattgtatg ggatctgatc tggggcctcg
7261 gtgcacatgc tttacatgtg tttagtcgag gttaaaaaaa cgtctaggcc ccccgaacca
7321 cggggacgtg gttttccttt gaaaaacacg atgataagct taccggtcca ccatggtgag
7381 caagggcgag gagctgttca ccggggtggt gcccatcctg gtcgagctgg acggcgacgt
7441 aaacggccac aagttcagcg tgtccggcga gggcgagggc gatgccacct acggcaagct
7501 gaccctgaag ttcatctgca ccaccggcaa gctgcccgtg ccctggccca ccctcgtgac
7561 caccctgacc tacggcgtgc agtgcttcag ccgctacccc gaccacatga agcagcacga
7621 cttcttcaag tccgccatgc ccgaaggcta cgtccaggag cgcaccatct tcttcaagga
7681 cgacggcaac tacaagaccc gcgccgaggt gaagttcgag ggcgacaccc tggtgaaccg
7741 catcgagctg aagggcatcg acttcaagga ggacggcaac atcctggggc acaagctgga
7801 gtacaactac aacagccaca acgtctatat catggccgac aagcagaaga acggcatcaa
7861 ggtgaacttc aagatccgcc acaacatcga ggacggcagc gtgcagctcg ccgaccacta
7921 ccagcagaac acccccatcg gcgacggccc cgtgctgctg cccgacaacc actacctgag
7981 cacccagtcc gccctgagca aagaccccaa cgagaagcgc gatcacatgg tcctgctgga

8041 gttcgtgacc gccgccggga tcactctcgg catggacgag ctgtacaagt aagcatgccc
8101 gacggcgagg atctcgtcgt gacccatggc gatgcctgct tgccgaatat catggtggaa
8161 aatggccgct tttctggatt catcgactgt ggccggctgg gtgtggcgga ccgctatcag
8221 gacatagcgt tggctacccg tgatattgct gaagagcttg gcggcgaatg ggctgaccgc
8281 ttcctcgtgc tttacggtat cgccgctccc gattcgcagc gcatcgcctt ctatcgcctt
8341 cttgacgagt tcttctgatg tacaagtaaa gcggccgcga ctctagatca taatcagcca
8401 taccacattt gtagaggttt tacttgcttt aaaaaacctc ccacacctcc ccctgaacct
8461 gaaacataaa atgaatgcaa ttgttgttgt ttagtccctc ccaattcgat atcaagctta
8521 tcgcgataga tcctaatcaa cctctggatt acaaaatttg tgaaagattg actggtattc
8581 ttaactatgt tgctcctttt acgctatgtg gatacgctgc tttaatgcct ttgtatcatg
8641 ctattgcttc ccgtatggct ttcattttct cctccttgta taaatcctgg ttgctgtctc
8701 tttatgagga gttgtggccc gttgtcaggc aacgtggcgt ggtgtgcact gtgtttgctg
8761 acgcaacccc cactggttgg ggcattgcca ccacctgtca gctcctttcc gggactttcg
8821 ctttccccct ccctattgcc acggcggaac tcatcgccgc ctgccttgcc cgctgctgga
8881 caggggctcg gctgttgggc actgacaatt ccgtggtgtt gtcggggaaa tcatcgtcct
8941 ttccttggct gctcgcctgt gttgccacct ggattctgcg cgggacgtcc ttctgctacg
9001 tcccttcggc cctcaatcca gcggaccttc cttcccgcgg cctgctgccg gctctgcggc
9061 ctcttccgcg tcttcgcctt cgccctcaga cgagtcggat ctccctttgg gccgcctccc
9121 cgcctgagat cctttaagac caatgactta caaggcagct gtagatctta gccacttttt
9181 aaaagaaaag gggggactgg aagggctaat tcactcccaa cgaagacaag atctgctttt
9241 tgcttgtact gggtctctct ggttagacca gatctgagcc tgggagctct ctggctaact
9301 agggaaccca ctgcttaagc ctcaataaag cttgccttga gtgcttcaag tagtgtgtgc
9361 ccgtctgttg tgtgactctg gtaactagag atccctcaga cccttttagt cagtgtggaa
9421 aatctctagc agtagtagtt catgtcatct tattattcag tatttataac ttgcaaagaa
9481 atgaatatca gagagtgaga ggcccgggtt aattaaggaa agggctagat cattcttgaa
9541 gacgaaaggg cctcgtgata cgcctatttt tataggttaa tgtcatgata ataatggttt
9601 cttagacgtc aggtggcact tttcggggaa atgtgcgcgg aacccctatt tgtttatttt
9661 tctaaataca ttcaaatatg tatccgctca tgagacaata accctgataa atgcttcaat
9721 aatattgaaa aaggaagagt atgagtattc aacatttccg tgtcgccctt attccctttt
9781 ttgcggcatt ttgccttcct gtttttgctc acccagaaac gctggtgaaa gtaaaagatg
9841 ctgaagatca gttgggtgca cgagtgggtt acatcgaact ggatctcaac agcggtaaga
9901 tccttgagag ttttcgcccc gaagaacgtt ttccaatgat gagcactttt aaagttctgc
9961 tatgtggcgc ggtattatcc cgtgttgacg ccgggcaaga gcaactcggt cgccgcatac
10021 actattctca gaatgacttg gttgagtact caccagtcac agaaaagcat cttacggatg
10081 gcatgacagt aagagaatta tgcagtgctg ccataaccat gagtgataac actgcggcca
10141 acttacttct gacaacgatc ggaggaccga aggagctaac cgcttttttg cacaacatgg
10201 gggatcatgt aactcgcctt gatcgttggg aaccggagct gaatgaagcc ataccaaacg
10261 acgagcgtga caccacgatg cctgtagcaa tggcaacaac gttgcgcaaa ctattaactg
10321 gcgaactact tactctagct tcccggcaac aattaataga ctggatggag gcggataaag
10381 ttgcaggacc acttctgcgc tcggcccttc cggctggctg gtttattgct gataaatctg
10441 gagccggtga gcgtgggtct cgcggtatca ttgcagcact ggggccagat ggtaagccct
10501 cccgtatcgt agttatctac acgacgggga gtcaggcaac tatggatgaa cgaaatagac
10561 agatcgctga gataggtgcc tcactgatta agcattggta actgtcagac caagtttact
10621 catatatact ttagattgat ttaaaacttc atttttaatt taaaaggatc taggtgaaga
10681 tcctttttga taatctcatg accaaaatcc cttaacgtga gttttcgttc cactgagcgt
10741 cagaccccgt agaaaagatc aaaggatctt cttgagatcc tttttttctg cgcgtaatct
10801 gctgcttgca aacaaaaaaa ccaccgctac cagcggtggt ttgtttgccg gatcaagagc
10861 taccaactct ttttccgaag gtaactggct tcagcagagc gcagatacca aatactgttc
10921 ttctagtgta gccgtagtta ggccaccact tcaagaactc tgtagcaccg cctacatacc
10981 tcgctctgct aatcctgtta ccagtggctg ctgccagtgg cgataagtcg tgtcttaccg
11041 ggttggactc aagacgatag ttaccggata aggcgcagcg gtcgggctga acggggggtt
11101 cgtgcacaca gcccagcttg gagcgaacga cctacaccga actgagatac ctacagcgtg
11161 agctatgaga aagcgccacg cttcccgaag ggagaaaggc ggacaggtat ccggtaagcg
11221 gcagggtcgg aacaggagag cgcacgaggg agcttccagg gggaaacgcc tggtatcttt
11281 atagtcctgt cgggtttcgc cacctctgac ttgagcgtcg atttttgtga tgctcgtcag
11341 gggggcggag cctatggaaa aacgccagca acgcggcctt tttacggttc ctggcctttt
11401 gctggccttt tgctcacatg ttctttcctg cgttatcccc tgattctgtg gataaccgta
11461 ttaccgcctt tgagtgagct gataccgctc gccgcagccg aacgaccgag cgcagcgagt
11521 cagtgagcga ggaagcggaa gagcgcccaa tacgcaaacc gcctctcccc gcgcgttggc
11581 cgattcatta atgcagcaag ctcatggctg actaattttt tttatttatg cagaggccga
11641 ggccgcctcg gcctctgagc tattccagaa gtagtgagga ggcttttttg gaggcctagg
11701 cttttgcaaa aagctccccg tggcacgaca ggtttcccga ctggaaagcg ggcagtgagc
11761 gcaacgcaat taatgtgagt tagctcactc attaggcacc ccaggcttta cactttatgc
11821 ttccggctcg tatgttgtgt ggaattgtga gcggataaca atttcacaca ggaaacagct
11881 atgacatgat tacgaatttc acaaataaag catttttttc actgcattct agttgtggtt
11941 tgtccaaact catcaatgta tcttatcatg tctggatcaa ctggataact caagctaacc
12001 aaaatcatcc caaacttccc accccatacc ctattaccac tgccaattac ctgtggtttc

12061 atttactcta aacctgtgat tcctctgaat tattttcatt ttaaagaaat tgtatttgtt
12121 aaatatgtac tacaaactta gtagt

#### 2. pINDUCER21 (ORF-EG)_27675_WT

**Description** pINDUCER21 (ORF-EG) carrying wild-type PA14_27675 (dear) as the Gateway insert; doxycycline-inducible DeaR expression with constitutive IRES-EGFP.

**Length** 12,645 bp, circular

**Source** This paper

**Note** *Gateway cassette (attR1-CmR-ccdB-attR2, 1,705 bp of the parent) replaced by attB1-Kozak-PA14_27675-attB2 (2,198 bp): 12,145 - 1,705 + 2,198 + 7 = 12,645 bp. The 7 extra bp are a homopolymer-length difference from the parent in the poly-C tract between rtTA-Advanced and the IRES (non-coding spacer; see note on plasmid 3). The HA epitope at 4951..4977 is vector-derived and lies downstream of both attB2 and the PA14_27675 stop codon, so it is not translated: the expressed protein is untagged DeaR.*

##### Features

| Location | Type | Feature |
|---|---|---|
| 1..634 | LTR | 5' LTR (HIV-1) |
| 681..806 | misc_feature | HIV-1 Psi packaging signal |
| 1303..1536 | misc_feature | RRE (Rev response element) |
| 2058..2175 | misc_feature | cPPT/CTS |
| 2230..2500 | protein_bind | TRE (7x tet operator) |
| 2533..2571 | promoter | minimal CMV promoter |
| 2701..2725 | misc_recomb | attB1 |
| 2726..2734 | misc_feature | Kozak sequence |
| 2732..4873 | CDS | PA14_27675 (dear), wild type, 713 aa |
| 4874..4898 | misc_recomb | attB2 |
| 4951..4977 | misc_feature | HA epitope, vector-derived (not translated) |
| 5197..6375 | promoter | EF-1-alpha promoter |
| 5428..6366 | intron | EF-1-alpha intron A |
| 6445..6478 | protein_bind | loxP |
| 6479..6487 | misc_feature | Kozak sequence |
| 6485..7231 | CDS | rtTA-Advanced (248 aa) |
| 7395..7857 | misc_feature | IRES (EMCV) |
| 7873..8592 | CDS | EGFP (239 aa) |
| 9035..9623 | misc_feature | WPRE |
| 9698..9931 | LTR | 3' LTR (Delta-U3) |
| 10136..10240 | promoter | bla promoter |
| 10241..11101 | CDS | bla (ampicillin resistance, 286 aa) |
| 11272..11860 | rep_origin | ori (pUC/ColE1) |
| 12307..12337 | promoter | lac promoter |
| 12345..12361 | protein_bind | lac operator |

##### Sequence (12,645 bp)

```
1 tggaagggct aattcactcc caaagaagac aagatatcct tgatctgtgg atctaccaca
```

61 cacaaggcta cttccctgat tagcagaact acacaccagg gccaggggtc agatatccac
121 tgacctttgg atggtgctac aagctagtac cagttgagcc agataaggta gaagaggcca
181 ataaaggaga gaacaccagc ttgttacacc ctgtgagcct gcatgggatg gatgacccgg
241 agagagaagt gttagagtgg aggtttgaca gccgcctagc atttcatcac gtggcccgag
301 agctgcatcc ggagtacttc aagaactgct gatatcgagc ttgctacaag ggactttccg
361 ctggggactt tccagggagg cgtggcctgg gcgggactgg ggagtggcga gccctcagat
421 cctgcatata agcagctgct ttttgcctgt actgggtctc tctggttaga ccagatctga
481 gcctgggagc tctctggcta actagggaac ccactgctta agcctcaata aagcttgcct
541 tgagtgcttc aagtagtgtg tgcccgtctg ttgtgtgact ctggtaacta gagatccctc
601 agaccctttt agtcagtgtg gaaaatctct agcagtggcg cccgaacagg gacttgaaag
661 cgaaagggaa accagaggag ctctctcgac gcaggactcg gcttgctgaa gcgcgcacgg
721 caagaggcga ggggcggcga ctggtgagta cgccaaaaat tttgactagc ggaggctaga
781 aggagagaga tgggtgcgag agcgtcagta ttaagcgggg gagaattaga tcgcgatggg
841 aaaaaattcg gttaaggcca gggggaaaga aaaaatataa attaaaacat atagtatggg
901 caagcaggga gctagaacga ttcgcagtta atcctggcct gttagaaaca tcagaaggct
961 gtagacaaat actgggacag ctacaaccat cccttcagac aggatcagaa gaacttagat
1021 cattatataa tacagtagca accctctatt gtgtgcatca aaggatagag ataaaagaca
1081 ccaaggaagc tttagacaag atagaggaag agcaaaacaa aagtaagacc accgcacagc
1141 aagcggccgg ccgctgatct tcagacctgg aggaggagat atgagggaca attggagaag
1201 tgaattatat aaatataaag tagtaaaaat tgaaccatta ggagtagcac ccaccaaggc
1261 aaagagaaga gtggtgcaga gagaaaaaag agcagtggga ataggagctt tgttccttgg
1321 gttcttggga gcagcaggaa gcactatggg cgcagcgtca atgacgctga cggtacaggc
1381 cagacaatta ttgtctggta tagtgcagca gcagaacaat ttgctgaggg ctattgaggc
1441 gcaacagcat ctgttgcaac tcacagtctg gggcatcaag cagctccagg caagaatcct
1501 ggctgtggaa agatacctaa aggatcaaca gctcctgggg atttggggtt gctctggaaa
1561 actcatttgc accactgctg tgccttggaa tgctagttgg agtaataaat ctctggaaca
1621 gatttggaat cacacgacct ggatggagtg ggacagagaa attaacaatt acacaagctt
1681 aatacactcc ttaattgaag aatcgcaaaa ccagcaagaa aagaatgaac aagaattatt
1741 ggaattagat aaatgggcaa gtttgtggaa ttggtttaac ataacaaatt ggctgtggta
1801 tataaaatta ttcataatga tagtaggagg cttggtaggt ttaagaatag tttttgctgt
1861 actttctata gtgaatagag ttaggcaggg atattcacca ttatcgtttc agacccacct
1921 cccaaccccg aggggacccg acaggcccga aggaatagaa gaagaaggtg gagagagaga
1981 cagagacaga tccattcgat tagtgaacgg atctcgacgg tatcgccgaa ttcacaaatg
2041 gcagtattca tccacaattt taaaagaaaa ggggggattg gggggtacag tgcaggggaa
2101 agaatagtag acataatagc aacagacata caaactaaag aattacaaaa acaaattaca
2161 aaaattcaaa attttcgggt ttattacagg gacagcagag atccagtttg gactaggatc
2221 ctttaccact ccctatcagt gatagagaaa agtgaaagtc gagtttacca ctccctatca
2281 gtgatagaga aaagtgaaag tcgagtttac cactccctat cagtgataga gaaaagtgaa
2341 agtcgagttt accactccct atcagtgata gagaaaagtg aaagtcgagt ttaccactcc
2401 ctatcagtga tagagaaaag tgaaagtcga gtttaccact ccctatcagt gatagagaaa
2461 agtgaaagtc gagtttacca ctccctatca gtgatagaga aaagtgaaag tcgagctcgg
2521 tacccgggtc gaggtaggcg tgtacggtgg gaggcctata taagcagagc tcgtttagtg
2581 aaccgtcaga tcgcctggag acgccatcca cgctgttttg acctccatag aagacaccgg
2641 gaccgatcca gcctccgcgg ccccgaacta gtccagtgtg gtggaattct gcagatatca
2701 acaagtttgt acaaaaaagc aggctgccac catggcttat ttcaaacttg aagaacccgt
2761 ccgatttcac cgatatccct ttgatttcca ctcccatttt gcgggaatct tgccagtcga
2821 atctaatagc cggtggacta gagatcgaag ggtgtttcga gtcggcgagc gccaagtatc
2881 tctcgagaaa ggtcaagagt tgagtctgat tggcctgctg atgagtgccc ggggtgtggc
2941 tcccgatgtt gacggcaagg cgctcgaaga ggcacgccaa gcggctcact atgagctttt
3001 cgaactggcg ttgcagcgaa tggttcggcg aaatccgttt gctgctactg accgccaagg
3061 ctatctgagg ggtgaatgtg ctgccgaaaa tatctacctt gcatgcctta tactggccca
3121 gcgcttcggt cgcacatcac cgcctgctgc gatcgaccaa ccagcgatat atcttgggac
3181 attggagctt cttggggcga gtgccgtgcg ggattccgaa actgaccagt tcgttcgcta
3241 ctttaatcgc aaaatttggt ccggcaataa atacacccct tttgacgacg catactgggc
3301 tcgcggggcc ataagggatc ggcaccctgg ggaattcgcc tgtttgactt tgggtttctt
3361 gctccatgag ggatctcac atacccagac cgcaaccggg gaggatgagg ttgcggttct
3421 tgactccctt ttcgaacaat ttaatgcgag tgaaaaaact gcttatcgat tgttggcaca
3481 tacggcacat ggctatgcat cagaagcggc atttgatgct gaactccacc gcatacttcg
3541 gcattttgag atccaacagg gacagcctcc acaagccaga cttgtaggta ttgatctcct
3601 gggcatggaa accgccactg gactgtatcg ccagtttttt gactttcttc tcgggcaggc
3661 ggctgtcttt cggcactatt tggatgggaa accagaaaca cgcaagatgg tgcttcacat
3721 acattgcgga gaaggaacgg gagtgagtga cgacaatagg tctctttgtg gctacttcct
3781 cagaaatgct aacgccctcg acgactttta tgcggcactc tctgcctacg cctggaagtg
3841 ctacgggaat actatccgac agggtaaagc acgcttgcgc gaacgagaga atcttcagga
3901 acgcgacaag gccccttcag cacttgctgg tctcttcgac gaacttttct tcggtaacag
3961 ccttaccgct agcggcctgc gcctccgcag attcgacata acatctggca caactcaggc
4021 tctcgttgca tattacgcaa ggacgaacgt cgtaaatctt tgccaagcac ttgccagtag

4081 agatgccgac ggaaactcct actaccggcg ccttctcgaa tcagatctct tttcccttcg
4141 gataggtcac gcctactatt accggaacta tctcgcttca aagttccccg aactctgttt
4201 tgacaccaac ctcgggtcta actttattac tggcgcttcc ggactgttcg atagtctgca
4261 agaatatcgc ttgaaccggg ggctcaggca tttggacggt tatgtgggga cagatcaatt
4321 gaaggagctg tcattggcca tcgcttacca gggagaacag cgactggacc ctcaacagat
4381 gcaatatgtt catgcgctgg cagagagcca gtcaggcttc gatgagctcg gtgagcacct
4441 gccaggcaca cccggctggg cgaaaccagc actggaacaa tttttgtta gtcagtgtgc
4501 actctatagg tctgaagagg accgctattt tcaattcgaa gcctaccggc gcttgtttgc
4561 gcaggtattg aactggcgat cttaccttct tggggcagac ggacaaggtg tagaacacag
4621 taatgtccag gacgaggcga tacggatggc ccttctgttg aattatgcag cggcggaccg
4681 ccatggccga gttcccgccg cgtccctgga aaatgcgcag cgcctcttgg tccaactcgg
4741 aagcgcctac tgggaagaga cgattggtgc tgtagacctc gcaggagctc ctcatcgaga
4801 ccgagaactc caacgcttcg aaggttttgc ggcccctgca agtgttgttc gaatctcaac
4861 tagatcttcc tgaacccagc tttcttgtac aaagtggttg atatccagca cagtggcggc
4921 cgctcgagtc tagagggccc gcggttcgaa tacccatacg acgtcccaga ctacgcttag
4981 taatgattaa ttaaactaga aattctaccg ggtaggggag gcgcttttcc caaggcagtc
5041 tggagtaacc cacccaagat ctggcctccg cgccgggttt tggcgcctcc cgcgggcgcc
5101 cccctcctca cggcgagccg cgtaagatct ggcctccgcg ccgggttttg gcgcctcccg
5161 cgggcgcccc cctcctcacg gcgagcctag tcgtgaggct ccggtgcccg tcagtgggca
5221 gagcgcacat cgcccacagt ccccgagaag ttggggggag gggtcggcaa ttgaaccggt
5281 gcctagagaa ggtggcgcgg ggtaaactgg gaaagtgatg tcgtgtactg gctccgcctt
5341 tttcccgagg gtgggggaga accgtatata agtgcagtag tcgccgtgaa cgttcttttt
5401 cgcaacgggt ttgccgccag aacacaggta agtgccgtgt gtggttcccg cgggcctggc
5461 ctctttacgg gttatggccc ttgcgtgcct tgaattactt ccacctggct gcagtacgtg
5521 attcttgatc ccgagcttcg ggttggaagt gggtgggaga gttcgaggcc ttgcgcttaa
5581 ggagcccctt cgcctcgtgc ttgagttgag gcctggcctg ggcgctgggg ccgccgcgtg
5641 cgaatctggt ggcaccttcg cgcctgtctc gctgctttcg ataagtctct agccatttaa
5701 aatttttgat gacctgctgc gacgcttttt ttctggcaag atagtcttgt aaatgcgggc
5761 caagatctgc acactggtat ttcggttttt ggggccgcgg gcggcgacgg ggcccgtgcg
5821 tcccagcgca catgttcggc gaggcggggc ctgcgagcgc ggccaccgag aatcggacgg
5881 gggtagtctc aagctggccg gcctgctctg gtgcctggcc tcgcgccgcc gtgtatcgcc
5941 ccgccctggg cggcaaggct ggcccggtcg gcaccagttg cgtgagcgga aagatggccg
6001 cttcccggcc ctgctgcagg gagctcaaaa tggaggacgc ggcgctcggg agagcgggcg
6061 ggtgagtcac ccacacaaag gaaaagggcc tttccgtcct cagccgtcgc ttcatgtgac
6121 tccacggagt accgggcgcc gtccaggcac ctcgattagt tctcgagctt ttggagtacg
6181 tcgtctttag gttgggggga ggggttttat gcgatggagt ttccccacac tgagtgggtg
6241 gagactgaag ttaggccagc ttggcacttg atgtaattct ccttggaatt tgcccttttt
6301 gagtttggat cttggttcat tctcaagcct cagacagtgg ttcaaagttt ttttcttcca
6361 tttcaggtgt cgtgaagcgg ccgccatgct agtaaattgt ccgctaaatt ctggccgttt
6421 ttggcttttt tgttagacgg atccataact tcgtataatg tatgctatac gaagttatgc
6481 caccatgtct agactggaca agagcaaagt cataaacgga gctctggaat tactcaatgg
6541 tgtcggtatc gaaggcctga cgacaaggaa actcgctcaa aagctgggag ttgagcagcc
6601 taccctgtac tggcacgtga agaacaagcg ggccctgctc gatgccctgc caatcgagat
6661 gctggacagg catcataccc acttctgccc cctggaaggc gagtcatggc aagactttct
6721 gcggaacaac gccaagtcat accgctgtgc tctcctctca catcgcgacg ggggctaaagt
6781 gcatctcggc acccgcccaa cagagaaaca gtacgaaacc ctggaaaatc agctcgcgtt
6841 cctgtgtcag caaggcttct ccctggagaa cgcactgtac gctctgtccg ccgtgggcca
6901 ctttacactg ggctgcgtat tggaggaaca ggagcatcaa gtagcaaaag aggaaagaga
6961 gacacctacc accgattcta tgcccccact tctgagacaa gcaattgagc tgttcgaccg
7021 gcagggagcc gaacctgcct tccttttcgg cctggaacta atcatatgtg gcctggagaa
7081 acagctaaag tgcgaaagcg gcgggccgac cgacgccctt gacgattttg acttagacat
7141 gctcccagcc gatgcccttg acgactttga ccttgatatg ctgcctgctg acgctcttga
7201 cgattttgac cttgacatgc tccccgggta actaagtaag gatccgcggc cgcactagag
7261 gaattccgcc cctctccccc ccccccctct ccctcccccc cccctaacgt tactggccga
7321 agccgcttgg aataaggccg gtgtgcgttt gtctatatgt tattttccac catattgccg
7381 tcttttggca atgtgagggc ccggaaacct ggccctgtct tcttgacgag cattcctagg
7441 ggtctttccc ctctcgccaa aggaatgcaa ggtctgttga atgtcgtgaa ggaagcagtt
7501 cctctggaag cttcttgaag acaaacaacg tctgtagcga ccctttgcag gcagcggaac
7561 cccccacctg gcgacaggtg cctctgcggc caaaagccac gtgtataaga tacacctgca
7621 aaggcggcac aaccccagtg ccacgttgtg agttggatag ttgtggaaag agtcaaatgg
7681 ctctcctcaa gcgtattcaa caaggggctg aaggatgccc agaaggtacc ccattgtatg
7741 ggatctgatc tggggcctcg gtgcacatgc tttacatgtg tttagtcgag gttaaaaaaa
7801 cgtctaggcc ccccgaacca cggggacgtg gttttccttt gaaaaacacg atgataagct
7861 taccggtcca ccatggtgag caagggcgag gagctgttca ccggggtggt gcccatcctg
7921 gtcgagctgg acggcgacgt aaacggccac aagttcagcg tgtccggcga gggcgagggc
7981 gatgccacct acggcaagct gaccctgaag ttcatctgca ccaccggcaa gctgcccgtg
8041 ccctggccca ccctcgtgac caccctgacc tacggcgtgc agtgcttcag ccgctacccc

8101 gaccacatga agcagcacga cttcttcaag tccgccatgc ccgaaggcta cgtccaggag
8161 cgcaccatct tcttcaagga cgacggcaac tacaagaccc gcgccgaggt gaagttcgag
8221 ggcgacaccc tggtgaaccg catcgagctg aagggcatcg acttcaagga ggacggcaac
8281 atcctggggc acaagctgga gtacaactac aacagccaca acgtctatat catggccgac
8341 aagcagaaga acggcatcaa ggtgaacttc aagatccgcc acaacatcga ggacggcagc
8401 gtgcagctcg ccgaccacta ccagcagaac acccccatcg gcgacggccc cgtgctgctg
8461 cccgacaacc actacctgag cacccagtcc gccctgagca aagaccccaa cgagaagcgc
8521 gatcacatgg tcctgctgga gttcgtgacc gccgccggga tcactctcgg catggacgag
8581 ctgtacaagt aagcatgccc gacggcgagg atctcgtcgt gacccatggc gatgcctgct
8641 tgccgaatat catggtggaa aatggccgct tttctggatt catcgactgt ggccggctgg
8701 gtgtggcgga ccgctatcag gacatagcgt tggctacccg tgatattgct gaagagcttg
8761 gcggcgaatg ggctgaccgc ttcctcgtgc tttacggtat cgccgctccc gattcgcagc
8821 gcatcgcctt ctatcgcctt cttgacgagt tcttctgatg tacaagtaaa gcggccgcga
8881 ctctagatca taatcagcca taccacattt gtagaggttt tacttgcttt aaaaaacctc
8941 ccacacctcc ccctgaacct gaaacataaa atgaatgcaa ttgttgttgt ttagtccctc
9001 ccaattcgat atcaagctta tcgcgataga tcctaatcaa cctctggatt acaaaatttg
9061 tgaaagattg actggtattc ttaactatgt tgctcctttt acgctatgtg gatacgctgc
9121 tttaatgcct ttgtatcatg ctattgcttc ccgtatggct ttcattttct cctccttgta
9181 taaatcctgg ttgctgtctc tttatgagga gttgtggccc gttgtcaggc aacgtggcgt
9241 ggtgtgcact gtgtttgctg acgcaacccc cactggttgg ggcattgcca ccacctgtca
9301 gctcctttcc gggactttcg ctttccccct ccctattgcc acggcggaac tcatcgccgc
9361 ctgccttgcc cgctgctgga caggggctcg gctgttgggc actgacaatt ccgtggtgtt
9421 gtcggggaaa tcatcgtcct ttccttggct gctcgcctgt gttgccacct ggattctgcg
9481 cgggacgtcc ttctgctacg tcccttcggc cctcaatcca gcggaccttc cttcccgcgg
9541 cctgctgccg gctctgcggc ctcttccgcg tcttcgcctt cgccctcaga cgagtcggat
9601 ctccctttgg gccgcctccc cgcctgagat cctttaagac caatgactta caaggcagct
9661 gtagatctta gccacttttt aaaagaaaag gggggactgg aagggctaat tcactcccaa
9721 cgaagacaag atctgctttt tgcttgtact gggtctctct ggttagacca gatctgagcc
9781 tgggagctct ctggctaact agggaaccca ctgcttaagc ctcaataaag cttgccttga
9841 gtgcttcaag tagtgtgtgc ccgtctgttg tgtgactctg gtaactagag atccctcaga
9901 cccttttagt cagtgtggaa aatctctagc agtagtagtt catgtcatct tattattcag
9961 tatttataac ttgcaaagaa atgaatatca gagagtgaga ggcccgggtt aattaaggaa
10021 agggctagat cattcttgaa gacgaaaggg cctcgtgata cgcctatttt tataggttaa
10081 tgtcatgata ataatggttt cttagacgtc aggtggcact tttcggggaa atgtgcgcgg
10141 aacccctatt tgtttatttt tctaaataca ttcaaatatg tatccgctca tgagacaata
10201 accctgataa atgcttcaat aatattgaaa aaggaagagt atgagtattc aacatttccg
10261 tgtcgccctt attccctttt ttgcggcatt ttgccttcct gtttttgctc acccagaaac
10321 gctggtgaaa gtaaaagatg ctgaagatca gttgggtgca cgagtgggtt acatcgaact
10381 ggatctcaac agcggtaaga tccttgagag ttttcgcccc gaagaacgtt ttccaatgat
10441 gagcactttt aaagttctgc tatgtggcgc ggtattatcc cgtgttgacg ccgggcaaga
10501 gcaactcggt cgccgcatac actattctca gaatgacttg gttgagtact caccagtcac
10561 agaaaagcat cttacggatg gcatgacagt aagagaatta tgcagtgctg ccataaccat
10621 gagtgataac actgcggcca acttacttct gacaacgatc ggaggaccga aggagctaac
10681 cgcttttttg cacaacatgg gggatcatgt aactcgcctt gatcgttggg aaccggagct
10741 gaatgaagcc ataccaaacg acgagcgtga caccacgatg cctgtagcaa tggcaacaac
10801 gttgcgcaaa ctattaactg gcgaactact tactctagct tcccggcaac aattaataga
10861 ctggatggag gcggataaag ttgcaggacc acttctgcgc tcggcccttc cggctggctg
10921 gtttattgct gataaatctg gagccggtga gcgtgggtct cgcggtatca ttgcagcact
10981 ggggccagat ggtaagccct cccgtatcgt agttatctac acgacgggga gtcaggcaac
11041 tatggatgaa cgaaatagac agatcgctga gataggtgcc tcactgatta agcattggta
11101 actgtcagac caagtttact catatatact ttagattgat ttaaaacttc atttttaatt
11161 taaaaggatc taggtgaaga tcctttttga taatctcatg accaaaatcc cttaacgtga
11221 gttttcgttc cactgagcgt cagaccccgt agaaaagatc aaaggatctt cttgagatcc
11281 tttttttctg cgcgtaatct gctgcttgca aacaaaaaaa ccaccgctac cagcggtggt
11341 ttgtttgccg gatcaagagc taccaactct ttttccgaag gtaactggct tcagcagagc
11401 gcagatacca aatactgttc ttctagtgta gccgtagtta ggccaccact tcaagaactc
11461 tgtagcaccg cctacatacc tcgctctgct aatcctgtta ccagtggctg ctgccagtgg
11521 cgataagtcg tgtcttaccg ggttggactc aagacgatag ttaccggata aggcgcagcg
11581 gtcgggctga acggggggtt cgtgcacaca gcccagcttg gagcgaacga cctacaccga
11641 actgagatac ctacagcgtg agctatgaga aagcgccacg cttcccgaag ggagaaaggc
11701 ggacaggtat ccggtaagcg gcagggtcgg aacaggagag cgcacgaggg agcttccagg
11761 gggaaacgcc tggtatcttt atagtcctgt cgggtttcgc cacctctgac ttgagcgtcg
11821 atttttgtga tgctcgtcag gggggcggag cctatggaaa aacgccagca acgcggcctt
11881 tttacggttc ctggcctttt gctggccttt tgctcacatg ttctttcctg cgttatcccc
11941 tgattctgtg gataaccgta ttaccgcctt tgagtgagct gataccgctc gccgcagccg
12001 aacgaccgag cgcagcgagt cagtgagcga ggaagcggaa gagcgcccaa tacgcaaacc
12061 gcctctcccc gcgcgttggc cgattcatta atgcagcaag ctcatggctg actaattttt

12121 tttatttatg cagaggccga ggccgcctcg gcctctgagc tattccagaa gtagtgagga
12181 ggcttttttg gaggcctagg cttttgcaaa aagctccccg tggcacgaca ggtttcccga
12241 ctggaaagcg ggcagtgagc gcaacgcaat taatgtgagt tagctcactc attaggcacc
12301 ccaggcttta cactttatgc ttccggctcg tatgttgtgt ggaattgtga gcggataaca
12361 atttcacaca ggaaacagct atgacatgat tacgaatttc acaaataaag catttttttc
12421 actgcattct agttgtggtt tgtccaaact catcaatgta tcttatcatg tctggatcaa
12481 ctggataact caagctaacc aaaatcatcc caaacttccc accccatacc ctattaccac
12541 tgccaattac ctgtggtttc atttactcta aacctgtgat tcctctgaat tattttcatt
12601 ttaaagaaat tgtatttgtt aaatatgtac tacaaactta gtagt

#### 3. pINDUCER21 (ORF-EG)_27675_H22A

**Description** As plasmid 2, carrying the active-site PA14_27675 H22A allele (codon 22 CAT->GCC).

**Length** 12,646 bp, circular

**Source** This paper

**Note** *Differs from plasmid 2 at exactly two positions: the intended H22A codon (2795-2797) and one additional C in the poly-C tract at 7295 (11 C vs 10 C), a non-coding spacer between rtTA-Advanced and the IRES. The latter is a homopolymer- length call, not an engineered change; it shifts all coordinates at and after 7295 by +1 relative to plasmid 2. As in plasmid 2, the HA epitope at 4951..4977 is vector-derived and not translated.*

##### Features

| Location | Type | Feature |
|---|---|---|
| 1..634 | LTR | 5' LTR (HIV-1) |
| 681..806 | misc_feature | HIV-1 Psi packaging signal |
| 1303..1536 | misc_feature | RRE (Rev response element) |
| 2058..2175 | misc_feature | cPPT/CTS |
| 2230..2500 | protein_bind | TRE (7x tet operator) |
| 2533..2571 | promoter | minimal CMV promoter |
| 2701..2725 | misc_recomb | attB1 |
| 2726..2734 | misc_feature | Kozak sequence |
| 2732..4873 | CDS | PA14_27675 (dear), H22A, 713 aa |
| 4874..4898 | misc_recomb | attB2 |
| 4951..4977 | misc_feature | HA epitope, vector-derived (not translated) |
| 5197..6375 | promoter | EF-1-alpha promoter |
| 5428..6366 | intron | EF-1-alpha intron A |
| 6445..6478 | protein_bind | loxP |
| 6479..6487 | misc_feature | Kozak sequence |
| 6485..7231 | CDS | rtTA-Advanced (248 aa) |
| 7396..7858 | misc_feature | IRES (EMCV) |
| 7874..8593 | CDS | EGFP (239 aa) |
| 9036..9624 | misc_feature | WPRE |
| 9699..9932 | LTR | 3' LTR (Delta-U3) |
| 10137..10241 | promoter | bla promoter |
| 10242..11102 | CDS | bla (ampicillin resistance, 286 aa) |
| 11273..11861 | rep_origin | ori (pUC/ColE1) |
| 12308..12338 | promoter | lac promoter |
| 12346..12362 | protein_bind | lac operator |

##### Sequence (12,646 bp)

```
  1 tggaagggct aattcactcc caaagaagac aagatatcct tgatctgtgg atctaccaca
 61 cacaaggcta cttccctgat tagcagaact acacaccagg gccaggggtc agatatccac
121 tgacctttgg atggtgctac aagctagtac cagttgagcc agataaggta gaagaggcca
181 ataaaggaga gaacaccagc ttgttacacc ctgtgagcct gcatgggatg gatgacccgg
```

241 agagagaagt gttagagtgg aggtttgaca gccgcctagc atttcatcac gtggcccgag
301 agctgcatcc ggagtacttc aagaactgct gatatcgagc ttgctacaag ggactttccg
361 ctggggactt tccagggagg cgtggcctgg gcgggactgg ggagtggcga gccctcagat
421 cctgcatata agcagctgct ttttgcctgt actgggtctc tctggttaga ccagatctga
481 gcctgggagc tctctggcta actagggaac ccactgctta agcctcaata aagcttgcct
541 tgagtgcttc aagtagtgtg tgcccgtctg ttgtgtgact ctggtaacta gagatccctc
601 agaccctttt agtcagtgtg gaaaatctct agcagtggcg cccgaacagg gacttgaaag
661 cgaaagggaa accagaggag ctctctcgac gcaggactcg gcttgctgaa gcgcgcacgg
721 caagaggcga ggggcggcga ctggtgagta cgccaaaaat tttgactagc ggaggctaga
781 aggagagaga tgggtgcgag agcgtcagta ttaagcgggg gagaattaga tcgcgatggg
841 aaaaaattcg gttaaggcca gggggaaaga aaaaatataa attaaaacat atagtatggg
901 caagcaggga gctagaacga ttcgcagtta atcctggcct gttagaaaca tcagaaggct
961 gtagacaaat actgggacag ctacaaccat cccttcagac aggatcagaa gaacttagat
1021 cattatataa tacagtagca accctctatt gtgtgcatca aaggatagag ataaaagaca
1081 ccaaggaagc tttagacaag atagaggaag agcaaaacaa aagtaagacc accgcacagc
1141 aagcggccgg ccgctgatct tcagacctgg aggaggagat atgagggaca attggagaag
1201 tgaattatat aaatataaag tagtaaaaat tgaaccatta ggagtagcac ccaccaaggc
1261 aaagagaaga gtggtgcaga gagaaaaaag agcagtggga ataggagctt tgttccttgg
1321 gttcttggga gcagcaggaa gcactatggg cgcagcgtca atgacgctga cggtacaggc
1381 cagacaatta ttgtctggta tagtgcagca gcagaacaat ttgctgaggg ctattgaggc
1441 gcaacagcat ctgttgcaac tcacagtctg gggcatcaag cagctccagg caagaatcct
1501 ggctgtggaa agatacctaa aggatcaaca gctcctgggg atttggggtt gctctggaaa
1561 actcatttgc accactgctg tgccttggaa tgctagttgg agtaataaat ctctggaaca
1621 gatttggaat cacacgacct ggatggagtg ggacagagaa attaacaatt acacaagctt
1681 aatacactcc ttaattgaag aatcgcaaaa ccagcaagaa aagaatgaac aagaattatt
1741 ggaattagat aaatgggcaa gtttgtggaa ttggtttaac ataacaaatt ggctgtggta
1801 tataaaatta ttcataatga tagtaggagg cttggtaggt ttaagaatag tttttgctgt
1861 actttctata gtgaatagag ttaggcaggg atattcacca ttatcgtttc agacccacct
1921 cccaaccccg aggggacccg acaggcccga aggaatagaa gaagaaggtg gagagagaga
1981 cagagacaga tccattcgat tagtgaacgg atctcgacgg tatcgccgaa ttcacaaatg
2041 gcagtattca tccacaattt taaaagaaaa ggggggattg gggggtacag tgcaggggaa
2101 agaatagtag acataatagc aacagacata caaactaaag aattacaaaa acaaattaca
2161 aaaattcaaa attttcgggt ttattacagg gacagcagag atccagtttg gactaggatc
2221 ctttaccact ccctatcagt gatagagaaa agtgaaagtc gagtttacca ctccctatca
2281 gtgatagaga aaagtgaaag tcgagtttac cactccctat cagtgataga gaaaagtgaa
2341 agtcgagttt accactccct atcagtgata gagaaaagtg aaagtcgagt ttaccactcc
2401 ctatcagtga tagagaaaag tgaaagtcga gtttaccact ccctatcagt gatagagaaa
2461 agtgaaagtc gagtttacca ctccctatca gtgatagaga aaagtgaaag tcgagctcgg
2521 tacccgggtc gaggtaggcg tgtacggtgg gaggcctata taagcagagc tcgtttagtg
2581 aaccgtcaga tcgcctggag acgccatcca cgctgttttg acctccatag aagacaccgg
2641 gaccgatcca gcctccgcgg ccccgaacta gtccagtgtg gtggaattct gcagatatca
2701 acaagtttgt acaaaaaagc aggctgccac catggcttat ttcaaacttg aagaacccgt
2761 ccgatttcac cgatatccct ttgatttcca ctccgccttt gcgggaatct tgccagtcga
2821 atctaatagc cggtggacta gagatcgaag ggtgtttcga gtcggcgagc gccaagtatc
2881 tctcgagaaa ggtcaagagt tgagtctgat tggcctgctg atgagtgccc ggggtgtggc
2941 tcccgatgtt gacggcaagg cgctcgaaga ggcacgccaa gcggctcact atgagctttt
3001 cgaactggcg ttgcagcgaa tggttcggcg aaatccgttt gctgctactg accgccaagg
3061 ctatctgagg ggtgaatgtg ctgccgaaaa tatctacctt gcatgcctta tactggccca
3121 gcgcttcggt cgcacatcac cgcctgctgc gatcgaccaa ccagcgatat atcttgggac
3181 attggagctt cttggggcga gtgccgtgcg ggattccgaa actgaccagt tcgttcgcta
3241 ctttaatcgc aaaatttggt ccggcaataa atacacccct tttgacgacg catactgggc
3301 tcgcggggcc ataagggatc ggcaccctgg ggaattcgcc tgtttgactt tgggtttctt
3361 gctccatgag ggatctcac atacccagac cgcaaccggg gaggatgagg ttgcggttct
3421 tgactccctt ttcgaacaat ttaatgcgag tgaaaaaact gcttatcgat tgttggcaca
3481 tacggcacat ggctatgcat cagaagcggc atttgatgct gaactccacc gcatacttcg
3541 gcattttgag atccaacagg gacagcctcc acaagccaga cttgtaggta ttgatctcct
3601 gggcatggaa accgccactg gactgtatcg ccagtttttt gactttcttc tcgggcaggc
3661 ggctgtcttt cggcactatt tggatgggaa accagaaaca cgcaagatgg tgcttcacat
3721 acattgcgga gaaggaacgg gagtgagtga cgacaatagg tctctttgtg gctacttcct
3781 cagaaatgct aacgccctcg acgactttta tgcggcactc tctgcctacg cctggaagtg
3841 ctacgggaat actatccgac agggtaaagc acgcttgcgc gaacgagaga atcttcagga
3901 acgcgacaag gccccttcag cacttgctgg tctcttcgac gaacttttct tcggtaacag
3961 ccttaccgct agcggcctgc gcctccgcag attcgacata acatctggca caactcaggc
4021 tctcgttgca tattacgcaa ggacgaacgt cgtaaatctt tgccaagcac ttgccagtag
4081 agatgccgac ggaaactcct actaccggcg ccttctcgaa tcagatctct tttcccttcg
4141 gataggtcac gcctactatt accggaacta tctcgcttca aagttccccg aactctgttt
4201 tgacaccaac ctcgggtcta actttattac tggcgcttcc ggactgttcg atagtctgca

4261 agaatatcgc ttgaaccggg ggctcaggca tttggacggt tatgtgggga cagatcaatt
4321 gaaggagctg tcattggcca tcgcttacca gggagaacag cgactggacc ctcaacagat
4381 gcaatatgtt catgcgctgg cagagagcca gtcaggcttc gatgagctcg gtgagcacct
4441 gccaggcaca cccggctggg cgaaaccagc actggaacaa ttttttgtta gtcagtgtgc
4501 actctatagg tctgaagagg accgctattt tcaattcgaa gcctaccggc gcttgtttgc
4561 gcaggtattg aactggcgat cttaccttct tggggcagac ggacaaggtg tagaacacag
4621 taatgtccag gacgaggcga tacggatggc ccttctgttg aattatgcag cggcggaccg
4681 ccatggccga gttcccgccg cgtccctgga aaatgcgcag cgcctcttgg tccaactcgg
4741 aagcgcctac tgggaagaga cgattggtgc tgtagacctc gcaggagctc ctcatcgaga
4801 ccgagaactc caacgcttcg aaggttttgc ggcccctgca agtgttgttc gaatctcaac
4861 tagatcttcc tgaacccagc tttcttgtac aaagtggttg atatccagca cagtggcggc
4921 cgctcgagtc tagagggccc gcggttcgaa tacccatacg acgtcccaga ctacgcttag
4981 taatgattaa ttaaactaga aattctaccg ggtaggggag gcgcttttcc caaggcagtc
5041 tggagtaacc cacccaagat ctggcctccg cgccgggttt tggcgcctcc cgcgggcgcc
5101 cccctcctca cggcgagccg cgtaagatct ggcctccgcg ccgggttttg gcgcctcccg
5161 cgggcgcccc cctcctcacg gcgagcctag tcgtgaggct ccggtgcccg tcagtgggca
5221 gagcgcacat cgcccacagt ccccgagaag ttggggggag gggtcggcaa ttgaaccggt
5281 gcctagagaa ggtggcgcgg ggtaaactgg gaaagtgatg tcgtgtactg gctccgcctt
5341 tttcccgagg gtgggggaga accgtatata agtgcagtag tcgccgtgaa cgttcttttt
5401 cgcaacgggt ttgccgccag aacacaggta agtgccgtgt gtggttcccg cgggcctggc
5461 ctctttacgg gttatggccc ttgcgtgcct tgaattactt ccacctggct gcagtacgtg
5521 attcttgatc ccgagcttcg ggttggaagt gggtgggaga gttcgaggcc ttgcgcttaa
5581 ggagcccctt cgcctcgtgc ttgagttgag gcctggcctg ggcgctgggg ccgccgcgtg
5641 cgaatctggt ggcaccttcg cgcctgtctc gctgctttcg ataagtctct agccatttaa
5701 aatttttgat gacctgctgc gacgcttttt ttctggcaag atagtcttgt aaatgcgggc
5761 caagatctgc acactggtat ttcggttttt ggggccgcgg gcggcgacgg ggcccgtgcg
5821 tcccagcgca catgttcggc gaggcggggc ctgcgagcgc ggccaccgag aatcggacgg
5881 gggtagtctc aagctggccg gcctgctctg gtgcctggcc tcgcgccgcc gtgtatcgcc
5941 ccgccctggg cggcaaggct ggcccggtcg gcaccagttg cgtgagcgga aagatggccg
6001 cttcccggcc ctgctgcagg gagctcaaaa tggaggacgc ggcgctcggg agagcgggcg
6061 ggtgagtcac ccacacaaag gaaaagggcc tttccgtcct cagccgtcgc ttcatgtgac
6121 tccacggagt accgggcgcc gtccaggcac ctcgattagt tctcgagctt ttggagtacg
6181 tcgtctttag gttgggggga ggggttttat gcgatggagt ttccccacac tgagtgggtg
6241 gagactgaag ttaggccagc ttggcacttg atgtaattct ccttggaatt tgcccttttt
6301 gagtttggat cttggttcat tctcaagcct cagacagtgg ttcaaagttt ttttcttcca
6361 tttcaggtgt cgtgaagcgg ccgccatgct agtaaattgt ccgctaaatt ctggccgttt
6421 ttggcttttt tgttagacgg atccataact tcgtataatg tatgctatac gaagttatgc
6481 caccatgtct agactggaca agagcaaagt cataaacgga gctctggaat tactcaatgg
6541 tgtcggtatc gaaggcctga cgacaaggaa actcgctcaa aagctgggag ttgagcagcc
6601 taccctgtac tggcacgtga agaacaagcg ggccctgctc gatgccctgc caatcgagat
6661 gctggacagg catcataccc acttctgccc cctggaaggc gagtcatggc aagactttct
6721 gcggaacaac gccaagtcat accgctgtgc tctcctctca catcgcgacg gggctaaagt
6781 gcatctcggc acccgcccaa cagagaaaca gtacgaaacc ctggaaaatc agctcgcgtt
6841 cctgtgtcag caaggcttct ccctggagaa cgcactgtac gctctgtccg ccgtgggcca
6901 ctttacactg ggctgcgtat tggaggaaca ggagcatcaa gtagcaaaag aggaaagaga
6961 gacacctacc accgattcta tgcccccact tctgagacaa gcaattgagc tgttcgaccg
7021 gcagggagcc gaacctgcct tccttttcgg cctggaacta atcatatgtg gcctggagaa
7081 acagctaaag tgcgaaagcg gcgggccgac cgacgccctt gacgattttg acttagacat
7141 gctcccagcc gatgcccttg acgactttga ccttgatatg ctgcctgctg acgctcttga
7201 cgattttgac cttgacatgc tccccgggta actaagtaag gatccgcggc cgcactagag
7261 gaattccgcc cctctccccc ccccccctct ccctcccccc ccccctaacg ttactggccg
7321 aagccgcttg gaataaggcc ggtgtgcgtt tgtctatatg ttattttcca ccatattgcc
7381 gtcttttggc aatgtgaggg cccggaaacc tggccctgtc ttcttgacga gcattcctag
7441 gggtctttcc cctctcgcca aaggaatgca aggtctgttg aatgtcgtga aggaagcagt
7501 tcctctggaa gcttcttgaa gacaaacaac gtctgtagcg accctttgca ggcagcggaa
7561 ccccccacct ggcgacaggt gcctctgcgg ccaaaagcca cgtgtataag atacacctgc
7621 aaaggcggca caaccccagt gccacgttgt gagttggata gttgtggaaa gagtcaaatg
7681 gctctcctca agcgtattca acaaggggct gaaggatgcc cagaaggtac cccattgtat
7741 gggatctgat ctggggcctc ggtgcacatg ctttacatgt gtttagtcga ggttaaaaaa
7801 acgtctaggc cccccgaacc acggggacgt ggttttcctt tgaaaaacac gatgataagc
7861 ttaccggtcc accatggtga gcaagggcga ggagctgttc accggggtgg tgcccatcct
7921 ggtcgagctg gacggcgacg taaacggcca caagttcagc gtgtccggcg agggcgaggg
7981 cgatgccacc tacggcaagc tgaccctgaa gttcatctgc accaccggca agctgcccgt
8041 gccctggccc accctcgtga ccaccctgac ctacggcgtg cagtgcttca gccgctaccc
8101 cgaccacatg aagcagcacg acttcttcaa gtccgccatg cccgaaggct acgtccagga
8161 gcgcaccatc ttcttcaagg acgacggcaa ctacaagacc cgcgccgagg tgaagttcga
8221 gggcgacacc ctggtgaacc gcatcgagct gaagggcatc gacttcaagg aggacggcaa

8281 catcctgggg cacaagctgg agtacaacta caacagccac aacgtctata tcatggccga
8341 caagcagaag aacggcatca aggtgaactt caagatccgc cacaacatcg aggacggcag
8401 cgtgcagctc gccgaccact accagcagaa cacccccatc ggcgacggcc ccgtgctgct
8461 gcccgacaac cactacctga gcacccagtc cgccctgagc aaagacccca acgagaagcg
8521 cgatcacatg gtcctgctgg agttcgtgac cgccgccggg atcactctcg gcatggacga
8581 gctgtacaag taagcatgcc cgacggcgag gatctcgtcg tgacccatgg cgatgcctgc
8641 ttgccgaata tcatggtgga aaatggccgc ttttctggat tcatcgactg tggccggctg
8701 ggtgtggcgg accgctatca ggacatagcg ttggctaccc gtgatattgc tgaagagctt
8761 ggcggcgaat gggctgaccg cttcctcgtg ctttacggta tcgccgctcc cgattcgcag
8821 cgcatcgcct tctatcgcct tcttgacgag ttcttctgat gtacaagtaa agcggccgcg
8881 actctagatc ataatcagcc ataccacatt tgtagaggtt ttacttgctt taaaaaacct
8941 cccacacctc cccctgaacc tgaaacataa aatgaatgca attgttgttg tttagtccct
9001 cccaattcga tatcaagctt atcgcgatag atcctaatca acctctggat tacaaaattt
9061 gtgaaagatt gactggtatt cttaactatg ttgctccttt tacgctatgt ggatacgctg
9121 ctttaatgcc tttgtatcat gctattgctt cccgtatggc tttcattttc tcctccttgt
9181 ataaatcctg gttgctgtct ctttatgagg agttgtggcc cgttgtcagg caacgtggcg
9241 tggtgtgcac tgtgtttgct gacgcaaccc ccactggttg gggcattgcc accacctgtc
9301 agctcctttc cgggactttc gctttccccc tccctattgc cacggcggaa ctcatcgccg
9361 cctgccttgc ccgctgctgg acaggggctc ggctgttggg cactgacaat tccgtggtgt
9421 tgtcggggaa atcatcgtcc tttccttggc tgctcgcctg tgttgccacc tggattctgc
9481 gcgggacgtc cttctgctac gtcccttcgg ccctcaatcc agcggacctt ccttcccgcg
9541 gcctgctgcc ggctctgcgg cctcttccgc gtcttcgcct tcgccctcag acgagtcgga
9601 tctccctttg ggccgcctcc ccgcctgaga tcctttaaga ccaatgactt acaaggcagc
9661 tgtagatctt agccactttt taaaagaaaa ggggggactg gaagggctaa ttcactccca
9721 acgaagacaa gatctgcttt ttgcttgtac tgggtctctc tggttagacc agatctgagc
9781 ctgggagctc tctggctaac tagggaaccc actgcttaag cctcaataaa gcttgccttg
9841 agtgcttcaa gtagtgtgtg cccgtctgtt gtgtgactct ggtaactaga gatccctcag
9901 acccttttag tcagtgtgga aaatctctag cagtagtagt tcatgtcatc ttattattca
9961 gtatttataa cttgcaaaga aatgaatatc agagagtgag aggcccgggt taattaagga
10021 aagggctaga tcattcttga agacgaaagg gcctcgtgat acgcctattt ttataggtta
10081 atgtcatgat aataatggtt tcttagacgt caggtggcac ttttcgggga aatgtgcgcg
10141 gaacccctat ttgtttattt ttctaaatac attcaaatat gtatccgctc atgagacaat
10201 aaccctgata aatgcttcaa taatattgaa aaaggaagag tatgagtatt caacatttcc
10261 gtgtcgccct tattcccttt tttgcggcat tttgccttcc tgtttttgct cacccagaaa
10321 cgctggtgaa agtaaaagat gctgaagatc agttgggtgc acgagtgggt tacatcgaac
10381 tggatctcaa cagcggtaag atccttgaga gttttcgccc cgaagaacgt tttccaatga
10441 tgagcacttt taaagttctg ctatgtggcg cggtattatc ccgtgttgac gccgggcaag
10501 agcaactcgg tcgccgcata cactattctc agaatgactt ggttgagtac tcaccagtca
10561 cagaaaagca tcttacggat ggcatgacag taagagaatt atgcagtgct gccataacca
10621 tgagtgataa cactgcggcc aacttacttc tgacaacgat cggaggaccg aaggagctaa
10681 ccgctttttt gcacaacatg ggggatcatg taactcgcct tgatcgttgg gaaccggagc
10741 tgaatgaagc cataccaaac gacgagcgtg acaccacgat gcctgtagca atggcaacaa
10801 cgttgcgcaa actattaact ggcgaactac ttactctagc ttcccggcaa caattaatag
10861 actggatgga ggcggataaa gttgcaggac cacttctgcg ctcggccctt ccggctggct
10921 ggtttattgc tgataaatct ggagccggtg agcgtgggtc tcgcggtatc attgcagcac
10981 tggggccaga tggtaagccc tcccgtatcg tagttatcta cacgacgggg agtcaggcaa
11041 ctatggatga acgaaataga cagatcgctg agataggtgc ctcactgatt aagcattggt
11101 aactgtcaga ccaagtttac tcatatatac tttagattga tttaaaactt catttttaat
11161 ttaaaaggat ctaggtgaag atcctttttg ataatctcat gaccaaaatc ccttaacgtg
11221 agttttcgtt ccactgagcg tcagaccccg tagaaaagat caaaggatct tcttgagatc
11281 cttttttttct gcgcgtaatc tgctgcttgc aaacaaaaaa accaccgcta ccagcggtgg
11341 tttgtttgcc ggatcaagag ctaccaactc tttttccgaa ggtaactggc ttcagcagag
11401 cgcagatacc aaatactgtt cttctagtgt agccgtagtt aggccaccac ttcaagaact
11461 ctgtagcacc gcctacatac ctcgctctgc taatcctgtt accagtggct gctgccagtg
11521 gcgataagtc gtgtcttacc gggttggact caagacgata gttaccggat aaggcgcagc
11581 ggtcgggctg aacggggggt tcgtgcacac agcccagctt ggagcgaacg acctacaccg
11641 aactgagata cctacagcgt gagctatgag aaagcgccac gcttcccgaa gggagaaagg
11701 cggacaggta tccggtaagc ggcagggtcg gaacaggaga gcgcacgagg gagcttccag
11761 ggggaaacgc ctggtatctt tatagtcctg tcgggtttcg ccacctctga cttgagcgtc
11821 gatttttgtg atgctcgtca ggggggcgga gcctatggaa aaacgccagc aacgcggcct
11881 ttttacggtt cctggccttt tgctggcctt ttgctcacat gttctttcct gcgttatccc
11941 ctgattctgt ggataaccgt attaccgcct ttgagtgagc tgataccgct cgccgcagcc
12001 gaacgaccga gcgcagcgag tcagtgagcg aggaagcgga agagcgccca atacgcaaac
12061 cgcctctccc cgcgcgttgg ccgattcatt aatgcagcaa gctcatggct gactaatttt
12121 ttttatttat gcagaggccg aggccgcctc ggcctctgag ctattccaga agtagtgagg
12181 aggctttttt ggaggcctag gcttttgcaa aaagctcccc gtggcacgac aggtttcccg
12241 actggaaagc gggcagtgag cgcaacgcaa ttaatgtgag ttagctcact cattaggcac

12301 cccaggcttt acactttatg cttccggctc gtatgttgtg tggaattgtg agcggataac
12361 aatttcacac aggaaacagc tatgacatga ttacgaattt cacaaataaa gcattttttt
12421 cactgcattc tagttgtggt ttgtccaaac tcatcaatgt atcttatcat gtctggatca
12481 actggataac tcaagctaac caaaatcatc ccaaacttcc caccccatac cctattacca
12541 ctgccaatta cctgtggttt catttactct aaacctgtga ttcctctgaa ttattttcat
12601 tttaaagaaa ttgtatttgt taaatatgta ctacaaactt agtagt

#### 4. pKG001 (pET-ccdB-GoldenGate)

**Description** pET-type IPTG-inducible *E. coli* expression vector configured as a Golden Gate destination vector: a ccdB counter-selection cassette flanked by inward-facing BsaI sites lies between a vector-encoded start codon and a C-terminal XhoI-LE-His6 tag (T7 promoter/lac operator, lacI, Kan(R)). Parent vector for plasmids 5 and 6.

**Length** 6,060 bp, circular

**Source** This paper

**Note** *BsaI digestion excises the 821-bp region 113..933 carrying ccdB. In plasmids 5 and 6 that cassette is replaced by a 2,173-bp insert encoding His6, a GGGGS linker and PA14_27675; the remaining 5,239 bp of backbone are identical to this vector. The C-terminal His6 at 945..962 is in frame only for an insert that lacks its own stop codon, and is therefore not translated in plasmids 5 and 6.*

##### Features

| Location | Type | Feature |
|---|---|---|
| 21..39 | promoter | T7 promoter |
| 40..64 | protein_bind | lac operator |
| 67..72 | misc_feature | XbaI |
| 79..101 | misc_feature | ribosome binding site |
| 109..111 | CDS | start codon, vector-encoded |
| c(113..118) | misc_feature | BsaI |
| 119..927 | misc_feature | ccdB counter-selection cassette |
| 454..759 | CDS | ccdB |
| 928..933 | misc_feature | BsaI |
| 939..944 | misc_feature | XhoI |
| 939..965 | CDS | XhoI-LE-His6 tag, vector-derived |
| 945..962 | CDS | His6 tag |
| 1029..1076 | terminator | T7 terminator |
| 1113..1568 | rep_origin | f1 ori |
| c(1660..2475) | CDS | aph (kanamycin resistance) |
| 2597..3185 | rep_origin | ori |
| 3371..3510 | misc_feature | bom (basis of mobility) |
| c(3612..3803) | CDS | rop |
| c(4612..5694) | CDS | lacI |
| c(5695..5772) | promoter | lacI promoter |

##### Sequence (6,060 bp)

```
  1 agatctcgat cccgcgaaat taatacgact cactataggg gaattgtgag cggataacaa
 61 ttcccctcta gaaataattt tgtttaactt taagaaggag atatacatat gtgagaccgg
121 atccggctta ctaaaagcca gataacagta tgcgtatttg cgcgctgatt tttgcggtat
181 aagaatatat actgatatgt atacccgaag tatgtcaaaa agaggtatgc tatgaagcag
241 cgtattacag tgacagttga cagcgacagc tatcagttgc tcaaggcata tatgatgtca
301 atatctccgg tctggtaagc acaaccatgc agaatgaagc ccgtcgtctg cgtgccgaac
361 gctggaaagc ggaaaatcag gaagggatgg ctgaggtcgc ccggtttatt gaaatgaacg
421 gctcttttgc tgacgagaac aggggctggt gaaatgcagt ttaaggttta cacctataaa
481 agagagagcc gttatcgtct gtttgtggat gtacagagtg atattattga cacgcccggg
```

541 cgacggatgg tgatccccct ggccagtgca cgtctgctgt cagataaagt cccccgtgaa
601 ctttacccgg tggtgcatat cggggatgaa agctggcgca tgatgaccac cgatatggcc
661 agtgtgccgg tttccgttat cggggaagaa gtggctgatc tcagccaccg cgaaaatgac
721 atcaaaaacg ccattaacct gatgttctgg ggaatataaa tgtcaggctc ccttatacac
781 agccagtctg caggtcgacc atagtgactg gatatgttgt gttttacagt attatgtagt
841 ctgtttttta tgcaaaatct aatttaatat attgatattt atatcatttt acgtttctcg
901 ttcagctttc ttgtacaaag tggttggggt ctcattggct cgagcaccac caccaccacc
961 actgagatcc ggctgctaac aaagcccgaa aggaagctga gttggctgct gccaccgctg
1021 agcaataact agcataaccc cttggggcct ctaaacgggt cttgaggggt tttttgctga
1081 aaggaggaac tatatccgga ttggcgaatg ggacgcgccc tgtagcggcg cattaagcgc
1141 ggcgggtgtg gtggttacgc gcagcgtgac cgctacactt gccagcgccc tagcgcccgc
1201 tcctttcgct ttcttccctt cctttctcgc cacgttcgcc ggctttcccc gtcaagctct
1261 aaatcggggg ctccctttag ggttccgatt tagtgcttta cggcacctcg accccaaaaa
1321 acttgattag ggtgatggtt cacgtagtgg gccatcgccc tgatagacgg tttttcgccc
1381 tttgacgttg gagtccacgt tctttaatag tggactcttg ttccaaactg gaacaacact
1441 caaccctatc tcggtctatt cttttgattt ataagggatt ttgccgattt cggcctattg
1501 gttaaaaaat gagctgattt aacaaaaatt taacgcgaat tttaacaaaa tattaacgct
1561 tacaatttag gtggcacttt tcggggaaat gtgcgcggaa cccctatttg tttatttttc
1621 taaatacatt caaatatgta tccgctcatg aattaattct tagaaaaact catcgagcat
1681 caaatgaaac tgcaatttat tcatatcagg attatcaata ccatattttt gaaaaagccg
1741 tttctgtaat gaaggagaaa actcaccgag gcagttccat aggatggcaa gatcctggta
1801 tcggtctgcg attccgactc gtccaacatc aatacaacct attaatttcc cctcgtcaaa
1861 aataaggtta tcaagtgaga aatcaccatg agtgacgact gaatccggtg agaatggcaa
1921 aagtttatgc atttctttcc agacttgttc aacaggccag ccattacgct cgtcatcaaa
1981 atcactcgca tcaaccaaac cgttattcat tcgtgattgc gcctgagcga gacgaaatac
2041 gcgatcgctg ttaaaaggac aattacaaac aggaatcgaa tgcaaccggc gcaggaacac
2101 tgccagcgca tcaacaatat tttcacctga atcaggatat tcttctaata cctggaatgc
2161 tgttttcccg gggatcgcag tggtgagtaa ccatgcatca tcaggagtac ggataaaatg
2221 cttgatggtc ggaagaggca taaattccgt cagccagttt agtctgacca tctcatctgt
2281 aacatcattg gcaacgctac ctttgccatg tttcagaaac aactctggcg catcgggctt
2341 cccatacaat cgatagattg tcgcacctga ttgcccgaca ttatcgcgag cccatttata
2401 cccatataaa tcagcatcca tgttggaatt taatcgcggc ctagagcaag acgtttcccg
2461 ttgaatatgg ctcataacac cccttgtatt actgtttatg taagcagaca gttttattgt
2521 tcatgaccaa aatcccttaa cgtgagtttt cgttccactg agcgtcagac cccgtagaaa
2581 agatcaaagg atcttcttga gatccttttt ttctgcgcgt aatctgctgc ttgcaaacaa
2641 aaaaaccacc gctaccagcg gtggtttgtt tgccggatca agagctacca actctttttc
2701 cgaaggtaac tggcttcagc agagcgcaga taccaaatac tgtccttcta gtgtagccgt
2761 agttaggcca ccacttcaag aactctgtag caccgcctac atacctcgct ctgctaatcc
2821 tgttaccagt ggctgctgcc agtggcgata agtcgtgtct taccgggttg gactcaagac
2881 gatagttacc ggataaggcg cagcggtcgg gctgaacggg gggttcgtgc acacagccca
2941 gcttggagcg aacgacctac accgaactga gatacctaca gcgtgagcta tgagaaagcg
3001 ccacgcttcc cgaagggaga aaggcggaca ggtatccggt aagcggcagg gtcggaacag
3061 gagagcgcac gagggagctt ccagggggaa acgcctggta tctttatagt cctgtcgggt
3121 ttcgccacct ctgacttgag cgtcgatttt tgtgatgctc gtcagggggg cggagcctat
3181 ggaaaaacgc cagcaacgcg gcctttttac ggttcctggc cttttgctgg ccttttgctc
3241 acatgttctt tcctgcgtta tcccctgatt ctgtggataa ccgtattacc gcctttgagt
3301 gagctgatac cgctcgccgc agccgaacga ccgagcgcag cgagtcagtg agcgaggaag
3361 cggaagagcg cctgatgcgg tattttctcc ttacgcatct gtgcggtatt tcacaccgca
3421 atggtgcact ctcagtacaa tctgctctga tgccgcatag ttaagccagt atacactccg
3481 ctatcgctac gtgactgggt catggctgcg ccccgacacc cgccaacacc cgctgacgcg
3541 ccctgacggg cttgtctgct cccggcatcc gcttacagac aagctgtgac cgtctccggg
3601 agctgcatgt gtcagaggtt ttcaccgtca tcaccgaaac gcgcgaggca gctgcggtaa
3661 agctcatcag cgtggtcgtg aagcgattca cagatgtctg cctgttcatc cgcgtccagc
3721 tcgttgagtt tctccagaag cgttaatgtc tggcttctga taaagcgggc catgttaagg
3781 gcggtttttt cctgtttggt cactgatgcc tccgtgtaag ggggatttct gttcatgggg
3841 gtaatgatac cgatgaaacg agagaggatg ctcacgatac gggttactga tgatgaacat
3901 gcccggttac tggaacgttg tgagggtaaa caactggcgg tatggatgcg gcgggaccag
3961 agaaaaatca ctcagggtca atgccagcgc ttcgttaata cagatgtagg tgttccacag
4021 ggtagccagc agcatcctgc gatgcagatc cggaacataa tggtgcaggg cgctgacttc
4081 cgcgtttcca gactttacga aacacggaaa ccgaagacca ttcatgttgt tgctcaggtc
4141 gcagacgttt tgcagcagca gtcgcttcac gttcgctcgc gtatcggtga ttcattctgc
4201 taaccagtaa ggcaaccccg ccagcctagc cgggtcctca acgacaggag cacgatcatg
4261 cgcacccgtg ggccgccat gccggcgata atggcctgct tctcgccgaa acgtttggtg
4321 gcgggaccag tgacgaaggc ttgagcgagg gcgtgcaaga ttccgaatac cgcaagcgac
4381 aggccgatca tcgtcgcgct ccagcgaaag cggtcctcgc cgaaaatgac ccagagcgct
4441 gccggcacct gtcctacgag ttgcatgata aagaagacag tcataagtgc ggcgacgata
4501 gtcatgcccc gcgcccaccg gaaggagctg actgggttga aggctctcaa gggcatcggt

4561 cgagatcccg gtgcctaatg agtgagctaa cttacattaa ttgcgttgcg ctcactgccc
4621 gctttccagt cgggaaacct gtcgtgccag ctgcattaat gaatcggcca acgcgcgggg
4681 agaggcggtt tgcgtattgg gcgccagggt ggtttttctt ttcaccagtg agacgggcaa
4741 cagctgattg cccttcaccg cctggccctg agagagttgc agcaagcggt ccacgctggt
4801 ttgccccagc aggcgaaaat cctgtttgat ggtggttaac ggcgggatat aacatgagct
4861 gtcttcggta tcgtcgtatc ccactaccga gatatccgca ccaacgcgca gcccggactc
4921 ggtaatggcg cgcattgcgc ccagcgccat ctgatcgttg gcaaccagca tcgcagtggg
4981 aacgatgccc tcattcagca tttgcatggt ttgttgaaaa ccggacatgg cactccagtc
5041 gccttcccgt tccgctatcg gctgaatttg attgcgagtg agatatttat gccagccagc
5101 cagacgcaga cgcgccgaga cagaacttaa tgggcccgct aacagcgcga tttgctggtg
5161 acccaatgcg accagatgct ccacgcccag tcgcgtaccg tcttcatggg agaaaataat
5221 actgttgatg ggtgtctggt cagagacatc aagaaataac gccggaacat tagtgcaggc
5281 agcttccaca gcaatggcat cctggtcatc cagcggatag ttaatgatca gcccactgac
5341 gcgttgcgcg agaagattgt gcaccgccgc tttacaggct tcgacgccgc ttcgttctac
5401 catcgacacc accacgctgg cacccagttg atcggcgcga gatttaatcg ccgcgacaat
5461 ttgcgacggc gcgtgcaggg ccagactgga ggtggcaacg ccaatcagca acgactgttt
5521 gcccgccagt tgttgtgcca cgcggttggg aatgtaattc agctccgcca tcgccgcttc
5581 cactttttcc cgcgttttcg cagaaacgtg gctggcctgg ttcaccacgc gggaaacggt
5641 ctgataagag acaccggcat actctgcgac atcgtataac gttactggtt tcacattcac
5701 caccctgaat tgactctctt ccgggcgcta tcatgccata ccgcgaaagg ttttgcgcca
5761 ttcgatggtg tccgggatct cgacgctctc ccttatgcga ctcctgcatt aggaagcagc
5821 ccagtagtag gttgaggccg ttgagcaccg ccgccgcaag gaatggtgca tgcaaggaga
5881 tggcgcccaa cagtcccccg gccacggggc ctgccaccat acccacgccg aaacaagcgc
5941 tcatgagccc gaagtggcga gcccgatctt ccccatcggt gatgtcggcg atataggcgc
6001 cagcaaccgc acctgtggcg ccggtgatgc cggccacgat gcgtccggcg tagaggatcg

#### 5. LD5562 (pRTP001_IPTG_WT_27675)

**Description** pET-type IPTG-inducible *E. coli* expression vector (T7 promoter/lac operator, Kan(R)) encoding N-terminally His6-tagged, *E. coli* codon-optimized wild-type PA14_27675. Built from plasmid 4 by Golden Gate assembly.

**Length** 7,412 bp, circular

**Source** This paper

**Note** *Expressed product is His6-GGGGS-DeaR, 725 aa (CDS 5231..7408): vector-encoded Met, His6 (5234..5251), GGGGS linker (5252..5266) and the PA14_27675 reading frame (5267..7408, 713 aa). The His6 at 7..24 is the parent vector's C-terminal tag, which lies downstream of the stop codon at 7406..7408 and is not translated.*

##### Features

| Location | Type | Feature |
|---|---|---|
| `1..6` | misc_feature | XhoI |
| `7..24` | misc_feature | His6 tag, vector-derived C-terminal tag (not translated; lies downstream of the stop codon at 7406..7408) |
| `91..138` | terminator | T7 terminator |
| `175..630` | rep_origin | f1 ori |
| `c(722..1537)` | CDS | aph (kanamycin resistance) |
| `1659..2247` | rep_origin | ori |
| `2433..2572` | misc_feature | bom (basis of mobility) |
| `c(2674..2865)` | CDS | rop |
| `c(3674..4756)` | CDS | lacI |
| `c(4757..4834)` | promoter | lacI promoter |
| `5143..5161` | promoter | T7 promoter |
| `5162..5186` | protein_bind | lac operator |
| `5189..5194` | misc_feature | XbaI |
| `5201..5223` | misc_feature | ribosome binding site |
| `5231..7408` | CDS | His6-PA14_27675 (dear), wild type, 725 aa |
| `5234..5251` | misc_feature | His6 tag |

##### Sequence (7,412 bp)

```
   1 ctcgagcacc accaccacca ccactgagat ccggctgcta acaaagcccg aaaggaagct
  61 gagttggctg ctgccaccgc tgagcaataa ctagcataac cccttggggc ctctaaacgg
 121 gtcttgaggg gttttttgct gaaaggagga actatatccg gattggcgaa tgggacgcgc
 181 cctgtagcgg cgcattaagc gcggcgggtg tggtggttac gcgcagcgtg accgctacac
 241 ttgccagcgc cctagcgccc gctcctttcg ctttcttccc ttcctttctc gccacgttcg
 301 ccggctttcc ccgtcaagct ctaaatcggg ggctcccttt agggttccga tttagtgctt
 361 tacggcacct cgaccccaaa aaacttgatt agggtgatgg ttcacgtagt gggccatcgc
 421 cctgatagac ggtttttcgc cctttgacgt tggagtccac gttctttaat agtggactct
 481 tgttccaaac tggaacaaca ctcaacccta tctcggtcta ttcttttgat ttataaggga
 541 ttttgccgat ttcggcctat tggttaaaaa atgagctgat ttaacaaaaa tttaacgcga
 601 attttaacaa aatattaacg cttacaattt aggtggcact tttcggggaa atgtgcgcgg
 661 aacccctatt tgtttatttt tctaaataca ttcaaatatg tatccgctca tgaattaatt
 721 cttagaaaaa ctcatcgagc atcaaatgaa actgcaattt attcatatca ggattatcaa
 781 taccatattt ttgaaaaagc cgtttctgta atgaaggaga aaactcaccg aggcagttcc
 841 ataggatggc aagatcctgg tatcggtctg cgattccgac tcgtccaaca tcaatacaac
 901 ctattaattt cccctcgtca aaaataaggt tatcaagtga gaaatcacca tgagtgacga
 961 ctgaatccgg tgagaatggc aaaagtttat gcatttcttt ccagacttgt tcaacaggcc
```

1021 agccattacg ctcgtcatca aaatcactcg catcaaccaa accgttattc attcgtgatt
1081 gcgcctgagc gagacgaaat acgcgatcgc tgttaaaagg acaattacaa acaggaatcg
1141 aatgcaaccg gcgcaggaac actgccagcg catcaacaat attttcacct gaatcaggat
1201 attcttctaa tacctggaat gctgttttcc cggggatcgc agtggtgagt aaccatgcat
1261 catcaggagt acggataaaa tgcttgatgg tcggaagagg cataaattcc gtcagccagt
1321 ttagtctgac catctcatct gtaacatcat tggcaacgct acctttgcca tgtttcagaa
1381 acaactctgg cgcatcgggc ttcccataca atcgatagat tgtcgcacct gattgcccga
1441 cattatcgcg agcccattta tacccatata aatcagcatc catgttggaa tttaatcgcg
1501 gcctagagca agacgtttcc cgttgaatat ggctcataac accccttgta ttactgttta
1561 tgtaagcaga cagttttatt gttcatgacc aaaatccctt aacgtgagtt ttcgttccac
1621 tgagcgtcag accccgtaga aaagatcaaa ggatcttctt gagatccttt ttttctgcgc
1681 gtaatctgct gcttgcaaac aaaaaaacca ccgctaccag cggtggtttg tttgccggat
1741 caagagctac caactctttt tccgaaggta actggcttca gcagagcgca gataccaaat
1801 actgtccttc tagtgtagcc gtagttaggc caccacttca agaactctgt agcaccgcct
1861 acatacctcg ctctgctaat cctgttacca gtggctgctg ccagtggcga taagtcgtgt
1921 cttaccgggt tggactcaag acgatagtta ccggataagg cgcagcggtc gggctgaacg
1981 gggggttcgt gcacacagcc cagcttggag cgaacgacct acaccgaact gagataccta
2041 cagcgtgagc tatgagaaag cgccacgctt cccgaaggga gaaaggcgga caggtatccg
2101 gtaagcggca gggtcggaac aggagagcgc acgagggagc ttccaggggg aaacgcctgg
2161 tatctttata gtcctgtcgg gtttcgccac ctctgacttg agcgtcgatt tttgtgatgc
2221 tcgtcagggg ggcggagcct atggaaaaac gccagcaacg cggccttttt acggttcctg
2281 gccttttgct ggccttttgc tcacatgttc tttcctgcgt tatcccctga ttctgtggat
2341 aaccgtatta ccgcctttga gtgagctgat accgctcgcc gcagccgaac gaccgagcgc
2401 agcgagtcag tgagcgagga agcggaagag cgcctgatgc ggtattttct ccttacgcat
2461 ctgtgcggta tttcacaccg caatggtgca ctctcagtac aatctgctct gatgccgcat
2521 agttaagcca gtatacactc cgctatcgct acgtgactgg gtcatggctg cgccccgaca
2581 cccgccaaca cccgctgacg cgccctgacg ggcttgtctg ctcccggcat ccgcttacag
2641 acaagctgtg accgtctccg ggagctgcat gtgtcagagg ttttcaccgt catcaccgaa
2701 acgcgcgagg cagctgcggt aaagctcatc agcgtggtcg tgaagcgatt cacagatgtc
2761 tgcctgttca tccgcgtcca gctcgttgag tttctccaga agcgttaatg tctggcttct
2821 gataaagcgg gccatgttaa gggcggtttt ttcctgtttg gtcactgatg cctccgtgta
2881 agggggattt ctgttcatgg gggtaatgat accgatgaaa cgagagagga tgctcacgat
2941 acgggttact gatgatgaac atgcccggtt actggaacgt tgtgagggta aacaactggc
3001 ggtatggatg cggcgggacc agagaaaaat cactcagggt caatgccagc gcttcgttaa
3061 tacagatgta ggtgttccac agggtagcca gcagcatcct gcgatgcaga tccggaacat
3121 aatggtgcag ggcgctgact tccgcgtttc cagactttac gaaacacgga aaccgaagac
3181 cattcatgtt gttgctcagg tcgcagacgt tttgcagcag cagtcgcttc acgttcgctc
3241 gcgtatcggt gattcattct gctaaccagt aaggcaaccc cgccagccta gccgggtcct
3301 caacgacagg agcacgatca tgcgcacccg tggggccgcc atgccggcga taatggcctg
3361 cttctcgccg aaacgtttgg tggcgggacc agtgacgaag gcttgagcga gggcgtgcaa
3421 gattccgaat accgcaagcg acaggccgat catcgtcgcg ctccagcgaa agcggtcctc
3481 gccgaaaatg acccagagcg ctgccggcac ctgtcctacg agttgcatga taaagaagac
3541 agtcataagt gcggcgacga tagtcatgcc ccgcgcccac cggaaggagc tgactgggtt
3601 gaaggctctc aagggcatcg gtcgagatcc cggtgcctaa tgagtgagct aacttacatt
3661 aattgcgttg cgctcactgc ccgctttcca gtcgggaaac ctgtcgtgcc agctgcatta
3721 atgaatcggc caacgcgcgg ggagaggcgg tttgcgtatt gggcgccagg gtggtttttc
3781 ttttcaccag tgagacgggc aacagctgat tgcccttcac cgcctggccc tgagagagtt
3841 gcagcaagcg gtccacgctg gtttgcccca gcaggcgaaa atcctgtttg atggtggtta
3901 acggcgggat ataacatgag ctgtcttcgg tatcgtcgta tcccactacc gagatatccg
3961 caccaacgcg cagcccggac tcggtaatgg cgcgcattgc gcccagcgcc atctgatcgt
4021 tggcaaccag catcgcagtg ggaacgatgc cctcattcag catttgcatg gtttgttgaa
4081 aaccggacat ggcactccag tcgccttccc gttccgctat cggctgaatt tgattgcgag
4141 tgagatattt atgccagcca gccagacgca gacgcgccga gacagaactt aatgggcccg
4201 ctaacagcgc gatttgctgg tgacccaatg cgaccagatg ctccacgccc agtcgcgtac
4261 cgtcttcatg ggagaaaata atactgttga tgggtgtctg gtcagagaca tcaagaaata
4321 acgccggaac attagtgcag gcagcttcca cagcaatggc atcctggtca tccagcggat
4381 agttaatgat cagcccactg acgcgttgcg cgagaagatt gtgcaccgcc gctttacagg
4441 cttcgacgcc gcttcgttct accatcgaca ccaccacgct ggcacccagt tgatcggcgc
4501 gagatttaat cgccgcgaca atttgcgacg gcgcgtgcag ggccagactg gaggtggcaa
4561 cgccaatcag caacgactgt ttgcccgcca gttgttgtgc cacgcggttg ggaatgtaat
4621 tcagctccgc catcgccgct tccacttttt cccgcgtttt cgcagaaacg tggctggcct
4681 ggttcaccac gcgggaaacg gtctgataag agacaccggc atactctgcg acatcgtata
4741 acgttactgg tttcacattc accaccctga attgactctc ttccgggcgc tatcatgcca
4801 taccgcgaaa ggttttgcgc cattcgatgg tgtccgggat ctcgacgctc tcccttatgc
4861 gactcctgca ttaggaagca gcccagtagt aggttgaggc cgttgagcac cgccgccgca
4921 aggaatggtg catgcaagga gatggcgccc aacagtcccc cggccacggg gcctgccacc
4981 atacccacgc cgaaacaagc gctcatgagc ccgaagtggc gagcccgatc ttccccatcg

5041 gtgatgtcgg cgatataggc gccagcaacc gcacctgtgg cgccggtgat gccggccacg
5101 atgcgtccgg cgtagaggat cgagatctcg atcccgcgaa attaatacga ctcactatag
5161 gggaattgtg agcggataac aattcccctc tagaaataat tttgtttaac tttaagaagg
5221 agatatacat atgcatcacc accatcacca cggtggtggc ggatcaatgg catactttaa
5281 gctggaagaa cccgtccgct ttcaccgtta tccgtttgat tttcatagtc actttgcggg
5341 cattctgccg gtagaatcaa actcgcgctg gacacgcgat cgtcgcgtgt ttcgtgtggg
5401 agagcgtcaa gtatctctgg aaaaaggcca ggaactgtcg ctgatcggcc tgcttatgtc
5461 agcgcgcggc gtagcaccgg acgtcgatgg caaagcgttg gaggaagcac gccaggcggc
5521 gcactacgaa ctgtttgaat tggccttaca gcgcatggtg cgtcgcaacc ctttcgccgc
5581 cacggatcgc caaggttact tgcgcggaga gtgtgccgca gaaaatattt acttggcatg
5641 cctgattctg gcccagcgct ttggccgtac ctcaccgccg gcggccattg accaaccggc
5701 gatctatctg ggcacccttg aattactggg tgcgtcagcc gtacgcgact cagaaaccga
5761 ccagtttgta cgttatttta atcgcaaaat ctggagcggt aataagtaca ccccatttga
5821 tgatgcctat tgggcgcgcg gggcgattcg tgatcgccat ccgggcgaat ttgcatgttt
5881 aaccctgggt tttctgctgc atgaaggtat cagccacacc caaacagcaa ccggcgaaga
5941 tgaagtcgcc gtcctggata gcttgtttga acaattcaat gctagcgaaa aaaccgccta
6001 ccgccttctg gctcatacag cccatggcta tgcaagcgag gcagcattcg atgcagagct
6061 gcatcgcatc ttacgccatt tcgagattca gcagggtcag ccaccccagg cacgcctggt
6121 tggtattgat ctgttgggca tggaaacggc gaccggtctt taccgtcaat tttttgactt
6181 tttgctgggc caggcggccg tttttcgtca ttaccttgat ggcaagccgg agacgcgtaa
6241 aatggttctg catatccact gtggcgaagg cactggcgtg agtgacgata accgcagctt
6301 atgcggttat tttttgcgca acgcaaatgc tcttgacgac ttttatgctg ctttaagcgc
6361 atatgcgtgg aaatgctacg gaaatactat tcgtcagggt aaagcgcgct tgcgcgagcg
6421 cgaaaacctg caagaacgcg ataaagcgcc gtcggcgctg gcgggcttat ttgatgaact
6481 ttttttcggg aacagcctga ccgcgtctgg cctgcgctta cgtcgttttg acatcacctc
6541 gggaacgact caggccctgg tggcgtatta cgcccgcacc aacgttgtta acttatgcca
6601 ggcgttggcg tctcgtgacg ccgatggtaa tagctactac cgccgccttt tggaatctga
6661 tttgttttct ctgcgcatcg gccatgcata ttattatcgt aattatcttg cctcaaagtt
6721 tccggaatta tgtttcgata ctaatttggg ctctaacttt atcactgggg cgagcggttt
6781 attcgattct cttcaggagt accgtttaaa ccgcggcctg cgccatctgg atggatatgt
6841 cggcacagat caacttaaag agttatcgct ggcaattgcc taccagggtg agcaacgtct
6901 ggaccctcag cagatgcagt atgttcatgc tctggcggaa tcccagagcg gctttgacga
6961 actgggcgaa cacctgccag gaacgccggg ctgggcaaaa ccggctttag aacagttttt
7021 tgtcagccag tgtgccctgt accgttccga ggaggatcgt tactttcagt ttgaggccta
7081 tcgtcgtttg tttgcacaag ttctgaactg gcgttcctat cttctgggtg ccgatggtca
7141 aggtgtggaa cacagcaatg tgcaggacga agcaattcgc atggcactgt tgctgaatta
7201 cgcggcggcg gatcgccatg gccgtgttcc ggccgcatcg ctggaaaatg cgcagcgctt
7261 gcttgttcag ctgggaagtg catattggga ggaaaccatt ggcgcagtcg atctggcggg
7321 cgccccgcat cgcgatcgtg aactgcagcg ctttgaaggc tttgccgcgc cggcgagcgt
7381 tgtgcgcatt agcacgcgct cttcttaatt gg

#### 6. LD5563 (pRTP010_27675_HA_IPTG)

**Description** As plasmid 5, encoding the His6-tagged H22A allele of PA14_27675 (*E. coli* codon-optimized). Built from plasmid 4 by Golden Gate assembly.

**Length** 7,412 bp, circular

**Source** This paper

**Note** *H22A codon at 5330..5332 (GCC). Expressed product is His6-GGGGS-DeaR(H22A), 725 aa. As in plasmid 5, the His6 at 7..24 is the parent vector's C-terminal tag and is not translated.*

##### Features

| Location | Type | Feature |
|---|---|---|
| 1..6 | misc_feature | XhoI |
| 7..24 | misc_feature | His6 tag, vector-derived C-terminal tag (not translated; lies downstream of the stop codon at 7406..7408) |
| 91..138 | terminator | T7 terminator |
| 175..630 | rep_origin | f1 ori |
| c(722..1537) | CDS | aph (kanamycin resistance) |
| 1659..2247 | rep_origin | ori |
| 2433..2572 | misc_feature | bom (basis of mobility) |
| c(2674..2865) | CDS | rop |
| c(3674..4756) | CDS | lacI |
| c(4757..4834) | promoter | lacI promoter |
| 5143..5161 | promoter | T7 promoter |
| 5162..5186 | protein_bind | lac operator |
| 5189..5194 | misc_feature | XbaI |
| 5201..5223 | misc_feature | ribosome binding site |
| 5231..7408 | CDS | His6-PA14_27675 (dear), H22A, 725 aa |
| 5234..5251 | misc_feature | His6 tag |
| 5330..5332 | misc_feature | H22A codon (GCC) |

##### Sequence (7,412 bp)

```
   1 ctcgagcacc accaccacca ccactgagat ccggctgcta acaaagcccg aaaggaagct
  61 gagttggctg ctgccaccgc tgagcaataa ctagcataac cccttggggc ctctaaacgg
 121 gtcttgaggg gttttttgct gaaaggagga actatatccg gattggcgaa tgggacgcgc
 181 cctgtagcgg cgcattaagc gcggcgggtg tggtggttac gcgcagcgtg accgctacac
 241 ttgccagcgc cctagcgccc gctcctttcg ctttcttccc ttcctttctc gccacgttcg
 301 ccggctttcc ccgtcaagct ctaaatcggg ggctcccttt agggttccga tttagtgctt
 361 tacggcacct cgaccccaaa aaacttgatt agggtgatgg ttcacgtagt gggccatcgc
 421 cctgatagac ggtttttcgc cctttgacgt tggagtccac gttctttaat agtggactct
 481 tgttccaaac tggaacaaca ctcaacccta tctcggtcta ttcttttgat ttataaggga
 541 ttttgccgat ttcggcctat tggttaaaaa atgagctgat ttaacaaaaa tttaacgcga
 601 attttaacaa aatattaacg cttacaattt aggtggcact tttcggggaa atgtgcgcgg
 661 aacccctatt tgtttatttt tctaaataca ttcaaatatg tatccgctca tgaattaatt
 721 cttagaaaaa ctcatcgagc atcaaatgaa actgcaattt attcatatca ggattatcaa
 781 taccatattt ttgaaaaagc cgtttctgta atgaaggaga aaactcaccg aggcagttcc
 841 ataggatggc aagatcctgg tatcggtctg cgattccgac tcgtccaaca tcaatacaac
 901 ctattaattt cccctcgtca aaaataaggt tatcaagtga gaaatcacca tgagtgacga
 961 ctgaatccgg tgagaatggc aaaagtttat gcatttcttt ccagacttgt tcaacaggcc
1021 agccattacg ctcgtcatca aaatcactcg catcaaccaa accgttattc attcgtgatt
1081 gcgcctgagc gagacgaaat acgcgatcgc tgttaaaagg acaattacaa acaggaatcg
```

1141 aatgcaaccg gcgcaggaac actgccagcg catcaacaat attttcacct gaatcaggat
1201 attcttctaa tacctggaat gctgttttcc cggggatcgc agtggtgagt aaccatgcat
1261 catcaggagt acggataaaa tgcttgatgg tcggaagagg cataaattcc gtcagccagt
1321 ttagtctgac catctcatct gtaacatcat tggcaacgct acctttgcca tgtttcagaa
1381 acaactctgg cgcatcgggc ttcccataca atcgatagat tgtcgcacct gattgcccga
1441 cattatcgcg agcccattta tacccatata aatcagcatc catgttggaa tttaatcgcg
1501 gcctagagca agacgtttcc cgttgaatat ggctcataac accccttgta ttactgttta
1561 tgtaagcaga cagttttatt gttcatgacc aaaatccctt aacgtgagtt ttcgttccac
1621 tgagcgtcag accccgtaga aaagatcaaa ggatcttctt gagatccttt ttttctgcgc
1681 gtaatctgct gcttgcaaac aaaaaaacca ccgctaccag cggtggtttg tttgccggat
1741 caagagctac caactctttt tccgaaggta actggcttca gcagagcgca gataccaaat
1801 actgtccttc tagtgtagcc gtagttaggc caccacttca agaactctgt agcaccgcct
1861 acatacctcg ctctgctaat cctgttacca gtggctgctg ccagtggcga taagtcgtgt
1921 cttaccgggt tggactcaag acgatagtta ccggataagg cgcagcggtc gggctgaacg
1981 gggggttcgt gcacacagcc cagcttggag cgaacgacct acaccgaact gagataccta
2041 cagcgtgagc tatgagaaag cgccacgctt cccgaaggga gaaaggcgga caggtatccg
2101 gtaagcggca gggtcggaac aggagagcgc acgagggagc ttccaggggg aaacgcctgg
2161 tatctttata gtcctgtcgg gtttcgccac ctctgacttg agcgtcgatt tttgtgatgc
2221 tcgtcagggg ggcggagcct atggaaaaac gccagcaacg cggccttttt acggttcctg
2281 gccttttgct ggccttttgc tcacatgttc tttcctgcgt tatcccctga ttctgtggat
2341 aaccgtatta ccgcctttga gtgagctgat accgctcgcc gcagccgaac gaccgagcgc
2401 agcgagtcag tgagcgagga agcggaagag cgcctgatgc ggtattttct ccttacgcat
2461 ctgtgcggta tttcacaccg caatggtgca ctctcagtac aatctgctct gatgccgcat
2521 agttaagcca gtatacactc cgctatcgct acgtgactgg gtcatggctg cgccccgaca
2581 cccgccaaca cccgctgacg cgccctgacg ggcttgtctg ctcccggcat ccgcttacag
2641 acaagctgtg accgtctccg ggagctgcat gtgtcagagg ttttcaccgt catcaccgaa
2701 acgcgcgagg cagctgcggt aaagctcatc agcgtggtcg tgaagcgatt cacagatgtc
2761 tgcctgttca tccgcgtcca gctcgttgag tttctccaga agcgttaatg tctggcttct
2821 gataaagcgg gccatgttaa gggcggtttt ttcctgtttg gtcactgatg cctccgtgta
2881 agggggattt ctgttcatgg gggtaatgat accgatgaaa cgagagagga tgctcacgat
2941 acgggttact gatgatgaac atgcccggtt actggaacgt tgtgagggta aacaactggc
3001 ggtatggatg cggcgggacc agagaaaaat cactcagggt caatgccagc gcttcgttaa
3061 tacagatgta ggtgttccac agggtagcca gcagcatcct gcgatgcaga tccggaacat
3121 aatggtgcag ggcgctgact tccgcgtttc cagactttac gaaacacgga aaccgaagac
3181 cattcatgtt gttgctcagg tcgcagacgt tttgcagcag cagtcgcttc acgttcgctc
3241 gcgtatcggt gattcattct gctaaccagt aaggcaaccc cgccagccta gccgggtcct
3301 caacgacagg agcacgatca tgcgcacccg tggggccgcc atgccggcga taatggcctg
3361 cttctcgccg aaacgtttgg tggcgggacc agtgacgaag gcttgagcga gggcgtgcaa
3421 gattccgaat accgcaagcg acaggccgat catcgtcgcg ctccagcgaa agcggtcctc
3481 gccgaaaatg acccagagcg ctgccggcac ctgtcctacg agttgcatga taaagaagac
3541 agtcataagt gcggcgacga tagtcatgcc ccgcgcccac cggaaggagc tgactgggtt
3601 gaaggctctc aagggcatcg gtcgagatcc cggtgcctaa tgagtgagct aacttacatt
3661 aattgcgttg cgctcactgc ccgctttcca gtcgggaaac ctgtcgtgcc agctgcatta
3721 atgaatcggc caacgcgcgg ggagaggcgg tttgcgtatt gggcgccagg gtggtttttc
3781 ttttcaccag tgagacgggc aacagctgat tgcccttcac cgcctggccc tgagagagtt
3841 gcagcaagcg gtccacgctg gtttgcccca gcaggcgaaa atcctgtttg atggtggtta
3901 acggcgggat ataacatgag ctgtcttcgg tatcgtcgta tcccactacc gagatatccg
3961 caccaacgcg cagcccggac tcggtaatgg cgcgcattgc gcccagcgcc atctgatcgt
4021 tggcaaccag catcgcagtg ggaacgatgc cctcattcag catttgcatg gtttgttgaa
4081 aaccggacat ggcactccag tcgccttccc gttccgctat cggctgaatt tgattgcgag
4141 tgagatattt atgccagcca gccagacgca gacgcgccga gacagaactt aatgggcccg
4201 ctaacagcgc gatttgctgg tgacccaatg cgaccagatg ctccacgccc agtcgcgtac
4261 cgtcttcatg ggagaaaata atactgttga tgggtgtctg gtcagagaca tcaagaaata
4321 acgccggaac attagtgcag gcagcttcca cagcaatggc atcctggtca tccagcggat
4381 agttaatgat cagcccactg acgcgttgcg cgagaagatt gtgcaccgcc gctttacagg
4441 cttcgacgcc gcttcgttct accatcgaca ccaccacgct ggcacccagt tgatcggcgc
4501 gagatttaat cgccgcgaca atttgcgacg gcgcgtgcag ggccagactg gaggtggcaa
4561 cgccaatcag caacgactgt ttgcccgcca gttgttgtgc cacgcggttg ggaatgtaat
4621 tcagctccgc catcgccgct tccacttttt cccgcgtttt cgcagaaacg tggctggcct
4681 ggttcaccac gcgggaaacg gtctgataag agacaccggc atactctgcg acatcgtata
4741 acgttactgg tttcacattc accaccctga attgactctc ttccgggcgc tatcatgcca
4801 taccgcgaaa ggttttgcgc cattcgatgg tgtccgggat ctcgacgctc tcccttatgc
4861 gactcctgca ttaggaagca gcccagtagt aggttgaggc cgttgagcac cgccgccgca
4921 aggaatggtg catgcaagga gatggcgccc aacagtcccc cggccacggg gcctgccacc
4981 atacccacgc cgaaacaagc gctcatgagc ccgaagtggc gagcccgatc ttccccatcg
5041 gtgatgtcgg cgatataggc gccagcaacc gcacctgtgg cgccggtgat gccggccacg
5101 atgcgtccgg cgtagaggat cgagatctcg atcccgcgaa attaatacga ctcactatag

5161 gggaattgtg agcggataac aattcccctc tagaaataat tttgtttaac tttaagaagg
5221 agatatacat atgcatcacc accatcacca cggtggtggc ggatcaatgg cctactttaa
5281 actggaggaa ccggtgcgct tccaccgcta tccgttcgat ttccatagcg cgtttgcggg
5341 tattctgccg gtcgagagca acagccgctg gacacgtgac cgccgcgtgt ttcgcgtagg
5401 cgaacgccag gtttcccttg aaaaaggtca ggagttatca ttgatcggct tgctgatgag
5461 cgcccgcggt gtcgcgccgg atgtggatgg taaagcgtta gaggaagcgc gccaagccgc
5521 ccattatgaa ctgtttgaat tagccctgca acgtatggtt cgtcgtaatc cattcgcggc
5581 gacggatcgt cagggctact tacgcggtga atgcgcagcg gaaaatatct acctggcgtg
5641 tttaatcttg gcacagcgct ttggtcgtac cagcccgccg gcagcaatcg atcaaccggc
5701 catttacctg ggcactctgg agctgcttgg cgcgagcgca gtgcgcgata gcgaaaccga
5761 tcagtttgtg cgttatttca atcgtaaaat ctggtccggc aataaataca ctccatttga
5821 tgacgcgtat tgggcgcgtg gcgcgattcg cgaccgtcac ccggcgaat ttgcttgctt
5881 aactctgggc ttcctgttac acgaagggat tagtcatacc cagaccgcca caggcgaaga
5941 tgaggttgcg gtgctggatt cattattcga acagtttaat gcctctgaga agacagccta
6001 ccgtcttctg gcccatactg ctcacggcta cgcgtcagaa gcggcgttcg atgcagagtt
6061 gcaccgcatt ttgcgtcact ttgaaattca gcaggggcaa ccaccccagg cgcgtctggt
6121 gggtattgat ctgctgggca tggaaacagc aaccgggttg tatcgccagt tcttcgattt
6181 tttgttgggc caggcagcgg tgtttcgtca ctacttggat ggcaaaccgg aaacgcgtaa
6241 gatggtgtta catattcact gcggtgaggg tacgggcgtg agcgatgaca atcgctctct
6301 gtgcggctat tttctgcgta acgccaatgc gctggatgat ttttatgcgg cgctgagcgc
6361 gtacgcgtgg aagtgctacg gcaacactat ccgtcagggt aaagcacgcc tgcgtgagcg
6421 cgagaatctg caggaacgcg acaaggcccc tagtgccctg gcaggtctgt ttgatgaact
6481 tttttttgga aactcgctga ccgcatcagg ccttcgttta cgccgttttg acatcacatc
6541 cggtacgacc caggcgcttg tagcttatta cgcgcgcaca aatgtcgtga acctgtgcca
6601 ggctctggcg tctcgcgatg cggatggaaa cagctactat cgtcgtctgc tggaatctga
6661 tctgttcagc ttacgcattg gtcacgcata ttactaccgc aactatttag cgagcaaatt
6721 cccggagctg tgttttgata ccaatttagg cagcaatttt atcacgggcg cgagcggctt
6781 atttgattct ttgcaggaat atcgccttaa ccgtggactt cgccatctgg atggctatgt
6841 cggcacggat caactgaaag aactgtcgct ggcaattgcg tatcagggcg aacaacgcct
6901 ggatccgcag cagatgcagt acgtgcatgc ccttgcggaa tcgcagagcg ggttcgacga
6961 acttggcgaa cacttgcctg gcacaccggg ttgggcgaaa ccagccttag aacagttttt
7021 cgtaagccag tgtgccttgt accgcagcga agaagaccgc tatttccagt ttgaggccta
7081 ccgtcgcctg tttgctcagg ttctgaactg gcgctcatac ctgctgggcg cggatggcca
7141 gggcgttgag catagcaacg ttcaagacga agcgatccgt atggccctgc ttctgaacta
7201 tgcagcagcc gaccgccacg gccgcgtgcc tgccgcgagc ttagagaacg cacagcgtct
7261 gttggttcag ttgggaagcg cgtattggga ggaaaccatc ggcgccgtgg atctggcggg
7321 agcaccccat cgcgatcgcg aacttcagcg ttttgaaggc ttcgcggcgc ctgcctccgt
7381 ggtgcgcatc tcgacccgta gttcgtaatt gg

#### 7. pMQ30-Δ27675

**Description** Allelic-exchange construct for the in-frame deletion of PA14_27675 (dear) in *P. aeruginosa* PA14. pMQ30 allelic-exchange (suicide) vector: Gm(R) aacC1, sacB counter-selection, oriT, URA3 + CEN6/ARSH4 for yeast gap-repair assembly (Shanks et al. 2006)

**Length** 9,178 bp, circular

**Source** This paper

**Note** *Insert carries the PA14_27650-PA14_27680 flanking region with PA14_27675 deleted; 39 bp of the native gene are retained at 4338..4376.*

##### Features

| Location | Type | Feature |
|---|---|---|
| c(209..1630) | gene | sacB |
| 2104..2928 | misc_feature | oriT (conjugal transfer origin, from pEX18-Gm) |
| 3341..3816 | misc_feature | PA14_27650 |
| 3870..4109 | misc_feature | PA14_27660 |
| 4338..4376 | misc_feature | PA14_27675 deletion junction (39 bp retained) |
| 4427..5059 | misc_feature | PA14_27680 |
| 5724..6324 | rep_origin | ori (ColE1) |
| c(6588..7121) | gene | aacC1 (gentamicin resistance) |
| c(7557..8360) | gene | URA3 |
| 8599..9112 | misc_feature | CEN6/ARSH4; *S. cerevisiae* replicon from pYC2-CT |

##### Sequence (9,178 bp)

```
   1 tcgcgcgttt cggtgatgac ggtgaaaacc tctgacacat gcagctcccg gagacggtca
  61 cagcttgtct gtaagcggat gccgggagca gacaagcccg tcagggcgcg tcagcgggtg
 121 ttggcgggtg tcggggctgg cttaactatg cggcatcaga gcagattgta ctgagagtgc
 181 accataatcg gcattttctt ttgcgttttt atttgttaac tgttaattgt ccttgttcaa
 241 ggatgctgtc tttgacaaca gatgttttct tgcctttgat gttcagcagg aagctaggcg
 301 caaacgttga ttgtttgtct gcgtagaatc ctctgtttgt catatagctt gtaatcacga
 361 cattgtttcc tttcgcttga ggtacagcga agtgtgagta agtaaaggtt acatcgttag
 421 gatcaagatc cattttaac acaaggccag ttttgttcag cggcttgtat gggccagtta
 481 aagaattaga aacataacca agcatgtaaa tatcgttaga cgtaatgccg tcaatcgtca
 541 tttttgatcc gcgggagtca gtgaacagat accatttgcc gttcatttta aagacgttcg
 601 cgcgttcaat ttcatctgtt actgtgttag atgcaatcag cggtttcatc actttttca
 661 gtgtgtaatc atcgtttagc tcaatcatac cgagagcgcc gtttgctaac tcagccgtgc
 721 gtttttatc gctttgcaga agtttttgac tttcttgacg gaagaatgat gtgcttttgc
 781 catagtatgc tttgttaaat aaagattctt cgccttggta gccatcttca gttccagtgt
 841 ttgcttcaaa tactaagtat ttgtggcctt tatcttctac gtagtgagga tctctcagcg
 901 tatggttgtc gcctgagctg tagttgcctt catcgatgaa ctgctgtaca ttttgatacg
 961 tttttccgtc accgtcaaag attgatttat aatcctctac accgttgatg ttcaaagagc
1021 tgtctgatgc tgatacgtta acttgtgcag ttgtcagtgt ttgtttgccg taatgtttac
1081 cggagaaatc agtgtagaat aaacggattt ttccgtcaga tgtaaatgtg gctgaacctg
1141 accattcttg tgtttggtct tttaggatag aatcatttgc atcgaatttg tcgctgtctt
1201 taaagacgcg gccagcgttt ttccagctgt caatagaagt ttcgccgact ttttgataga
1261 acatgtaaat cgatgtgtca tccgcatttt taggatctcc ggctaatgca aagacgatgt
1321 ggtagccgtg atagtttgcg acagtgccgt cagcgttttg taatggccag ctgtcccaaa
1381 cgtccaggcc ttttgcagaa gagatatttt taattgtgga cgaatcgaac tcaggaactt
1441 gatatttttc atttttttgc tgttcaggga tttgcagcat atcatggcgt gtaatatggg
1501 aaatgccgta tgtttcctta tatggctttt ggttcgtttc tttcgcaaac gcttgagttg
1561 cgcctcctgc cagcagtgcg gtagtaaagg ttaatactgt tgcttgtttt gcaaactttt
1621 tgatgttcat cgttcatgtc tcctttttta tgtactgtgt tagcggtctg cttcttccag
1681 ccctcctgtt tgaagatggc aagttagtta cgcacaataa aaaaagacct aaaatatgta
1741 aggggtgacg ccaaagtata cactttgccc tttacacatt ttaggtcttg cctgctttat
```

1801 cagtaacaaa cccgcgcgat ttacttttcg acctcattct attagactct cgtttggatt
1861 gcaactggtc tattttcctc ttttgtttga tagaaaatca taaaaggatt tgcagactac
1921 gggcctaaag aactaaaaaa tctatctgtt tcttttcatt ctctgtattt tttatagttt
1981 ctgttgcatg ggcataaagt tgccttttta atcacaattc agaaaatatc ataatatctc
2041 atttcactaa ataatagtga acggcaggta tatgtgatgg gttaaaaagg atcgatcctc
2101 tagctagagt cgatcttcgc cagcagggcg aggatcgtgg catcaccgaa ccgcgccgtg
2161 cgcgggtcgt cggtgagcca gagtttcagc aggccgccca ggcggcccag gtcgccattg
2221 atgcgggcca gctcgcggac gtgctcatag tccacgacgc ccgtgatttt gtagccctgg
2281 ccgacggcca gcaggtaggc cgacaggctc atgccggccg ccgccgcctt ttcctcaatc
2341 gctcttcgtt cgtctggaag gcagtacacc ttgataggtg ggctgccctt cctggttggc
2401 ttggtttcat cagccatccg cttgccctca tctgttacgc cggcggtagc cggccagcct
2461 cgcagagcag gattcccgtt gagcaccgcc aggtgcgaat aagggacagt gaagaaggaa
2521 cacccgctcg cgggtgggcc tacttcacct atcctgcccg gctgacgccg ttggatacac
2581 caaggaaagt ctacacgaac cctttggcaa aatcctgtat atcgtgcgaa aaaggatgga
2641 tataccgaaa aaatcgctat aatgaccccg aagcagggtt atgcagcgga aaagcgctgc
2701 ttccctgctg ttttgtggaa tatctaccga ctggaaacag gcaaatgcag gaaattactg
2761 aactgagggg acaggcgaga gacgatgcca aagagctaca ccgacgagct ggccgagtgg
2821 gttgaatccc gcgcggccaa gaagcgccgg cgtgatgagg ctgcggttgc gttcctggcg
2881 gtgagggcgg atgtcgatat gcgtaaggag aaaataccgc atcaggcgca tatttgaatg
2941 tatttagaaa aataaacaaa aagagtttgt agaaacgcaa aaaggccatc cgtcaggatg
3001 gccttctgct taatttgatg cctggcagtt tatggcgggc gtcctgcccg ccaccctccg
3061 ggccgttgct tcgcaacgtt caaatccgct cccggcggat ttgtcctact caggagagcg
3121 ttcaccgaca aacaacagat aaaacgaaag gcccagtctt tcgactgagc ctttcgtttt
3181 atttgatgcc tggcagttcc ctactctcgc atggggagac cccacactac catcggcgct
3241 acggcgtttc acttctgagt tcggcatggg gtcaggtggg accaccgcgc tactgccgcc
3301 aggcaaattc tgttttatca gaccgcttct gcgttctgat gcgatagcgt tctccagttg
3361 ctgcttgagc gagccgaagt tctgctcgaa gcacaggcgt acctcgctct tcactaggtt
3421 ggccaggcgg tcggcaatct gcgacatcac caactggttg gcgcttgcct ggccgctctg
3481 caggtggccg ggacgcagtt cggccaggct gccggcggca ttgaccgcct cgagaatcgc
3541 ctgctcgccg ctgtcctggg cgcggagggc ctgggtcgag gcctgctgga cgttctgcag
3601 gatctggcgg atggagcctt cggaacgctg ccggccgttc tcgatcacct gttcggcctg
3661 ctgtcgcgcg tgcacggccg gatcgctgct ggcgtcgttg ccgaggccga gctggaactg
3721 tctcaactgc tcgtgttctt cgctggtgag gggcgtttc agccagcgtt gcgccagctg
3781 gaccagattc cttcccaatt cttccgagat cgacatgcgc ggactcctct gcgatggacg
3841 cgcggtgcgc ggggtgtcgt ggcgaggctt cagtactgct gcgtgccgtc cgccgagggg
3901 gcctgcgcgc tgctgccggt atcgccgggg taggcggcgg cagcggcctg cagggtgctg
3961 taggcggtgc tcaactgcac cgccaactgt ccgaggctgg ccgggtcgtc cacgctgtct
4021 ccctgggaca gggcctggag ccagccttgc aggcattcgg tgaggagtgc cttgacctgg
4081 gtgacgaggg tttcgagttc ttccttcatg acgcttctcc tggggtgggt gctcatccga
4141 gcagtttcag cacggccaag agcttggccg catcgctgtc cgatatttcc agcaggcgct
4201 ttccgctcgc cggcctgctg gggggcggtt ggtcccgtgc ctcctcgctg gcgtccggcg
4261 aacgctgcgc cgggcggtcc gcggcggcga gcgcgcggcg tctgatgtag cgtgcgtagg
4321 cgctcatcgg tcgctcctca actcctgacc ggttcttcca gcttgaaata ggccatgacg
4381 gctctcctgt gccagggccg gacgccgccg gccccgccgg tggtggctac gattgggtgt
4441 ccttgccgcc gcgactggcg agggtctcgc tgtacttggc gatgctctgg tcgatcttgt
4501 cgatggcgaa gctggccgat tccgcctcga tcgccccggc cgccgcggct accgcggcgg
4561 cggcgcccat cagcaggtcg gtggcgaggg cgccgagcgc gtcctcggca gcctgggcca
4621 gcttgttttc ggcgatgttc tgggtcatct gccggagcag gatcgcctgg ctggccaggg
4681 cgatcttgct gactccgtcg aagtagctgg cggcggattg cgccaccagc ccggaggcct
4741 ggctgatcat catgttcggc gcgatggcga tctgcgacgg cgcatagccg gcggtttcgc
4801 tgttgctcat ctgcaccgcc tgcacgactt gactgttcag ggcgctggtg tccggcggcg
4861 gcgcctcggt ctgcttgagc aggcgctcgc tcatgccggc cagcggcgaa gggcgcgacg
4921 gcagcttcac gtcgccgttg gcggacgccc tgccgccgct ggctggcgtc tgccggccgg
4981 cctcgccggg ggcggcggct tgctcgctgt cgccggatgg cgcgggcgcg ggcgcggatt
5041 tcttgccgaa gccgaacatg tctgggcact cctgggatca aggggcgggt cagatacctt
5101 tcttgtcttc tatcgccatg atcatcgaca ccgccttgga aaccaccgcg gtggagatct
5161 ggttcagggc ctgctggctg tagactgcgt tctgcaggga caggccggcg gaatggctgg
5221 cgatctggta gagcgcggcg atggcctggg ccggggcttc ggcgaccacc ttgacgttgg
5281 tctgggtgac ggcatcggtg atctgggcat tcacagccgt ggggtcagcc atgggactct
5341 ccttgaatgg gcagtttcct gtgtgaaatt gttatccgct cacaattcca cacaacatac
5401 gagccggaag cataaagtgt aaagcctggg gtgcctaatg agtgagctaa ctcacattaa
5461 ttgcgttgcg ctcactgccc gctttccagt cgggaaacct gtcgtgccag ctgcattaat
5521 gaatcggcca acgcgcgggg agaggcggtt tgcgtattgg gcgctcttcc gcttcctcgc
5581 tcactgactc gctgcgctcg gtcgttcggc tgcggcgagc ggtatcagct cactcaaagg
5641 cggtaatacg gttatccaca gaatcagggg ataacgcagg aaagaacatg tgagcaaaag
5701 gccagcaaaa ggccaggaac cgtaaaaagg ccgcgttgct ggcgtttttc cataggctcc
5761 gcccccctga cgagcatcac aaaaatcgac gctcaagtca gaggtggcga aacccgacag

5821 gactataaag ataccaggcg tttccccctg gaagctccct cgtgcgctct cctgttccga
5881 ccctgccgct taccggatac ctgtccgcct ttctcccttc gggaagcgtg gcgctttctc
5941 atagctcacg ctgtaggtat ctcagttcgg tgtaggtcgt tcgctccaag ctgggctgtg
6001 tgcacgaacc ccccgttcag cccgaccgct gcgccttatc cggtaactat cgtcttgagt
6061 ccaacccggt aagacacgac ttatcgccac tggcagcagc cactggtaac aggattagca
6121 gagcgaggta tgtaggcggt gctacagagt tcttgaagtg gtggcctaac tacggctaca
6181 ctagaaggac agtatttggt atctgcgctc tgctgaagcc agttaccttc ggaaaaagag
6241 ttggtagctc ttgatccggc aaacaaacca ccgctggtag cggtggtttt tttgtttgca
6301 agcagcagat tacgcgcaga aaaaaaggat ctcaagaaga tcctttgatc ttttctacgg
6361 ggtctgacgc tcagtggaac gaaaactcac gttaagggat tttggtcatg agattatcaa
6421 aaaggatctt cacctagatc cttttaaatt aaaaatgaag ttttaaatca atctaaagta
6481 tatatgagta aacttggtct gacaatcgat gcgaattggc cgcggcgttg tgacaattta
6541 ccgaacaact ccgcggccgg gaagccgatc tcggcttgaa cgaattgtta ggtggcggta
6601 cttgggtcga tatcaaagtg catcacttct tcccgtatgc ccaactttgt atagagagcc
6661 actgcgggat cgtcaccgta atctgcttgc acgtagatca cataagcacc aagcgcgttg
6721 gcctcatgct tgaggagatt gatgagcgcg gtggcaatgc cctgcctccg gtgctcgccg
6781 gagactgcga gatcatagat atagatctca ctacgcggct gctcaaacct gggcagaacg
6841 taagccgcga gagcgccaac aaccgcttct tggtcgaagg cagcaagcgc gatgaatgtc
6901 ttactacgga gcaagttccc gaggtaatcg gagtccggct gatgttggga gtaggtggct
6961 acgtctccga actcacgacc gaaaagatca agagcagccc gcatggattt gacttggtca
7021 gggccgagcc tacatgtgcg aatgatgccc atacttgagc cacctaactt tgttttaggg
7081 cgactgccct gctgcgtaac atcgttgctg ctgcgtaaca tcgttgctgc tccataacat
7141 caaacatcga cccacggcgt aacgcgcttg ctgcttggat gcccgaggca tagactgtac
7201 aaaaaaacag tcataacaag ccatgaaaac cgccactgcg ccgttaccac cgctgcgttc
7261 ggtcaaggtt ctggaccagt tgcgtgagcg catacgctac ttgcattaca gtttacgaac
7321 cgaacaggct tatgtcaatt cggttgaata ctcatactct tcctttttca atattattga
7381 agcatttatc agggttattg tctcatgagc ggatacatat ttgaatgtat ttagaaaaat
7441 aaacaaatag gggttccgcg cacatttccc cgaaaagtgg gtaataactg atataattaa
7501 attgaagctc taatttgtga gtttagtata catgcattta cttataatac agttttttag
7561 ttttgctggc cgcatcttct caaatatgct tcccagcctg cttttctgta acgttcaccc
7621 tctaccttag catcccttcc ctttgcaaat agtcctcttc caacaataat aatgtcagat
7681 cctgtagaga ccacatcatc cacggttcta tactgttgac ccaatgcgtc tcccttgtca
7741 tctaaaccca caccgggtgt cataatcaac caatcgtaac cttcatctct tccacccatg
7801 tctctttgag caataaagcc gataacaaaa tctttgtcgc tcttcgcaat gtcaacagta
7861 cccttagtat attctccagt agatagggag cccttgcatg acaattctgc taacatcaaa
7921 aggcctctag gttcctttgt tacttcttct gccgcctgct tcaaaccgct aacaatacct
7981 gggcccacca caccgtgtgc attcgtaatg tctgcccatt ctgctattct gtatacaccc
8041 gcagagtact gcaatttgac tgtattacca atgtcagcaa attttctgtc ttcgaagagt
8101 aaaaaattgt acttggcgga taatgccttt agcggcttaa ctgtgccctc catggaaaaa
8161 tcagtcaaga tatccacatg tgtttttagt aaacaaattt tgggacctaa tgcttcaact
8221 aactccagta attccttggt ggtacgaaca tccaatgaag cacacaagtt tgtttgcttt
8281 tcgtgcatga tattaaatag cttggcagca acaggactag gatgagtagc agcacgttcc
8341 ttatatgtag ctttcgacat gatttatctt cgtttcctgc aggtttttgt tctgtgcagt
8401 tgggttaaga atactgggca atttcatgtt tcttcaacac tacatatgcg tatatatacc
8461 aatctaagtc tgtgctcctt ccttcgttct tccttctgtt cggagattac cgaatcaaaa
8521 aaatttcaaa gaaaccgaaa tcaaaaaaaa gaataaaaaa aaaatgatga attgaattga
8581 aaagctagct tatcgatggg tccttttcat cacgtgctat aaaaataatt ataatttaaa
8641 ttttttaata taaatatata aattaaaaat agaaagtaaa aaaagaaatt aaagaaaaaa
8701 tagtttttgt tttccgaaga tgtaaaagac tctaggggga tcgccaacaa atactacctt
8761 ttatcttgct cttcctgctc tcaggtatta atgccgaatt gtttcatctt gtctgtgtag
8821 aagaccacac acgaaaatcc tgtgatttta cattttactt atcgttaatc gaatgtatat
8881 ctatttaatc tgcttttctt gtctaataaa tatatatgta aagtacgctt tttgttgaaa
8941 ttttttaaac ctttgtttat tttttttttct tcattccgta actcttctac cttctttatt
9001 tactttctaa aatccaaata caaaacataa aaataaataa acacagagta aattcccaaa
9061 ttattccatc attaaaagat acgaggcgcg tgtaagttac aggcaagcga tctctaagaa
9121 accattatta tcatgacatt aacctataaa aataggcgta tcacgaggcc ctttcgtc

#### 8. pMQ30-ΔrcgAfecI2

**Description** Allelic-exchange construct for the in-frame deletion of rcgA (PA14_27700) and fecI2 (PA14_27690) in *P. aeruginosa* PA14. pMQ30 allelic-exchange (suicide) vector: Gm(R) aacC1, sacB counter-selection, oriT, URA3 + CEN6/ARSH4 for yeast gap-repair assembly (Shanks et al. 2006)

**Length** 9,183 bp, circular

**Source** This paper

**Note** *Deletion junctions retain 41 bp of PA14_27690 (4804..4844) and 49 bp of PA14_27700 (4845..4893).*

##### Features

| Location | Type | Feature |
|---|---|---|
| c(67..580) | misc_feature | CEN6/ARSH4; *S. cerevisiae* replicon from pYC2-CT |
| 819..1622 | gene | URA3 |
| 2058..2591 | gene | aacC1 (gentamicin resistance) |
| c(2855..3455) | rep_origin | ori (ColE1) |
| 4804..4844 | misc_feature | PA14_27690 (fecI2, RNA polymerase sigma factor) |
| 4845..4893 | misc_feature | PA14_27700 (rcgA, transcription regulator) |
| c(6256..7080) | misc_feature | oriT (conjugal transfer origin, from pEX18-Gm) |
| 7554..8975 | gene | sacB |

##### Sequence (9,183 bp)

```
   1 gacgaaaggg cctcgtgata cgcctatttt tataggttaa tgtcatgata ataatggttt
  61 cttagagatc gcttgcctgt aacttacacg cgcctcgtat cttttaatga tggaataatt
 121 tgggaattta ctctgtgttt atttattttt atgttttgta tttggatttt agaaagtaaa
 181 taaagaaggt agaagagtta cggaatgaag aaaaaaaaat aaacaaaggt ttaaaaaatt
 241 tcaacaaaaa gcgtacttta catatatatt tattagacaa gaaaagcaga ttaaatagat
 301 atacattcga ttaacgataa gtaaaatgta aaatcacagg attttcgtgt gtggtcttct
 361 acacagacaa gatgaaacaa ttcggcatta atacctgaga gcaggaagag caagataaaa
 421 ggtagtattt gttggcgatc cccctagagt cttttacatc ttcggaaaac aaaaactatt
 481 ttttctttaa tttctttttt tactttctat ttttaattta tatatttata ttaaaaaatt
 541 taaattataa ttatttttat agcacgtgat gaaaaggacc catcgataag ctagcttttc
 601 aattcaattc atcatttttt ttttattctt ttttttgatt tcggtttctt tgaaattttt
 661 ttgattcggt aatctccgaa cagaaggaag aacgaaggaa ggagcacaga cttagattgg
 721 tatatatacg catatgtagt gttgaagaaa catgaaattg cccagtattc ttaacccaac
 781 tgcacagaac aaaaacctgc aggaaacgaa gataaatcat gtcgaaagct acatataagg
 841 aacgtgctgc tactcatcct agtcctgttg ctgccaagct atttaatatc atgcacgaaa
 901 agcaaacaaa cttgtgtgct tcattggatg ttcgtaccac caaggaatta ctggagttag
 961 ttgaagcatt aggtcccaaa atttgtttac taaaaacaca tgtggatatc ttgactgatt
1021 tttccatgga gggcacagtt aagccgctaa aggcattatc cgccaagtac aatttttac
1081 tcttcgaaga cagaaaattt gctgacattg gtaatacagt caaattgcag tactctgcgg
1141 gtgtatacag aatagcagaa tgggcagaca ttacgaatgc acacggtgtg gtgggcccag
1201 gtattgttag cggtttgaag caggcggcag aagaagtaac aaaggaacct agaggccttt
1261 tgatgttagc agaattgtca tgcaagggct ccctatctac tggagaatat actaagggta
1321 ctgttgacat tgcgaagagc gacaaagatt ttgttatcgg ctttattgct caaagagaca
1381 tgggtggaag agatgaaggt tacgattggt tgattatgac acccggtgtg ggtttagatg
1441 acaagggaga cgcattgggt caacagtata gaaccgtgga tgatgtggtc tctacaggat
1501 ctgacattat tattgttgga agaggactat ttgcaaaggg aagggatgct aaggtagagg
1561 gtgaacgtta cagaaaagca ggctgggaag catatttgag aagatgcggc cagcaaaact
1621 aaaaaactgt attataagta aatgcatgta tactaaactc acaaattaga gcttcaattt
1681 aattatatca gttattaccc acttttcggg gaaatgtgcg cggaacccct atttgtttat
1741 ttttctaaat acattcaaat atgtatccgc tcatgagaca ataaccctga taaatgcttc
1801 aataatattg aaaaaggaag agtatgagta ttcaaccgaa ttgacataag cctgttcggt
1861 tcgtaaactg taatgcaagt agcgtatgcg ctcacgcaac tggtccagaa ccttgaccga
1921 acgcagcggt ggtaacggcg cagtggcggt tttcatggct tgttatgact gtttttttgt
1981 acagtctatg cctcgggcat ccaagcagca agcgcgttac gccgtgggtc gatgtttgat
```

2041 gttatggagc agcaacgatg ttacgcagca gcaacgatgt tacgcagcag ggcagtcgcc
2101 ctaaaacaaa gttaggtggc tcaagtatgg gcatcattcg cacatgtagg ctcggccctg
2161 accaagtcaa atccatgcgg gctgctcttg atcttttcgg tcgtgagttc ggagacgtag
2221 ccacctactc ccaacatcag ccggactccg attacctcgg gaacttgctc cgtagtaaga
2281 cattcatcgc gcttgctgcc ttcgaccaag aagcggttgt tggcgctctc gcggcttacg
2341 ttctgcccag gtttgagcag ccgcgtagtg agatctatat ctatgatctc gcagtctccg
2401 gcgagcaccg gaggcagggc attgccaccg cgctcatcaa tctcctcaag catgaggcca
2461 acgcgcttgg tgcttatgtg atctacgtgc aagcagatta cggtgacgat cccgcagtgg
2521 ctctctatac aaagttgggc atacgggaag aagtgatgca ctttgatatc gacccaagta
2581 ccgccaccta acaattcgtt caagccgaga tcggcttccc ggccgcggag ttgttcggta
2641 aattgtcaca acgccgcggc caattcgcat cgattgtcag accaagttta ctcatatata
2701 ctttagattg atttaaaact tcatttttaa tttaaaagga tctaggtgaa gatccttttt
2761 gataatctca tgaccaaaat cccttaacgt gagttttcgt tccactgagc gtcagacccc
2821 gtagaaaaga tcaaaggatc ttcttgagat cctttttttc tgcgcgtaat ctgctgcttg
2881 caaacaaaaa aaccaccgct accagcggtg gtttgtttgc cggatcaaga gctaccaact
2941 ctttttccga aggtaactgg cttcagcaga gcgcagatac caaatactgt ccttctagtg
3001 tagccgtagt taggccacca cttcaagaac tctgtagcac cgcctacata cctcgctctg
3061 ctaatcctgt taccagtggc tgctgccagt ggcgataagt cgtgtcttac cgggttggac
3121 tcaagacgat agttaccgga taaggcgcag cggtcgggct gaacgggggg ttcgtgcaca
3181 cagcccagct tggagcgaac gacctacacc gaactgagat acctacagcg tgagctatga
3241 gaaagcgcca cgcttcccga agggagaaag gcggacaggt atccggtaag cggcagggtc
3301 ggaacaggag agcgcacgag ggagcttcca gggggaaacg cctggtatct ttatagtcct
3361 gtcgggtttc gccacctctg acttgagcgt cgatttttgt gatgctcgtc aggggggcgg
3421 agcctatgga aaaacgccag caacgcggcc tttttacggt tcctggcctt ttgctggcct
3481 tttgctcaca tgttctttcc tgcgttatcc cctgattctg tggataaccg tattaccgcc
3541 tttgagtgag ctgataccgc tcgccgcagc cgaacgaccg agcgcagcga gtcagtgagc
3601 gaggaagcgg aagagcgccc aatacgcaaa ccgcctctcc ccgcgcgttg gccgattcat
3661 taatgcagct ggcacgacag gtttcccgac tggaaagcgg gcagtgagcg caacgcaatt
3721 aatgtgagtt agctcactca ttaggcaccc caggctttac actttatgct tccggctcgt
3781 atgttgtgtg gaattgtgag cggataacaa tttcacacag gaaacagctc tacgattggg
3841 tgtccttgcc gccgcgactg gcgagggtct cgctgtactt ggcgatgctc tggtcgatct
3901 tgtcgatggc gaagctggcc gattccgcct cgatcgcccc ggccgccgcg gctaccgcgg
3961 cggcggcgcc catcagcagg tcggtggcga gggcgccgag cgcgtcctcg gcagcctggg
4021 ccagcttgtt ttcggcgatg ttctgggtca tctgccggag caggatcgcc tggctggcca
4081 gggcgatctt gctgactccg tcgaagtagc tggcggcgga ttgcgccacc agcccggagg
4141 cctggctgat catcatgttc ggcgcgatgg cgatctgcga cggcgcatag ccggcggttt
4201 cgctgttgct catctgcacc gcctgcacga cttgactgtt cagggcgctg gtgtccggcg
4261 gcggcgcctc ggtctgcttg agcaggcgct cgctcatgcc ggccagcggc gaagggcgcg
4321 acggcagctt cacgtcgccg ttggcggacg ccctgccgcc gctggctggc gtctgccggc
4381 cggcctcgcc gggggcggcg gcttgctcgc tgtcgccgga tggcgcgggc gcgggcgcgg
4441 atttcttgcc gaagccgaac atgtctgggc actcctggga tcaaggggcg ggtcagatac
4501 ctttcttgtc ttctatcgcc atgatcatcg acaccgcctt ggaaaccacc gcggtggaga
4561 tctggttcag ggcctgctgg ctgtagactg cgttctgcag ggacaggccg gcggaatggc
4621 tggcgatctg gtagagcgcg gcgatggcct gggccggggc ttcggcgacc accttgacgt
4681 tggtctgggt gacggcatcg gtgatctggg cattcacagc cgtggggtca gccatgggac
4741 tctccttgaa tggcaggtg cttgtcgggc agggggccgg tcgcacgggc ctcgactcgt
4801 ttgtcaggaa tgaacgcggc tgcgtctgtc ttgtgaaacg tcgcgggtgt agtggcgtga
4861 acagcggtct tccgttgact cgttcgagtg catgcctgcg ccttcgccag attgatcggc
4921 agctccggcg aggtacgaca tgctcgctgc agagcaatca gtggggacgt gtgcggcgaa
4981 ggggtgtgaa gctccggcta cccggcagaa ctgctaggta cggactcggc gcgctccggc
5041 aatcgggcgg gcggtccgtt ttttattcgg aaagggtagc gaggtttggg ctggcgggag
5101 gcggccccga gcgcacggat ggcggtcggg gcgggacggc cggtggccgt ggcgattcag
5161 tgccgcgggc tgtcctcggg cagcgcgagc aactgctttt cgcggttcca gtcgaacgtt
5221 tcgccgcgct cttcggcttc gtagcggcgc tcgtcgaggc gctggaacag gtcgatctct
5281 tcgtccggca tgaagtgcag gcaatcgccg ccgaagaacc acagcaggtc gcgcggtacc
5341 aggtgggcga tctgcggata gcggtggaag acctgggaaa tcaggtcctg gccgaggtat
5401 tgggcgcctt ccgggtcgcg cggcagttcg tccatcagtt cgtcgaaacg ctcgaggaac
5461 agcccgtggt ttcttccgg cacctgctcg gcctcgccca gggcgccgag gatgttgcgc
5521 aggtgctgga gcagggcgag gtggtgctcg aggtaggagt tggccatgac gggtcctctg
5581 gaaaaaagct gggcagtata aagggtttcg cggcgccggc cagtttcagc ggatcgccgg
5641 ggactggctc tggcgccggc ggttttccag ttgccggcgt tcgctgtcga ggcgccgctg
5701 gtcctgctcc aggcgccagc gctgctggtc cagttgctgc tggcggatca ggttgtcctg
5761 catctgtcgc tgctgccgtt gccgctgcag gttctgctgc tggcgttgca gttgatcctg
5821 ctgcagttgc aggcgttgtt cgtatcagaa cgcagaagcg gtctgataaa acagaatttg
5881 cctggcggca gtagcgcggt ggtcccacct gaccccatgc cgaactcaga agtgaaacgc
5941 cgtagcgccg atggtagtgt ggggtctccc catgcgagag tagggaactg ccaggcatca
6001 aataaaacga aaggctcagt cgaaagactg ggcctttcgt tttatctgtt gtttgtcggt

6061 gaacgctctc ctgagtagga caaatccgcc gggagcggat ttgaacgttg cgaagcaacg
6121 gcccggaggg tggcgggcag gacgcccgcc ataaactgcc aggcatcaaa ttaagcagaa
6181 ggccatcctg acggatggcc tttttgcgtt tctacaaact ctttttgttt atttttctaa
6241 atacattcaa atatgcgcct gatgcggtat tttctcctta cgcatatcga catccgccct
6301 caccgccagg aacgcaaccg cagcctcatc acgccggcgc ttcttggccg cgcgggattc
6361 aacccactcg gccagctcgt cggtgtagct ctttggcatc gtctctcgcc tgtcccctca
6421 gttcagtaat ttcctgcatt tgcctgtttc cagtcggtag atattccaca aaacagcagg
6481 gaagcagcgc ttttccgctg cataaccctg cttcggggtc attatagcga ttttttcggt
6541 atatccatcc tttttcgcac gatatacagg attttgccaa agggttcgtg tagactttcc
6601 ttggtgtatc caacggcgtc agccgggcag gataggtgaa gtaggcccac ccgcgagcgg
6661 gtgttccttc ttcactgtcc cttattcgca cctggcggtg ctcaacggga atcctgctct
6721 gcgaggctgg ccggctaccg ccggcgtaac agatgagggc aagcggatgg ctgatgaaac
6781 caagccaacc aggaagggca gcccacctat caaggtgtac tgccttccag acgaacgaag
6841 agcgattgag gaaaaggcgg cggcggccgg catgagcctg tcggcctacc tgctggccgt
6901 cggccagggc tacaaaatca cgggcgtcgt ggactatgag cacgtccgcg agctggcccg
6961 catcaatggc gacctgggcc gcctgggcgg cctgctgaaa ctctggctca ccgacgaccc
7021 gcgcacggcg cggttcggtg atgccacgat cctcgccctg ctggcgaaga tcgactctag
7081 ctagaggatc gatccttttt aacccatcac atatacctgc cgttcactat tatttagtga
7141 aatgagatat tatgatattt tctgaattgt gattaaaaag gcaactttat gcccatgcaa
7201 cagaaactat aaaaaataca gagaatgaaa agaaacagat agatttttta gttctttagg
7261 cccgtagtct gcaaatcctt ttatgatttt ctatcaaaca aaagaggaaa atagaccagt
7321 tgcaatccaa acgagagtct aatagaatga ggtcgaaaag taaatcgcgc gggtttgtta
7381 ctgataaagc aggcaagacc taaaatgtgt aaagggcaaa gtgtatactt tggcgtcacc
7441 ccttacatat tttaggtctt tttttattgt gcgtaactaa cttgccatct tcaaacagga
7501 gggctggaag aagcagaccg ctaacacagt acataaaaaa ggagacatga acgatgaaca
7561 tcaaaaagtt tgcaaaacaa gcaacagtat taacctttac taccgcactg ctggcaggag
7621 gcgcaactca agcgtttgcg aaagaaacga accaaaagcc atataaggaa acatacggca
7681 tttcccatat tacacgccat gatatgctgc aaatccctga acagcaaaaa aatgaaaaat
7741 atcaagttcc tgagttcgat tcgtccacaa ttaaaaatat ctcttctgca aaaggcctgg
7801 acgtttggga cagctggcca ttacaaaacg ctgacggcac tgtcgcaaac tatcacggct
7861 accacatcgt ctttgcatta gccggagatc ctaaaaatgc ggatgacaca tcgatttaca
7921 tgttctatca aaaagtcggc gaaacttcta ttgacagctg gaaaaacgct ggccgcgtct
7981 ttaaagacag cgacaaattc gatgcaaatg attctatcct aaaagaccaa acacaagaat
8041 ggtcaggttc agccacattt acatctgacg gaaaaatccg tttattctac actgatttct
8101 ccggtaaaca ttacggcaaa caaacactga caactgcaca agttaacgta tcagcatcag
8161 acagctcttt gaacatcaac ggtgtagagg attataaatc aatctttgac ggtgacggaa
8221 aaacgtatca aaatgtacag cagttcatcg atgaaggcaa ctacagctca ggcgacaacc
8281 atacgctgag agatcctcac tacgtagaag ataaaggcca caaatactta gtatttgaag
8341 caaacactgg aactgaagat ggctaccaag gcgaagaatc tttatttaac aaagcatact
8401 atggcaaaag cacatcattc ttccgtcaag aaagtcaaaa acttctgcaa agcgataaaa
8461 aacgcacggc tgagttagca aacggcgctc tcggtatgat tgagctaaac gatgattaca
8521 cactgaaaaa agtgatgaaa ccgctgattg catctaacac agtaacagat gaaattgaac
8581 gcgcgaacgt ctttaaaatg aacggcaaat ggtatctgtt cactgactcc cgcggatcaa
8641 aaatgacgat tgacggcatt acgtctaacg atatttacat gcttggttat gtttctaatt
8701 ctttaactgg cccatacaag ccgctgaaca aaactggcct tgtgttaaaa atggatcttg
8761 atcctaacga tgtaaccttt acttactcac acttcgctgt acctcaagcg aaaggaaaca
8821 atgtcgtgat tacaagctat atgacaaaca gaggattcta cgcagacaaa caatcaacgt
8881 ttgcgcctag cttcctgctg aacatcaaag gcaagaaaac atctgttgtc aaagacagca
8941 tccttgaaca aggacaatta acagttaaca aataaaaacg caaaagaaaa tgccgattat
9001 ggtgcactct cagtacaatc tgctctgatg ccgcatagtt aagccagccc cgacacccgc
9061 caacacccgc tgacgcgccc tgacgggctt gtctgctccc ggcatccgct tacagacaag
9121 ctgtgaccgt ctccgggagc tgcatgtgtc agaggttttc accgtcatca ccgaaacgcg
9181 cga

#### 9. pMQ30-H22A-27675

**Description** Allelic-exchange construct introducing the H22A point mutation at the native PA14_27675 locus in *P. aeruginosa* PA14. pMQ30 allelic-exchange (suicide) vector: Gm(R) aacC1, sacB counter-selection, oriT, URA3 + CEN6/ARSH4 for yeast gap-repair assembly (Shanks et al. 2006)

**Length** 11,373 bp, circular

**Source** This paper

**Note** *H22A codon at 4432..4434 (GCC); the allele is otherwise identical to wild type.*

##### Features

| Location | Type | Feature |
|---|---|---|
| c(209..1630) | gene | sacB |
| 2104..2928 | misc_feature | oriT (conjugal transfer origin, from pEX18-Gm) |
| 3686..4318 | misc_feature | PA14_27680 (rapA) |
| 4369..6510 | CDS | PA14_27675 (dear), H22A, 713 aa |
| 4432..4434 | misc_feature | H22A codon (GCC) |
| 6521..6715 | misc_feature | PA14_27670 |
| 6739..6978 | misc_feature | PA14_27660 |
| c(7562..7584) | misc_binding | lac operator |
| c(7588..7618) | promoter | lac promoter |
| 7919..8519 | rep_origin | ori (ColE1) |
| c(8783..9316) | gene | aacC1 (gentamicin resistance) |
| c(9674..10781) | gene | URA3 |
| 10794..11307 | misc_feature | CEN6/ARSH4; *S. cerevisiae* replicon from pYC2-CT |

##### Sequence (11,373 bp)

```
   1 tcgcgcgttt cggtgatgac ggtgaaaacc tctgacacat gcagctcccg gagacggtca
  61 cagcttgtct gtaagcggat gccgggagca gacaagcccg tcagggcgcg tcagcgggtg
 121 ttggcgggtg tcggggctgg cttaactatg cggcatcaga gcagattgta ctgagagtgc
 181 accataatcg gcattttctt ttgcgttttt atttgttaac tgttaattgt ccttgttcaa
 241 ggatgctgtc tttgacaaca gatgttttct tgcctttgat gttcagcagg aagctaggcg
 301 caaacgttga ttgtttgtct gcgtagaatc ctctgtttgt catatagctt gtaatcacga
 361 cattgtttcc tttcgcttga ggtacagcga agtgtgagta agtaaaggtt acatcgttag
 421 gatcaagatc catttttaac acaaggccag ttttgttcag cggcttgtat gggccagtta
 481 aagaattaga aacataacca agcatgtaaa tatcgttaga cgtaatgccg tcaatcgtca
 541 tttttgatcc gcgggagtca gtgaacagat accatttgcc gttcatttta aagacgttcg
 601 cgcgttcaat ttcatctgtt actgtgttag atgcaatcag cggtttcatc acttttttca
 661 gtgtgtaatc atcgtttagc tcaatcatac cgagagcgcc gtttgctaac tcagccgtgc
 721 gtttttttatc gctttgcaga agtttttgac tttcttgacg gaagaatgat gtgcttttgc
 781 catagtatgc tttgttaaat aaagattctt cgccttggta gccatcttca gttccagtgt
 841 ttgcttcaaa tactaagtat ttgtggcctt tatcttctac gtagtgagga tctctcagcg
 901 tatggttgtc gcctgagctg tagttgcctt catcgatgaa ctgctgtaca ttttgatacg
 961 tttttccgtc accgtcaaag attgatttat aatcctctac accgttgatg ttcaaagagc
1021 tgtctgatgc tgatacgtta acttgtgcag ttgtcagtgt ttgtttgccg taatgtttac
1081 cggagaaatc agtgtagaat aaacggattt ttccgtcaga tgtaaatgtg gctgaacctg
1141 accattcttg tgtttggtct tttaggatag aatcatttgc atcgaatttg tcgctgtctt
1201 taaagacgcg gccagcgttt ttccagctgt caatagaagt ttcgccgact ttttgataga
1261 acatgtaaat cgatgtgtca tccgcatttt taggatctcc ggctaatgca aagacgatgt
1321 ggtagccgtg atagtttgcg acagtgccgt cagcgttttg taatggccag ctgtcccaaa
1381 cgtccaggcc ttttgcagaa gagatatttt taattgtgga cgaatcgaac tcaggaactt
1441 gatatttttc atttttttgc tgttcaggga tttgcagcat atcatggcgt gtaatatggg
```

1501 aaatgccgta tgtttcctta tatggctttt ggttcgtttc tttcgcaaac gcttgagttg
1561 cgcctcctgc cagcagtgcg gtagtaaagg ttaatactgt tgcttgtttt gcaaactttt
1621 tgatgttcat cgttcatgtc tcctttttta tgtactgtgt tagcggtctg cttcttccag
1681 ccctcctgtt tgaagatggc aagttagtta cgcacaataa aaaaagacct aaaatatgta
1741 aggggtgacg ccaaagtata cactttgccc tttacacatt ttaggtcttg cctgctttat
1801 cagtaacaaa cccgcgcgat ttacttttcg acctcattct attagactct cgtttggatt
1861 gcaactggtc tattttcctc ttttgtttga tagaaaatca taaaaggatt tgcagactac
1921 gggcctaaag aactaaaaaa tctatctgtt tcttttcatt ctctgtattt tttatagttt
1981 ctgttgcatg ggcataaagt tgcctttttta atcacaattc agaaaatatc ataatatctc
2041 atttcactaa ataatagtga acggcaggta tatgtgatgg gttaaaaagg atcgatcctc
2101 tagctagagt cgatcttcgc cagcagggcg aggatcgtgg catcaccgaa ccgcgccgtg
2161 cgcgggtcgt cggtgagcca gagtttcagc aggccgccca ggcggcccag gtcgccattg
2221 atgcgggcca gctcgcggac gtgctcatag tccacgacgc ccgtgatttt gtagccctgg
2281 ccgacggcca gcaggtaggc cgacaggctc atgccggccg ccgccgcctt ttcctcaatc
2341 gctcttcgtt cgtctggaag gcagtacacc ttgataggtg ggctgccctt cctggttggc
2401 ttggtttcat cagccatccg cttgccctca tctgttacgc cggcggtagc cggccagcct
2461 cgcagagcag gattcccgtt gagcaccgcc aggtgcgaat aagggacagt gaagaaggaa
2521 cacccgctcg cgggtgggcc tacttcacct atcctgcccg gctgacgccg ttggatacac
2581 caaggaaagt ctacacgaac cctttggcaa aatcctgtat atcgtgcgaa aaaggatgga
2641 tataccgaaa aaatcgctat aatgaccccg aagcagggtt atgcagcgga aaagcgctgc
2701 ttccctgctg ttttgtggaa tatctaccga ctggaaacag gcaaatgcag gaaattactg
2761 aactgagggg acaggcgaga gacgatgcca aagagctaca ccgacgagct ggccgagtgg
2821 gttgaatccc gcgcggccaa gaagcgccgg cgtgatgagg ctgcggttgc gttcctggcg
2881 gtgagggcgg atgtcgatat gcgtaaggag aaaataccgc atcaggcgca tatttgaatg
2941 tatttagaaa aataaacaaa aagagtttgt agaaacgcaa aaaggccatc cgtcaggatg
3001 gccttctgct taatttgatg cctggcagtt tatggcgggc gtcctgcccg ccaccctccg
3061 ggccgttgct tcgcaacgtt caaatccgct cccggcggat ttgtcctact caggagagcg
3121 ttcaccgaca aacaacagat aaaacgaaag gcccagtctt tcgactgagc ctttcgtttt
3181 atttgatgcc tggcagttcc ctactctcgc atggggagac cccacactac catcggcgct
3241 acggcgtttc acttctgagt tcggcatggg gtcaggtggg accaccgcgc tactgccgcc
3301 aggcaaattc tgttttatca gaccgcttct gcgttctgat ctgacaaacg agtcgaggcc
3361 cgtgcgaccg gccccctgcc cgacaagcac ctgcccattc aaggagagtc ccatggctga
3421 ccccacggct gtgaatgccc agatcaccga tgccgtcacc cagaccaacg tcaaggtggt
3481 cgcccgaagcc ccggcccagg ccatcgccgc gctctaccag atcgccagcc attccgccgg
3541 cctgtccctg cagaacgcag tctacagcca gcaggccctg aaccagatct ccaccgcggt
3601 ggtttccaag gcggtgtcga tgatcatggc gatagaagac aagaaaggta tctgacccgc
3661 cccttgatcc caggagtgcc cagacatgtt cggcttcggc aagaaatccg cgcccgcgcc
3721 cgcgccatcc ggcgacagcg agcaagccgc cgccccggc gaggccggcc ggcagacgcc
3781 agccagcggc ggcagggcgt ccgccaacgg cgacgtgaag ctgccgtcgc gcccttcgcc
3841 gctggccggc atgagcgagc gcctgctcaa gcagaccgag gcgccgccgc cggacaccag
3901 cgccctgaac agtcaagtcg tgcaggcggt gcagatgagc aacagcgaaa ccgccggcta
3961 tgcgccgtcg cagatcgcca tcgcgccgaa catgatgatc agccaggcct ccgggctggt
4021 ggcgcaatcc gccgccagct acttcgacgg agtcagcaag atcgccctgg ccagccaggc
4081 gatcctgctc cggcagatga cccagaacat cgccgaaaac aagctggccc aggctgccga
4141 ggacgcgctc ggcgccctcg ccaccgacct gctgatgggc gccgccgccg cggtagccgc
4201 ggcggccggg gcgatcgagg cggaatcggc cagcttcgcc atcgacaaga tcgaccagag
4261 catcgccaag tacagcgaga ccctcgccag tcgcggcggc aaggacaccc aatcgtagcc
4321 accaccggcg gggccggcgg cgtccggccc tggcacagga gagccgtcat ggcctatttc
4381 aagctggaag aaccggtcag gttccatcgc tatcccttcg acttccatag tgccttcgcc
4441 ggcatcctgc cggtggaaag caacagccgc tggacgcgcg accgccgggt cttccgcgtc
4501 ggcgagcggc aagtgagcct ggagaaaggc caggagcttt ccctgatcgg gctgctgatg
4561 agtgcccgcg gcgtcgcccc ggacgtcgac ggcaaggcgc tggaggaggc tcgccaggcc
4621 gcgcactatg agctgttcga actggctctg cagcggatgg tcaggcgcaa tccgttcgcc
4681 gcgaccgacc gtcagggcta cctgcgcggc gaatgcgcgg cggagaacat ctatctcgcc
4741 tgcctcatcc tggcgcaacg ttttggccgg acctcgccgc cagcggccat cgaccagcca
4801 gcgatctacc tcggcaccct ggaactgctc ggtgcgagcg cggttcgcga cagcgagacg
4861 gaccagttcg tccgctactt caaccgcaag atctggagcg gcaacaagta cacgccgttc
4921 gacgacgcct actgggcccg cggcgccatc cgcgatcgcc atccgggcga attcgcctgc
4981 ctgacgctgg ggttcctcct ccacgaagga atcagccata cccagacggc gacgggggaa
5041 gacgaggtgg ccgtcctcga ttccctgttc gaacaattca acgcctccga gaagaccgcc
5101 taccgcctgc tcgcccacac cgcgcatggc tatgccagcg aagccgcctt cgatgccgag
5161 ctgcacagga tcctccgtca tttcgaaata cagcaggggc agccaccgca ggcccggctg
5221 gtcggcatcg acctgctcgg catggagacc gctaccggcc tgtaccgaca gttcttcgac
5281 ttcctcctcg gacaggccgc ggtgtttcgc cactacctgg acggcaagcc ggaaacccgc
5341 aagatggtcc tgcacatcca ttgtggagag gggaccgggg tgtccgacga caaccggtcg
5401 ctgtgcggtt acttcctgcg caacgccaac gcgctggacg atttctacgc agcgctctcc
5461 gcctatgcct ggaagtgcta tggcaacacc atccgccagg gcaaggcgcg gctacgcgaa

5521 cgggagaacc tgcaggaacg tgacaaggcg cccagcgccc tcgccggcct tttcgacgaa
5581 ctgtttttcg gcaacagcct gaccgcctcc ggcctgcggc tacgccgctt cgacatcacc
5641 agcgggacta cccaggccct ggtcgcctat tacgcccgca ccaacgtcgt gaacctctgc
5701 caggcgctgg ccagccgcga cgccgacggc aacagctact accgccgcct gctggagagc
5761 gatctgttca gcctgcgcat cggccatgcc tactactacc gcaactatct ggccagcaag
5821 ttccccgagc tgtgcttcga caccaacctg ggaagcaatt tcattaccgg cgcctccggc
5881 ctcttcgact cgctgcagga ataccgcctg aaccgtggcc tgcgccatct cgatggatac
5941 gtcggcaccg accagctcaa ggaactcagt ctggccatcg cctaccaggg cgaacagcgc
6001 ctggacccac aacagatgca gtacgtccat gccctggcgg aaagccagag cggcttcgac
6061 gaactgggcg aacacctccc gggtacaccc ggctgggcga aacccgcgct ggaacagttc
6121 ttcgtaagtc agtgcgcgct ctaccgctcc gaggaggatc gctacttcca gttcgaagcc
6181 taccggcgcc tcttcgccca ggtcctcaac tggcgctcct atctgctggg cgccgacggc
6241 cagggtgtcg agcactccaa cgtacaggac gaagcgatcc gcatggcgct cctgctcaac
6301 tatgccgccg cggaccgcca cggccgggtt ccggccgctt cgctggagaa tgcccaacgc
6361 ctgctggtgc aactgggcag cgcctactgg gaggaaacca tcggcgccgt cgacctggcc
6421 ggcgcgcccc atcgggaccg agagctgcag cgcttcgaag gattcgccgc ccccgcctcg
6481 gtcgtgcgca tctccacccg cagcagttga ggagcgaccg atgagcgcct acgcacgcta
6541 catcagacgc cgcgcgctcg ccgccgcgga ccgcccggcg cagcgttcgc cggacgccag
6601 cgaggaggca cgggaccaac cgccccccag caggccggcg agcggaaagc gcctgctgga
6661 aatatcggac agcgatgcgg ccaagctctt ggccgtgctg aaactgctcg gatgagcacc
6721 caccccagga gaagcgtcat gaaggaagaa ctcgaaaccc tcgtcaccca ggtcaaggca
6781 ctcctcaccg aatgcctgca aggctggctc caggccctgt cccagggaga cagcgtggac
6841 gacccggcca gcctcggaca gttggcggtg cagttgagca ccgcctacag caccctgcag
6901 gccgctgccg ccgcctaccc cggcgatacc ggcagcagcg cgcaggcccc ctcggcggac
6961 ggcacgcagc agtactgaag cctcgccacg acaccccgcg caccgcgcgt ccatcgcaga
7021 ggagtccgcg catgtcgatc tcggaagaat tgggaaggaa tctggtccag ctggcgcaac
7081 gctggctgaa acgccccctc accagcgaag aacacgagca gttgagacag ttccagctcg
7141 gcctcggcaa cgacgccagc agcgatccgg ccgtgcacgc gcgacagcag gccgaacagg
7201 tgatcgagaa cggccggcag cgttccgaag gctccatccg ccagatcctg cagaacgtcc
7261 agcaggcctc gacccaggcc ctccgcgccc aggacagcgg cgagcaggcg attctcgagg
7321 cggtcaatgc cgccggcagc ctggccgaac tgcgtcccgg ccacctgcag agcggccagg
7381 caagcgccaa ccagttggtg atgtcgcaga ttgccgaccg cctggccaac ctagtgaaga
7441 gcgaggtacg cctgtgcttc gagcagaact tcggctcgct caagcagcaa ctggagaacg
7501 ctatcgccat cttcaaccag agcaaccaca acacgtccgc accgagctgt ttcctgtgtg
7561 aaattgttat ccgctcacaa ttccacacaa catacgagcc ggaagcataa agtgtaaagc
7621 ctggggtgcc taatgagtga gctaactcac attaattgcg ttgcgctcac tgcccgcttt
7681 ccagtcggga aacctgtcgt gccagctgca ttaatgaatc ggccaacgcg cggggagagg
7741 cggtttgcgt attgggcgct cttccgcttc ctcgctcact gactcgctgc gctcggtcgt
7801 tcggctgcgg cgagcggtat cagctcactc aaaggcggta atacggttat ccacagaatc
7861 aggggataac gcaggaaaga acatgtgagc aaaaggccag caaaaggcca ggaaccgtaa
7921 aaaggccgcg ttgctggcgt ttttccatag gctccgcccc cctgacgagc atcacaaaaa
7981 tcgacgctca agtcagaggt ggcgaaaccc gacaggacta taaagatacc aggcgtttcc
8041 ccctggaagc tccctcgtgc gctctcctgt tccgaccctg ccgcttaccg gatacctgtc
8101 cgcctttctc ccttcgggaa gcgtggcgct ttctcatagc tcacgctgta ggtatctcag
8161 ttcggtgtag gtcgttcgct ccaagctggg ctgtgtgcac gaaccccccg ttcagcccga
8221 ccgctgcgcc ttatccggta actatcgtct tgagtccaac ccggtaagac acgacttatc
8281 gccactggca gcagccactg gtaacaggat tagcagagcg aggtatgtag gcggtgctac
8341 agagttcttg aagtggtggc ctaactacgg ctacactaga aggacagtat ttggtatctg
8401 cgctctgctg aagccagtta ccttcggaaa aagagttggt agctcttgat ccggcaaaca
8461 aaccaccgct ggtagcggtg gtttttttgt ttgcaagcag cagattacgc gcagaaaaaa
8521 aggatctcaa gaagatcctt tgatcttttc tacggggtct gacgctcagt ggaacgaaaa
8581 ctcacgttaa gggattttgg tcatgagatt atcaaaaagg atcttcacct agatcctttt
8641 aaattaaaaa tgaagtttta aatcaatcta aagtatatat gagtaaactt ggtctgacaa
8701 tcgatgcgaa ttggccgcgg cgttgtgaca atttaccgaa caactccgcg gccgggaagc
8761 cgatctcggc ttgaacgaat tgttaggtgg cggtacttgg gtcgatatca aagtgcatca
8821 cttcttcccg tatgcccaac tttgtataga gagccactgc gggatcgtca ccgtaatctg
8881 cttgcacgta gatcacataa gcaccaagcg cgttggcctc atgcttgagg agattgatga
8941 gcgcggtggc aatgccctgc ctccggtgct cgccggagac tgcgagatca tagatataga
9001 tctcactacg cggctgctca aacctgggca gaacgtaagc cgcgagagcg ccaacaaccg
9061 cttcttggtc gaaggcagca agcgcgatga atgtcttact acggagcaag ttcccgaggt
9121 aatcggagtc cggctgatgt tgggagtagg tggctacgtc tccgaactca cgaccgaaaa
9181 gatcaagagc agcccgcatg gatttgactt ggtcagggcc gagcctacat gtgcgaatga
9241 tgcccatact tgagccacct aactttgttt tagggcgact gccctgctgc gtaacatcgt
9301 tgctgctgcg taacatcgtt gctgctccat aacatcaaac atcgacccac ggcgtaacgc
9361 gcttgctgct tggatgcccg aggcatagac tgtacaaaaa aacagtcata acaagccatg
9421 aaaaccgcca ctgcgccgtt accaccgctg cgttcggtca aggttctgga ccagttgcgt
9481 gagcgcatac gctacttgca ttacagttta cgaaccgaac aggcttatgt caattcggtt

9541 gaatactcat actcttcctt tttcaatatt attgaagcat ttatcagggt tattgtctca
9601 tgagcggata catatttgaa tgtatttaga aaaataaaca aataggggtt ccgcgcacat
9661 ttccccgaaa agtgggtaat aactgatata attaaattga agctctaatt tgtgagttta
9721 gtatacatgc atttacttat aatacagttt tttagttttg ctggccgcat cttctcaaat
9781 atgcttccca gcctgctttt ctgtaacgtt caccctctac cttagcatcc cttccctttg
9841 caaatagtcc tcttccaaca ataataatgt cagatcctgt agagaccaca tcatccacgg
9901 ttctatactg ttgacccaat gcgtctccct tgtcatctaa acccacaccg ggtgtcataa
9961 tcaaccaatc gtaaccttca tctcttccac ccatgtctct ttgagcaata aagccgataa
10021 caaaatcttt gtcgctcttc gcaatgtcaa cagtaccctt agtatattct ccagtagata
10081 gggagccctt gcatgacaat tctgctaaca tcaaaaggcc tctaggttcc tttgttactt
10141 cttctgccgc ctgcttcaaa ccgctaacaa tacctgggcc caccacaccg tgtgcattcg
10201 taatgtctgc ccattctgct attctgtata cacccgcaga gtactgcaat ttgactgtat
10261 taccaatgtc agcaaatttt ctgtcttcga agagtaaaaa attgtacttg gcggataatg
10321 cctttagcgg cttaactgtg ccctccatgg aaaaatcagt caagatatcc acatgtgttt
10381 ttagtaaaca aattttggga cctaatgctt caactaactc cagtaattcc ttggtggtac
10441 gaacatccaa tgaagcacac aagtttgttt gcttttcgtg catgatatta aatagcttgg
10501 cagcaacagg actaggatga gtagcagcac gttccttata tgtagctttc gacatgattt
10561 atcttcgttt cctgcaggtt tttgttctgt gcagttgggt taagaatact gggcaatttc
10621 atgtttcttc aacactacat atgcgtatat ataccaatct aagtctgtgc tccttccttc
10681 gttcttcctt ctgttcggag attaccgaat caaaaaaatt tcaaagaaac cgaaatcaaa
10741 aaaaagaata aaaaaaaaat gatgaattga attgaaaagc tagcttatcg atgggtcctt
10801 ttcatcacgt gctataaaaa taattataat ttaaattttt taatataaat atataaatta
10861 aaaatagaaa gtaaaaaaag aaattaaaga aaaaatagtt tttgttttcc gaagatgtaa
10921 aagactctag ggggatcgcc aacaaatact accttttatc ttgctcttcc tgctctcagg
10981 tattaatgcc gaattgtttc atcttgtctg tgtagaagac cacacacgaa aatcctgtga
11041 ttttacattt tacttatcgt taatcgaatg tatatctatt taatctgctt ttcttgtcta
11101 ataaatatat atgtaaagta cgctttttgt tgaaattttt taaacctttg tttatttttt
11161 tttcttcatt ccgtaactct tctaccttct ttatttactt tctaaaatcc aaatacaaaa
11221 cataaaaata aataaacaca gagtaaattc ccaaattatt ccatcattaa aagatacgag
11281 gcgcgtgtaa gttacaggca agcgatctct aagaaaccat tattatcatg acattaacct
11341 ataaaaatag gcgtatcacg aggccctttc gtc

#### 10. pMQ30-Δ27675::27675

**Description** Allelic-exchange construct restoring wild-type PA14_27675 at the native locus (complementation of the Δ27675 strain). pMQ30 allelic-exchange (suicide) vector: Gm(R) aacC1, sacB counter-selection, oriT, URA3 + CEN6/ARSH4 for yeast gap-repair assembly (Shanks et al. 2006)

**Length** 11,286 bp, circular

**Source** This paper

##### Features

| Location | Type | Feature |
|---|---|---|
| c(209..1630) | gene | sacB |
| 2104..2928 | misc_feature | oriT (conjugal transfer origin, from pEX18-Gm) |
| 3636..4268 | misc_feature | PA14_27680 (rapA) |
| 4319..6460 | CDS | PA14_27675 (dear), wild type, 713 aa |
| 6471..6665 | misc_feature | PA14_27670 |
| 6689..6928 | misc_feature | PA14_27660 |
| c(7475..7497) | misc_binding | lac operator |
| c(7501..7531) | promoter | lac promoter |
| 7832..8432 | rep_origin | ori (ColE1) |
| c(8696..9229) | gene | aacC1 (gentamicin resistance) |
| c(9587..10694) | gene | URA3 |
| 10707..11220 | misc_feature | CEN6/ARSH4; *S. cerevisiae* replicon from pYC2-CT |

##### Sequence (11,286 bp)

```
   1 tcgcgcgttt cggtgatgac ggtgaaaacc tctgacacat gcagctcccg gagacggtca
  61 cagcttgtct gtaagcggat gccgggagca gacaagcccg tcagggcgcg tcagcgggtg
 121 ttggcgggtg tcggggctgg cttaactatg cggcatcaga gcagattgta ctgagagtgc
 181 accataatcg gcattttctt ttgcgttttt atttgttaac tgttaattgt ccttgttcaa
 241 ggatgctgtc tttgacaaca gatgttttct tgcctttgat gttcagcagg aagctaggcg
 301 caaacgttga ttgtttgtct gcgtagaatc ctctgtttgt catatagctt gtaatcacga
 361 cattgtttcc tttcgcttga ggtacagcga agtgtgagta agtaaaggtt acatcgttag
 421 gatcaagatc catttttaac acaaggccag ttttgttcag cggcttgtat gggccagtta
 481 aagaattaga aacataacca agcatgtaaa tatcgttaga cgtaatgccg tcaatcgtca
 541 tttttgatcc gcgggagtca gtgaacagat accatttgcc gttcatttta aagacgttcg
 601 cgcgttcaat ttcatctgtt actgtgttag atgcaatcag cggtttcatc actttttca
 661 gtgtgtaatc atcgtttagc tcaatcatac cgagagcgcc gtttgctaac tcagccgtgc
 721 gtttttatc gctttgcaga agtttttgac tttcttgacg gaagaatgat gtgcttttgc
 781 catagtatgc tttgttaaat aaagattctt cgccttggta gccatcttca gttccagtgt
 841 ttgcttcaaa tactaagtat ttgtggcctt tatcttctac gtagtgagga tctctcagcg
 901 tatggttgtc gcctgagctg tagttgcctt catcgatgaa ctgctgtaca ttttgatacg
 961 tttttccgtc accgtcaaag attgatttat aatcctctac accgttgatg ttcaaagagc
1021 tgtctgatgc tgatacgtta acttgtgcag ttgtcagtgt ttgtttgccg taatgtttac
1081 cggagaaatc agtgtagaat aaacggattt ttccgtcaga tgtaaatgtg gctgaacctg
1141 accattcttg tgtttggtct tttaggatag aatcatttgc atcgaatttg tcgctgtctt
1201 taaagacgcg gccagcgttt ttccagctgt caatagaagt ttcgccgact ttttgataga
1261 acatgtaaat cgatgtgtca tccgcatttt taggatctcc ggctaatgca aagacgatgt
1321 ggtagccgtg atagtttgcg acagtgccgt cagcgttttg taatggccag ctgtcccaaa
1381 cgtccaggcc ttttgcagaa gagatatttt taattgtgga cgaatcgaac tcaggaactt
1441 gatatttttc atttttttgc tgttcaggga tttgcagcat atcatggcgt gtaatatggg
1501 aaatgccgta tgtttcctta tatggctttt ggttcgtttc tttcgcaaac gcttgagttg
1561 cgcctcctgc cagcagtgcg gtagtaaagg ttaatactgt tgcttgtttt gcaaactttt
1621 tgatgttcat cgttcatgtc tccttttta tgtactgtgt tagcggtctg cttcttccag
1681 ccctcctgtt tgaagatggc aagttagtta cgcacaataa aaaaagacct aaaatatgta
```

1741 aggggtgacg ccaaagtata cactttgccc tttacacatt ttaggtcttg cctgctttat
1801 cagtaacaaa cccgcgcgat ttacttttcg acctcattct attagactct cgtttggatt
1861 gcaactggtc tattttcctc ttttgtttga tagaaaatca taaaaggatt tgcagactac
1921 gggcctaaag aactaaaaaa tctatctgtt tcttttcatt ctctgtattt tttatagttt
1981 ctgttgcatg ggcataaagt tgccttttta atcacaattc agaaaatatc ataatatctc
2041 atttcactaa ataatagtga acggcaggta tatgtgatgg gttaaaaagg atcgatcctc
2101 tagctagagt cgatcttcgc cagcagggcg aggatcgtgg catcaccgaa ccgcgccgtg
2161 cgcgggtcgt cggtgagcca gagtttcagc aggccgccca ggcggcccag gtcgccattg
2221 atgcgggcca gctcgcggac gtgctcatag tccacgacgc ccgtgatttt gtagccctgg
2281 ccgacggcca gcaggtaggc cgacaggctc atgccggccg ccgccgcctt ttcctcaatc
2341 gctcttcgtt cgtctggaag gcagtacacc ttgataggtg ggctgccctt cctggttggc
2401 ttggtttcat cagccatccg cttgccctca tctgttacgc cggcggtagc cggccagcct
2461 cgcagagcag gattcccgtt gagcaccgcc aggtgcgaat aagggacagt gaagaaggaa
2521 cacccgctcg cgggtgggcc tacttcacct atcctgcccg gctgacgccg ttggatacac
2581 caaggaaagt ctacacgaac cctttggcaa aatcctgtat atcgtgcgaa aaaggatgga
2641 tataccgaaa aaatcgctat aatgaccccg aagcagggtt atgcagcgga aaagcgctgc
2701 ttccctgctg ttttgtggaa tatctaccga ctggaaacag gcaaatgcag gaaattactg
2761 aactgagggg acaggcgaga gacgatgcca aagagctaca ccgacgagct ggccgagtgg
2821 gttgaatccc gcgcggccaa gaagcgccgg cgtgatgagg ctgcggttgc gttcctggcg
2881 gtgagggcgg atgtcgatat gcgtaaggag aaaataccgc atcaggcgca tatttgaatg
2941 tatttagaaa aataaacaaa aagagtttgt agaaacgcaa aaaggccatc cgtcaggatg
3001 gccttctgct taatttgatg cctggcagtt tatggcgggc gtcctgcccg ccaccctccg
3061 ggccgttgct tcgcaacgtt caaatccgct cccggcggat ttgtcctact caggagagcg
3121 ttcaccgaca aacaacagat aaaacgaaag gcccagtctt tcgactgagc ctttcgtttt
3181 atttgatgcc tggcagttcc ctactctcgc atggggagac cccacactac catcggcgct
3241 acggcgtttc acttctgagt tcggcatggg gtcaggtggg accaccgcgc tactgccgcc
3301 aggcaaattc tgttttatca gaccgcttct gcgttctgat ctgcccattc aaggagagtc
3361 ccatggctga ccccacggct gtgaatgccc agatcaccga tgccgtcacc cagaccaacg
3421 tcaaggtggt cgccgaagcc ccggcccagg ccatcgccgc gctctaccag atcgccagcc
3481 attccgccgg cctgtccctg cagaacgcag tctacagcca gcaggccctg aaccagatct
3541 ccaccgcggt ggtttccaag gcggtgtcga tgatcatggc gatagaagac aagaaaggta
3601 tctgacccgc cccttgatcc caggagtgcc cagacatgtt cggcttcggc aagaaatccg
3661 cgcccgcgcc cgcgccatcc ggcgacagcg agcaagccgc cgcccccggc gaggccggcc
3721 ggcagacgcc agccagcggc ggcagggcgt ccgccaacgg cgacgtgaag ctgccgtcgc
3781 gcccttcgcc gctggccggc atgagcgagc gcctgctcaa gcagaccgag gcgccgccgc
3841 cggacaccag cgccctgaac agtcaagtcg tgcaggcggt gcagatgagc aacagcgaaa
3901 ccgccggcta tgcgccgtcg cagatcgcca tcgcgccgaa catgatgatc agccaggcct
3961 ccgggctggt ggcgcaatcc gccgccagct acttcgacgg agtcagcaag atcgccctgg
4021 ccagccaggc gatcctgctc cggcagatga cccagaacat cgccgaaaac aagctggccc
4081 aggctgccga ggacgcgctc ggcgccctcg ccaccgacct gctgatgggc gccgccgccg
4141 cggtagccgc ggcggccggg gcgatcgagg cggaatcggc cagcttcgcc atcgacaaga
4201 tcgaccagag catcgccaag tacagcgaga ccctcgccag tcgcggcggc aaggacaccc
4261 aatcgtagcc accaccggcg gggccggcgg cgtccggccc tggcacagga gagccgtcat
4321 ggcctatttc aagctggaag aaccggtcag gttccatcgc tatcccttcg acttccatag
4381 tcacttcgcc ggcatcctgc cggtggaaag caacagccgc tggacgcgcg accgccgggt
4441 cttccgcgtc ggcgagcggc aagtgagcct ggagaaaggc caggagcttt ccctgatcgg
4501 gctgctgatg agtgcccgcg gcgtcgcccc ggacgtcgac ggcaaggcgc tggaggaggc
4561 tcgccaggcc gcgcactatg agctgttcga actggctctg cagcggatgg tcaggcgcaa
4621 tccgttcgcc gcgaccgacc gtcagggcta cctgcgcggc gaatgcgcgg cggagaacat
4681 ctatctcgcc tgcctcatcc tggcgcaacg ttttggccgg acctcgccgc cagcggccat
4741 cgaccagcca gcgatctacc tcggcaccct ggaactgctc ggtgcgagcg cggttcgcga
4801 cagcgagacg gaccagttcg tccgctactt caaccgcaag atctggagcg gcaacaagta
4861 cacgccgttc gacgacgcct actgggcccg cggcgccatc cgcgatcgcc atccgggcga
4921 attcgcctgc ctgacgctgg ggttcctcct ccacgaagga atcagccata cccagacggc
4981 gacgggggaa gacgaggtgg ccgtcctcga ttccctgttc gaacaattca acgcctccga
5041 gaagaccgcc taccgcctgc tcgcccacac cgcgcatggc tatgccagcg aagccgcctt
5101 cgatgccgag ctgcacagga tcctccgtca tttcgaaata cagcaggggc agccaccgca
5161 ggcccggctg gtcggcatcg acctgctcgg catggagacc gctaccggcc tgtaccgaca
5221 gttcttcgac ttcctcctcg gacaggccgc ggtgtttcgc cactacctgg acggcaagcc
5281 ggaaacccgc aagatggtcc tgcacatcca ttgtggagag gggaccgggg tgtccgacga
5341 caaccggtcg ctgtgcggtt acttcctgcg caacgccaac gcgctggacg atttctacgc
5401 agcgctctcc gcctatgcct ggaagtgcta tggcaacacc atccgccagg gcaaggcgcg
5461 gctacgcgaa cgggagaacc tgcaggaacg tgacaaggcg cccagcgccc tcgccggcct
5521 tttcgacgaa ctgtttttcg gcaacagcct gaccgcctcc ggcctgcggc tacgccgctt
5581 cgacatcacc agcgggacta cccaggccct ggtcgcctat tacgcccgca ccaacgtcgt
5641 gaacctctgc caggcgctgg ccagccgcga cgccgacggc aacagctact accgccgcct
5701 gctggagagc gatctgttca gcctgcgcat cggccatgcc tactactacc gcaactatct

5761 ggccagcaag ttccccgagc tgtgcttcga caccaacctg ggaagcaatt tcattaccgg
5821 cgcctccggc ctcttcgact cgctgcagga ataccgcctg aaccgtggcc tgcgccatct
5881 cgatggatac gtcggcaccg accagctcaa ggaactcagt ctggccatcg cctaccaggg
5941 cgaacagcgc ctggacccac aacagatgca gtacgtccat gccctggcgg aaagccagag
6001 cggcttcgac gaactgggcg aacacctccc gggtacaccc ggctgggcga aacccgcgct
6061 ggaacagttc ttcgtaagtc agtgcgcgct ctaccgctcc gaggaggatc gctacttcca
6121 gttcgaagcc taccggcgcc tcttcgccca ggtcctcaac tggcgctcct atctgctggg
6181 cgccgacggc cagggtgtcg agcactccaa cgtacaggac gaagcgatcc gcatggcgct
6241 cctgctcaac tatgccgccg cggaccgcca cggccgggtt ccggccgctt cgctggagaa
6301 tgcccaacgc ctgctggtgc aactgggcag cgcctactgg gaggaaacca tcggcgccgt
6361 cgacctggcc ggcgcgcccc atcgggaccg agagctgcag cgcttcgaag gattcgccgc
6421 ccccgcctcg gtcgtgcgca tctccacccg cagcagttga ggagcgaccg atgagcgcct
6481 acgcacgcta catcagacgc cgcgcgctcg ccgccgcgga ccgcccggcg cagcgttcgc
6541 cggacgccag cgaggaggca cgggaccaac cgccccccag caggccggcg agcggaaagc
6601 gcctgctgga aatatcggac agcgatgcgg ccaagctctt ggccgtgctg aaactgctcg
6661 gatgagcacc caccccagga gaagcgtcat gaaggaagaa ctcgaaaccc tcgtcaccca
6721 ggtcaaggca ctcctcaccg aatgcctgca aggctggctc caggccctgt cccagggaga
6781 cagcgtggac gacccggcca gcctcggaca gttggcggtg cagttgagca ccgcctacag
6841 caccctgcag gccgctgccg ccgcctaccc cggcgatacc ggcagcagcg cgcaggcccc
6901 ctcggcggac ggcacgcagc agtactgaag cctcgccacg acaccccgcg caccgcgcgt
6961 ccatcgcaga ggagtccgcg catgtcgatc tcggaagaat tgggaaggaa tctggtccag
7021 ctggcgcaac gctggctgaa acgcccccctc accagcgaag aacacgagca gttgagacag
7081 ttccagctcg gcctcggcaa cgacgccagc agcgatccgg ccgtgcacgc gcgacagcag
7141 gccgaacagg tgatcgagaa cggccggcag cgttccgaag gctccatccg ccagatcctg
7201 cagaacgtcc agcaggcctc gacccaggcc ctccgcgccc aggacagcgg cgagcaggcg
7261 attctcgagg cggtcaatgc cgccggcagc ctggccgaac tgcgtcccgg ccacctgcag
7321 agcggccagg caagcgccaa ccagttggtg atgtcgcaga ttgccgaccg cctggccaac
7381 ctagtgaaga gcgaggtacg cctgtgcttc gagcagaact tcggctcgct caagcagcaa
7441 ctggagaacg ctatcgcagc tgtttcctgt gtgaaattgt tatccgctca caattccaca
7501 caacatacga gccggaagca taaagtgtaa agcctggggt gcctaatgag tgagctaact
7561 cacattaatt gcgttgcgct cactgcccgc tttccagtcg ggaaacctgt cgtgccagct
7621 gcattaatga atcggccaac gcgcggggag aggcggtttg cgtattgggc gctcttccgc
7681 ttcctcgctc actgactcgc tgcgctcggt cgttcggctg cggcgagcgg tatcagctca
7741 ctcaaaggcg gtaatacggt tatccacaga atcaggggat aacgcaggaa agaacatgtg
7801 agcaaaaggc cagcaaaagg ccaggaaccg taaaaaggcc gcgttgctgg cgtttttcca
7861 taggctccgc ccccctgacg agcatcacaa aaatcgacgc tcaagtcaga ggtggcgaaa
7921 cccgacagga ctataaagat accaggcgtt tccccctgga agctccctcg tgcgctctcc
7981 tgttccgacc ctgccgctta ccggatacct gtccgccttt ctcccttcgg gaagcgtggc
8041 gctttctcat agctcacgct gtaggtatct cagttcggtg taggtcgttc gctccaagct
8101 gggctgtgtg cacgaacccc ccgttcagcc cgaccgctgc gccttatccg gtaactatcg
8161 tcttgagtcc aacccggtaa gacacgactt atcgccactg gcagcagcca ctggtaacag
8221 gattagcaga gcgaggtatg taggcggtgc tacagagttc ttgaagtggt ggcctaacta
8281 cggctacact agaaggacag tatttggtat ctgcgctctg ctgaagccag ttaccttcgg
8341 aaaaagagtt ggtagctctt gatccggcaa acaaaccacc gctggtagcg gtggtttttt
8401 tgtttgcaag cagcagatta cgcgcagaaa aaaaggatct caagaagatc ctttgatctt
8461 ttctacgggg tctgacgctc agtggaacga aaactcacgt taagggattt tggtcatgag
8521 attatcaaaa aggatcttca cctagatcct tttaaattaa aaatgaagtt ttaaatcaat
8581 ctaaagtata tatgagtaaa cttggtctga caatcgatgc gaattggccg cggcgttgtg
8641 acaatttacc gaacaactcc gcggccggga agccgatctc ggcttgaacg aattgttagg
8701 tggcggtact tgggtcgata tcaaagtgca tcacttcttc ccgtatgccc aactttgtat
8761 agagagccac tgcgggatcg tcaccgtaat ctgcttgcac gtagatcaca taagcaccaa
8821 gcgcgttggc ctcatgcttg aggagattga tgagcgcggt ggcaatgccc tgcctccggt
8881 gctcgccgga gactgcgaga tcatagatat agatctcact acgcggctgc tcaaacctgg
8941 gcagaacgta agccgcgaga gcgccaacaa ccgcttcttg gtcgaaggca gcaagcgcga
9001 tgaatgtctt actacggagc aagttcccga ggtaatcgga gtccggctga tgttgggagt
9061 aggtggctac gtctccgaac tcacgaccga aaagatcaag agcagcccgc atggatttga
9121 cttggtcagg gccgagccta catgtgcgaa tgatgcccat acttgagcca cctaactttg
9181 ttttagggcg actgccctgc tgcgtaacat cgttgctgct gcgtaacatc gttgctgctc
9241 cataacatca aacatcgacc cacggcgtaa cgcgcttgct gcttggatgc ccgaggcata
9301 gactgtacaa aaaaacagtc ataacaagcc atgaaaaccg ccactgcgcc gttaccaccg
9361 ctgcgttcgg tcaaggttct ggaccagttg cgtgagcgca tacgctactt gcattacagt
9421 ttacgaaccg aacaggctta tgtcaattcg gttgaatact catactcttc ctttttcaat
9481 attattgaag catttatcag ggttattgtc tcatgagcgg atacatattt gaatgtattt
9541 agaaaaataa acaaataggg gttccgcgca catttccccg aaaagtgggt aataactgat
9601 ataattaaat tgaagctcta atttgtgagt ttagtataca tgcatttact tataatacag
9661 tttttagtt ttgctggccg catcttctca aatatgcttc ccagcctgct tttctgtaac
9721 gttcaccctc taccttagca tcccttccct ttgcaaatag tcctcttcca acaataataa

```
 9781 tgtcagatcc tgtagagacc acatcatcca cggttctata ctgttgaccc aatgcgtctc
 9841 ccttgtcatc taaacccaca ccgggtgtca taatcaacca atcgtaacct tcatctcttc
 9901 cacccatgtc tctttgagca ataaagccga taacaaaatc tttgtcgctc ttcgcaatgt
 9961 caacagtacc cttagtatat tctccagtag atagggagcc cttgcatgac aattctgcta
10021 acatcaaaag gcctctaggt tcctttgtta cttcttctgc cgcctgcttc aaaccgctaa
10081 caatacctgg gcccaccaca ccgtgtgcat tcgtaatgtc tgcccattct gctattctgt
10141 atacacccgc agagtactgc aatttgactg tattaccaat gtcagcaaat tttctgtctt
10201 cgaagagtaa aaaattgtac ttggcggata atgcctttag cggcttaact gtgccctcca
10261 tggaaaaatc agtcaagata tccacatgtg tttttagtaa acaaattttg ggacctaatg
10321 cttcaactaa ctccagtaat tccttggtgg tacgaacatc caatgaagca cacaagtttg
10381 tttgcttttc gtgcatgata ttaaatagct tggcagcaac aggactagga tgagtagcag
10441 cacgttcctt atatgtagct ttcgacatga tttatcttcg tttcctgcag gtttttgttc
10501 tgtgcagttg ggttaagaat actgggcaat ttcatgtttc ttcaacacta catatgcgta
10561 tatataccaa tctaagtctg tgctccttcc ttcgttcttc cttctgttcg gagattaccg
10621 aatcaaaaaa atttcaaaga aaccgaaatc aaaaaaaaga ataaaaaaaa aatgatgaat
10681 tgaattgaaa agctagctta tcgatgggtc cttttcatca cgtgctataa aaataattat
10741 aatttaaatt ttttaatata aatatataaa ttaaaaatag aaagtaaaaa aagaaattaa
10801 agaaaaaata gtttttgttt tccgaagatg taaaagactc tagggggatc gccaacaaat
10861 actacctttt atcttgctct tcctgctctc aggtattaat gccgaattgt ttcatcttgt
10921 ctgtgtagaa gaccacacac gaaaatcctg tgattttaca ttttacttat cgttaatcga
10981 atgtatatct atttaatctg cttttcttgt ctaataaata tatatgtaaa gtacgctttt
11041 tgttgaaatt ttttaaacct ttgtttattt ttttttcttc attccgtaac tcttctacct
11101 tctttattta ctttctaaaa tccaaataca aaacataaaa ataaataaac acagagtaaa
11161 ttcccaaatt attccatcat taaaagatac gaggcgcgtg taagttacag gcaagcgatc
11221 tctaagaaac cattattatc atgacattaa cctataaaaa taggcgtatc acgaggccct
11281 ttcgtc
```

#### 11. pMQ30-27675-mScarlet

**Description** Allelic-exchange construct creating a C-terminal mScarlet-I translational fusion to PA14_27675 at the native locus. pMQ30 allelic-exchange (suicide) vector: Gm(R) aacC1, sacB counter-selection, oriT, URA3 + CEN6/ARSH4 for yeast gap-repair assembly (Shanks et al. 2006)

**Length** 9,880 bp, circular

**Source** This paper

**Note** *PA14_27675 3' homology arm (3342..4337, C-terminal 332 aa), GSHASG linker (4338..4355) and mScarlet-I (4356..5051, 231 aa) form one continuous 569-codon reading frame (3342..5051) with no internal stop codons.*

##### Features

| Location | Type | Feature |
|---|---|---|
| c(209..1630) | gene | sacB |
| 2104..2928 | misc_feature | oriT (conjugal transfer origin, from pEX18-Gm) |
| 3341..4337 | misc_feature | PA14_27675 3' homology arm |
| 3342..4337 | CDS | PA14_27675 (dear) 3' portion, C-terminal 332 aa (partial) |
| 4338..4355 | misc_feature | linker (GSHASG) |
| 4356..5051 | CDS | mScarlet-I (231 aa) |
| 5062..5256 | misc_feature | PA14_27670 |
| 5280..5519 | misc_feature | PA14_27660 |
| c(6069..6091) | misc_binding | lac operator |
| c(6095..6125) | promoter | lac promoter |
| 6426..7026 | rep_origin | ori (ColE1) |
| c(7290..7823) | gene | aacC1 (gentamicin resistance) |
| c(8181..9288) | gene | URA3 |
| 9301..9814 | misc_feature | CEN6/ARSH4; *S. cerevisiae* replicon from pYC2-CT |

##### Sequence (9,880 bp)

```
   1 tcgcgcgttt cggtgatgac ggtgaaaacc tctgacacat gcagctcccg gagacggtca
  61 cagcttgtct gtaagcggat gccgggagca gacaagcccg tcagggcgcg tcagcgggtg
 121 ttggcgggtg tcggggctgg cttaactatg cggcatcaga gcagattgta ctgagagtgc
 181 accataatcg gcattttctt ttgcgttttt atttgttaac tgttaattgt ccttgttcaa
 241 ggatgctgtc tttgacaaca gatgttttct tgcctttgat gttcagcagg aagctaggcg
 301 caaacgttga ttgtttgtct gcgtagaatc ctctgtttgt catatagctt gtaatcacga
 361 cattgtttcc tttcgcttga ggtacagcga agtgtgagta agtaaaggtt acatcgttag
 421 gatcaagatc catttttaac acaaggccag ttttgttcag cggcttgtat gggccagtta
 481 aagaattaga aacataacca agcatgtaaa tatcgttaga cgtaatgccg tcaatcgtca
 541 tttttgatcc gcgggagtca gtgaacagat accatttgcc gttcatttta aagacgttcg
 601 cgcgttcaat ttcatctgtt actgtgttag atgcaatcag cggtttcatc acttttttca
 661 gtgtgtaatc atcgtttagc tcaatcatac cgagagcgcc gtttgctaac tcagccgtgc
 721 gtttttttatc gctttgcaga agtttttgac tttcttgacg gaagaatgat gtgcttttgc
 781 catagtatgc tttgttaaat aaagattctt cgccttggta gccatcttca gttccagtgt
 841 ttgcttcaaa tactaagtat ttgtggcctt tatcttctac gtagtgagga tctctcagcg
 901 tatggttgtc gcctgagctg tagttgcctt catcgatgaa ctgctgtaca ttttgatacg
 961 tttttccgtc accgtcaaag attgatttat aatcctctac accgttgatg ttcaaagagc
1021 tgtctgatgc tgatacgtta acttgtgcag ttgtcagtgt ttgtttgccg taatgtttac
1081 cggagaaatc agtgtagaat aaacggattt ttccgtcaga tgtaaatgtg gctgaacctg
1141 accattcttg tgtttggtct tttaggatag aatcatttgc atcgaatttg tcgctgtctt
1201 taaagacgcg gccagcgttt ttccagctgt caatagaagt ttcgccgact ttttgataga
```

1261 acatgtaaat cgatgtgtca tccgcatttt taggatctcc ggctaatgca aagacgatgt
1321 ggtagccgtg atagtttgcg acagtgccgt cagcgttttg taatggccag ctgtcccaaa
1381 cgtccaggcc ttttgcagaa gagatatttt taattgtgga cgaatcgaac tcaggaactt
1441 gatatttttc atttttttgc tgttcaggga tttgcagcat atcatggcgt gtaatatggg
1501 aaatgccgta tgtttcctta tatggctttt ggttcgtttc tttcgcaaac gcttgagttg
1561 cgcctcctgc cagcagtgcg gtagtaaagg ttaatactgt tgcttgtttt gcaaactttt
1621 tgatgttcat cgttcatgtc tcctttttta tgtactgtgt tagcggtctg cttcttccag
1681 ccctcctgtt tgaagatggc aagttagtta cgcacaataa aaaaagacct aaaatatgta
1741 aggggtgacg ccaaagtata cactttgccc tttacacatt ttaggtcttg cctgctttat
1801 cagtaacaaa cccgcgcgat ttacttttcg acctcattct attagactct cgtttggatt
1861 gcaactggtc tattttcctc ttttgtttga tagaaaatca taaaaggatt tgcagactac
1921 gggcctaaag aactaaaaaa tctatctgtt tcttttcatt ctctgtattt tttatagttt
1981 ctgttgcatg ggcataaagt tgcctttta atcacaattc agaaaatatc ataatatctc
2041 atttcactaa ataatagtga acggcaggta tatgtgatgg gttaaaaagg atcgatcctc
2101 tagctagagt cgatcttcgc cagcagggcg aggatcgtgg catcaccgaa ccgcgccgtg
2161 cgcgggtcgt cggtgagcca gagtttcagc aggccgccca ggcggcccag gtcgccattg
2221 atgcgggcca gctcgcggac gtgctcatag tccacgacgc ccgtgatttt gtagccctgg
2281 ccgacggcca gcaggtaggc cgacaggctc atgccggccg ccgccgcctt ttcctcaatc
2341 gctcttcgtt cgtctggaag gcagtacacc ttgataggtg ggctgccctt cctggttggc
2401 ttggtttcat cagccatccg cttgccctca tctgttacgc cggcggtagc cggccagcct
2461 cgcagagcag gattcccgtt gagcaccgcc aggtgcgaat aagggacagt gaagaaggaa
2521 cacccgctcg cgggtgggcc tacttcacct atcctgcccg gctgacgccg ttggatacac
2581 caaggaaagt ctacacgaac cctttggcaa aatcctgtat atcgtgcgaa aaaggatgga
2641 tataccgaaa aaatcgctat aatgaccccg aagcagggtt atgcagcgga aaagcgctgc
2701 ttccctgctg ttttgtggaa tatctaccga ctggaaacag gcaaatgcag gaaattactg
2761 aactgagggg acaggcgaga gacgatgcca aagagctaca ccgacgagct ggccgagtgg
2821 gttgaatccc gcgcggccaa gaagcgccgg cgtgatgagg ctgcggttgc gttcctggcg
2881 gtgagggcgg atgtcgatat gcgtaaggag aaaataccgc atcaggcgca tatttgaatg
2941 tatttagaaa aataaacaaa aagagtttgt agaaacgcaa aaaggccatc cgtcaggatg
3001 gccttctgct taatttgatg cctggcagtt tatggcgggc gtcctgcccg ccaccctccg
3061 ggccgttgct tcgcaacgtt caaatccgct cccggcggat ttgtcctact caggagagcg
3121 ttcaccgaca aacaacagat aaaacgaaag gcccagtctt tcgactgagc ctttcgtttt
3181 atttgatgcc tggcagttcc ctactctcgc atggggagac cccacactac catcggcgct
3241 acggcgtttc acttctgagt tcggcatggg gtcaggtggg accaccgcgc tactgccgcc
3301 aggcaaattc tgttttatca gaccgcttct gcgttctgat gctacgcgaa cgggagaacc
3361 tgcaggaacg tgacaaggcg cccagcgccc tcgccggcct tttcgacgaa ctgtttttcg
3421 gcaacagcct gaccgcctcc ggcctgcggc tacgccgctt cgacatcacc agcgggacta
3481 cccaggccct ggtcgcctat tacgcccgca ccaacgtcgt gaacctctgc caggcgctgg
3541 ccagccgcga cgccgacggc aacagctact accgccgcct gctggagagc gatctgttca
3601 gcctgcgcat cggccatgcc tactactacc gcaactatct ggccagcaag ttccccgagc
3661 tgtgcttcga caccaacctg ggaagcaatt tcattaccgg cgcctccggc ctcttcgact
3721 cgctgcagga ataccgcctg aaccgtggcc tgcgccatct cgatggatac gtcggcaccg
3781 accagctcaa ggaactcagt ctggccatcg cctaccaggg cgaacagcgc ctggacccac
3841 aacagatgca gtacgtccat gccctggcgg aaagccagag cggcttcgac gaactgggcg
3901 aacacctccc gggtacaccc ggctgggcga aacccgcgct ggaacagttc ttcgtaagtc
3961 agtgcgcgct ctaccgctcc gaggaggatc gctacttcca gttcgaagcc taccggcgcc
4021 tcttcgccca ggtcctcaac tggcgctcct atctgctggg cgccgacggc cagggtgtcg
4081 agcactccaa cgtacaggac gaagcgatcc gcatggcgct cctgctcaac tatgccgccg
4141 cggaccgcca cggccgggtt ccggccgctt cgctggagaa tgcccaacgc ctgctggtgc
4201 aactgggcag cgcctactgg gaggaaacca tcggcgccgt cgacctggcc ggcgcgcccc
4261 atcgggaccg agagctgcag cgcttcgaag gattcgccgc ccccgcctcg gtcgtgcgca
4321 tctccacccg cagcagtggc tcgcatgcgt ccggcatgag taaaggagaa gctgtgatta
4381 aagagttcat gcgcttcaaa gttcacatgg agggttctat gaacggtcac gagttcgaga
4441 tcgaaggcga aggcgagggc cgtccgtatg aaggcaccca gaccgccaaa ctgaaagtga
4501 ctaaaggcgg cccgctgcct ttttcctggg acatcctgag cccgcaattt atgtacggtt
4561 ctagggcgtt catcaaacac ccagcggata tcccggacta ttataagcag tcttttccgg
4621 aaggtttcaa gtgggaacgc gtaatgaatt ttgaagatgg tggtgccgtg accgtcactc
4681 aggacacctc cctggaggat ggcaccctga tctataaagt taaactgcgt ggtactaatt
4741 ttccacctga tggcccggtg atgcagaaaa agacgatggg ttgggaggcg tctaccgaac
4801 gcttgtatcc ggaagatggt gtgctgaaag gcgacattaa aatggccctg cgcctgaaag
4861 atggcggccg ctatctggct gacttcaaaa ccacgtacaa agccaagaaa cctgtgcaga
4921 tgcctggcgc gtacaatgtg gaccgcaaac tggacatcac ctctcataat gaagattata
4981 cggtggtaga gcaatatgag cgctccgagg gtcgtcattc taccggtggc atggatgaac
5041 tatacaaata aggagcgacc gatgagcgcc tacgcacgct acatcagacg ccgcgcgctc
5101 gccgccgcgg accgcccggc gcagcgttcg ccggacgcca gcgaggaggc acgggaccaa
5161 ccgcccccca gcaggccggc gagcggaaag cgcctgctgg aaatatcgga cagcgatgcg
5221 gccaagctct tggccgtgct gaaactgctc ggatgagcac ccaccccagg agaagcgtca

5281 tgaaggaaga actcgaaacc ctcgtcaccc aggtcaaggc actcctcacc gaatgcctgc
5341 aaggctggct ccaggccctg tcccagggag acagcgtgga cgacccggcc agcctcggac
5401 agttggcggt gcagttgagc accgcctaca gcaccctgca ggccgctgcc gccgcctacc
5461 ccggcgatac cggcagcagc gcgcaggccc cctcggcgga cggcacgcag cagtactgaa
5521 gcctcgccac gacaccccgc gcaccgcgcg tccatcgcag aggagtccgc gcatgtcgat
5581 ctcggaagaa ttgggaagga atctggtcca gctggcgcaa cgctggctga aacgcccccct
5641 caccagcgaa gaacacgagc agttgagaca gttccagctc ggcctcggca acgacgccag
5701 cagcgatccg gccgtgcacg cgcgacagca ggccgaacag gtgatcgaga acggccggca
5761 gcgttccgaa ggctccatcc gccagatcct gcagaacgtc cagcaggcct cgacccaggc
5821 cctccgcgcc caggacagcg gcgagcaggc gattctcgag gcggtcaatg ccgccggcag
5881 cctggccgaa ctgcgtcccg gccacctgca gagcggccag gcaagcgcca accagttggt
5941 gatgtcgcag attgccgacc gcctggccaa cctagtgaag agcgaggtac gcctgtgctt
6001 cgagcagaac ttcggctcgc tcaagcagca actggagaac gctatcgcca tagctgtttc
6061 ctgtgtgaaa ttgttatccg ctcacaattc cacacaacat acgagccgga agcataaagt
6121 gtaaagcctg gggtgcctaa tgagtgagct aactcacatt aattgcgttg cgctcactgc
6181 ccgctttcca gtcgggaaac ctgtcgtgcc agctgcatta atgaatcggc caacgcgcgg
6241 ggagaggcgg tttgcgtatt gggcgctctt ccgcttcctc gctcactgac tcgctgcgct
6301 cggtcgttcg gctgcggcga gcggtatcag ctcactcaaa ggcggtaata cggttatcca
6361 cagaatcagg ggataacgca ggaaagaaca tgtgagcaaa aggccagcaa aaggccagga
6421 accgtaaaaa ggccgcgttg ctggcgtttt tccataggct ccgcccccct gacgagcatc
6481 acaaaaatcg acgctcaagt cagaggtggc gaaacccgac aggactataa agataccagg
6541 cgtttccccc tggaagctcc ctcgtgcgct ctcctgttcc gaccctgccg cttaccggat
6601 acctgtccgc ctttctccct tcgggaagcg tggcgctttc tcatagctca cgctgtaggt
6661 atctcagttc ggtgtaggtc gttcgctcca agctgggctg tgtgcacgaa ccccccgttc
6721 agcccgaccg ctgcgcctta tccggtaact atcgtcttga gtccaacccg gtaagacacg
6781 acttatcgcc actggcagca gccactggta acaggattag cagagcgagg tatgtaggcg
6841 gtgctacaga gttcttgaag tggtggccta actacggcta cactagaagg acagtatttg
6901 gtatctgcgc tctgctgaag ccagttacct tcggaaaaag agttggtagc tcttgatccg
6961 gcaaacaaac caccgctggt agcggtggtt tttttgtttg caagcagcag attacgcgca
7021 gaaaaaaagg atctcaagaa gatcctttga tcttttctac ggggtctgac gctcagtgga
7081 acgaaaactc acgttaaggg attttggtca tgagattatc aaaaaggatc ttcacctaga
7141 tccttttaaa ttaaaaatga agttttaaat caatctaaag tatatatgag taaacttggt
7201 ctgacaatcg atgcgaattg gccgcggcgt tgtgacaatt taccgaacaa ctccgcggcc
7261 gggaagccga tctcggcttg aacgaattgt taggtggcgg tacttgggtc gatatcaaag
7321 tgcatcactt cttcccgtat gcccaacttt gtatagagag ccactgcggg atcgtcaccg
7381 taatctgctt gcacgtagat cacataagca ccaagcgcgt tggcctcatg cttgaggaga
7441 ttgatgagcg cggtggcaat gccctgcctc cggtgctcgc cggagactgc gagatcatag
7501 atatagatct cactacgcgg ctgctcaaac ctgggcagaa cgtaagccgc gagagcgcca
7561 acaaccgctt cttggtcgaa ggcagcaagc gcgatgaatg tcttactacg gagcaagttc
7621 ccgaggtaat cggagtccgg ctgatgttgg gagtaggtgg ctacgtctcc gaactcacga
7681 ccgaaaagat caagagcagc ccgcatggat ttgacttggt cagggccgag cctacatgtg
7741 cgaatgatgc ccatacttga gccacctaac tttgttttag ggcgactgcc ctgctgcgta
7801 acatcgttgc tgctgcgtaa catcgttgct gctccataac atcaaacatc gacccacggc
7861 gtaacgcgct tgctgcttgg atgcccgagg catagactgt acaaaaaaac agtcataaca
7921 agccatgaaa accgccactg cgccgttacc accgctgcgt tcggtcaagg ttctggacca
7981 gttgcgtgag cgcatacgct acttgcatta cagtttacga accgaacagg cttatgtcaa
8041 ttcggttgaa tactcatact cttccttttt caatattatt gaagcattta tcagggttat
8101 tgtctcatga gcggatacat atttgaatgt atttagaaaa ataaacaaat aggggttccg
8161 cgcacatttc cccgaaaagt gggtaataac tgatataatt aaattgaagc tctaatttgt
8221 gagtttagta tacatgcatt tacttataat acagtttttt agttttgctg gccgcatctt
8281 ctcaaatatg cttcccagcc tgcttttctg taacgttcac cctctacctt agcatccctt
8341 ccctttgcaa atagtcctct tccaacaata ataatgtcag atcctgtaga gaccacatca
8401 tccacggttc tatactgttg acccaatgcg tctcccttgt catctaaacc cacaccgggt
8461 gtcataatca accaatcgta accttcatct cttccaccca tgtctctttg agcaataaag
8521 ccgataacaa aatctttgtc gctcttcgca atgtcaacag taccccttagt atattctcca
8581 gtagataggg agcccttgca tgacaattct gctaacatca aaaggcctct aggttccttt
8641 gttacttctt ctgccgcctg cttcaaaccg ctaacaatac ctgggcccac cacaccgtgt
8701 gcattcgtaa tgtctgccca ttctgctatt ctgtatacac ccgcagagta ctgcaatttg
8761 actgtattac caatgtcagc aaattttctg tcttcgaaga gtaaaaaatt gtacttggcg
8821 gataatgcct ttagcggctt aactgtgccc tccatggaaa aatcagtcaa gatatccaca
8881 tgtgttttta gtaaacaaat tttgggacct aatgcttcaa ctaactccag taattccttg
8941 gtggtacgaa catccaatga agcacacaag tttgtttgct tttcgtgcat gatattaaat
9001 agcttggcag caacaggact aggatgagta gcagcacgtt ccttatatgt agctttcgac
9061 atgatttatc ttcgtttcct gcaggttttt gttctgtgca gttgggttaa gaatactggg
9121 caatttcatg tttcttcaac actacatatg cgtatatata ccaatctaag tctgtgctcc
9181 ttccttcgtt cttccttctg ttcggagatt accgaatcaa aaaaatttca aagaaaccga
9241 aatcaaaaaa aagaataaaa aaaaaatgat gaattgaatt gaaaagctag cttatcgatg

9301 ggtccttttc atcacgtgct ataaaaataa ttataattta aatttttttaa tataaatata
9361 taaattaaaa atagaaagta aaaaaagaaa ttaaagaaaa aatagttttt gttttccgaa
9421 gatgtaaaag actctagggg gatcgccaac aaatactacc ttttatcttg ctcttcctgc
9481 tctcaggtat taatgccgaa ttgtttcatc ttgtctgtgt agaagaccac acacgaaaat
9541 cctgtgattt tacattttac ttatcgttaa tcgaatgtat atctatttaa tctgcttttc
9601 ttgtctaata aatatatatg taaagtacgc tttttgttga aatttttttaa acctttgttt
9661 attttttttt cttcattccg taactcttct accttcttta tttactttct aaaatccaaa
9721 tacaaaacat aaaaataaat aaacacagag taaattccca aattattcca tcattaaaag
9781 atacgaggcg cgtgtaagtt acaggcaagc gatctctaag aaaccattat tatcatgaca
9841 ttaacctata aaaataggcg tatcacgagg ccctttcgtc

#### 12. pAKN69-rhaSR-PrhaBAD-rcgA

**Description** Rhamnose-inducible single-copy expression construct for rcgA (PA14_27700), Tn7-based chromosomal integration vector (Gm(R), Cm(R), oriT/traJ, RhaS/RhaR, PrhaBAD).

**Length** 9,725 bp, circular

**Source** This paper

##### Features

| Location | Type | Feature |
|---|---|---|
| 1..589 | misc_feature | ori |
| c(1428..1799) | misc_feature | traJ |
| 1832..1941 | misc_feature | oriT |
| c(2895..2981) | misc_feature | rrnB T1 terminator |
| c(3046..3705) | misc_feature | cat (chloramphenicol resistance) |
| c(3745..3843) | misc_feature | terminator |
| c(3847..4572) | CDS | rcgA (PA14_27700), 241 aa |
| c(4596..4714) | misc_feature | PrhaBAD promoter |
| 4860..5696 | misc_feature | rhaS |
| c(5770..6618) | misc_feature | rhaR |
| 7016..7549 | misc_feature | aacC1 (gentamicin resistance) |
| c(7738..7766) | misc_feature | Pc promoter |
| 7890..8055 | misc_feature | Tn7L |
| 8590..8694 | misc_feature | bla promoter |
| 8695..9555 | misc_feature | bla (ampicillin resistance) |

##### Sequence (9,725 bp)

```
   1 ttgagatcct tttttctgc gcgtaatctg ctgcttgcaa acaaaaaaac caccgctacc
  61 agcggtggtt tgtttgccgg atcaagagct accaactctt tttccgaagg taactggctt
 121 cagcagagcg cagataccaa atactgttct tctagtgtag ccgtagttag gccaccactt
 181 caagaactct gtagcaccgc ctacatacct cgctctgcta atcctgttac cagtggctgc
 241 tgccagtggc gataagtcgt gtcttaccgg gttggactca agacgatagt taccggataa
 301 ggcgcagcgg tcgggctgaa cggggggttc gtgcacacag cccagcttgg agcgaacgac
 361 ctacaccgaa ctgagatacc tacagcgtga gctatgagaa agcgccacgc ttcccgaagg
 421 gagaaaggcg gacaggtatc cggtaagcgg cagggtcgga acaggagagc gcacgaggga
 481 gcttccaggg ggaaacgcct ggtatcttta tagtcctgtc gggtttcgcc acctctgact
 541 tgagcgtcga tttttgtgat gctcgtcagg ggggcggagc ctatggaaaa acgccagcaa
 601 cgcggccttt ttacggttcc tggccttttg ctggcctttt gctcacatgt tctttcctgc
 661 gttatcccct gattctgtgg ataaccgtat taccgccttt gagtgagctg ataccgctcg
 721 ccgcagccga acgaccgagc gcagcgagtc agtgagcgag gaagcggaag agcgcccaat
 781 acgcaaaccg cctctccccg cgcgttggcc gattcattaa tgcaggactc tagaggatcc
 841 agccgaccag gctttccacg cccgcgtgcc gctccatgtc gttcgcgcgg ttctcggaaa
 901 cgcgctgccg cgtttcgtga ttgtcacgct caagcccgta gtcccgttcg agcgtcgcgc
 961 agaggtcagc gagggcgcgg taggcccgat acggctcatg gatggtgttt cgggtcgggt
1021 gaatcttgtt gatggcgata tggatgtgca ggttgtcggt gtcgtgatgc acggcactga
1081 cgcgctgatg ctcggcgaag ccaagcccag cgcagatgcg gtcctcaatc gcgcgcaacg
1141 tctccgcgtc gggcttctct cccgcgcgga agctaaccag caggtgatag gtcttgtcgg
1201 cctcggaacg ggtgttgccg tgctgggtcg ccatcacctc ggccatgaca gcgggcaggg
1261 tgtttgcctc gcagttcgtg acgcgcacgt gacccaggcg ctcggtcttg ccttgctcgt
1321 cggtgatgta cttcaccagc tccgcgaagt cgctcttctt gatggagcgc atggggacgt
1381 gcttggcaat cacgcgcacc ccccggccgt tttagcggct aaaaaagtca tggctctgcc
1441 ctcgggcgga ccacgcccat catgaccttg ccaagctcgt cctgcttctc ttcgatcttc
1501 gccagcaggg cgaggatcgt ggcatcaccg aaccgcgccg tgcgcgggtc gtcggtgagc
```

1561 cagagtttca gcaggccgcc caggcggccc aggtcgccat tgatgcgggc cagctcgcgg
1621 acgtgctcat agtccacgac gcccgtgatt ttgtagccct ggccgacggc cagcaggtag
1681 gccgacaggc tcatgccggc cgccgccgcc ttttcctcaa tcgctcttcg ttcgtctgga
1741 aggcagtaca ccttgatagg tgggctgccc ttcctggttg gcttggtttc atcagccatc
1801 cgcttgccct catctgttac gccggcggta gccggccagc ctcgcagagc aggattcccg
1861 ttgagcaccg ccaggtgcga ataagggaca gtgaagaagg aacacccgct cgcgggtggg
1921 cctacttcac ctatcctgcc cggctgacgc cgttggatac accaaggaaa gtctacacga
1981 accctttggc aaaatcctgt atatcgtgcg aaaaaggatg gatataccga aaaaatcgct
2041 ataatgaccc cgaagcaggg ttatgcagcg gaaaagcgct gcttccctgc tgttttgtgg
2101 aatatctacc gactggaaac aggcaaatgc aggaaattac tgaactgagg ggacaggcga
2161 gagacgatgc caaagagcta caccgacgag ctggccgagt gggttgaatc ccgcgcggcc
2221 aagaagcgcc ggcgtgatga ggctgcggtt gcgttcctgg cggtgagggc ggatgtcgag
2281 gcggcgttag cgtccggcta tgcgctcgtc accatttggg agcacatgcg ggaaacgggg
2341 aaggtcaagt tctcctacga gacgttccgc tcgcacgcca ggcggcacat caaggccaag
2401 cccgccgatg tgcccgcacc gcaggccaag gctgcggaac ccgcgccggc acccaagacg
2461 ccggagccac ggcggccgaa gcaggggggc aaggctgaaa agccggcccc cgctgcggcc
2521 ccgaccggct tcaccttcaa cccaacaccg gacaaaaagg atcctctaga ggaccagccg
2581 cgtaacctgg caaaatcggt tacggttgag taataaatgg atgccctgcg taagcgggtg
2641 tgggcggaca ataaagtctt aaactgaaca aaatagatct aaactatgac aataaagtct
2701 taaactagac agaatagttg taaactgaaa tcagtccagt tatgctgtga aaaagcatac
2761 tggacttttg ttatggctaa agcaaactct tcattttctg aagtgcaaat tgcccgtcgt
2821 attaaagagg ggcgtggggt cgaaattccc ggggatccac gcgtcttaag gcggccgcaa
2881 gctcctagcg gcggatttgt cctactcagg agagcgttca ccgacaaaca acagataaaa
2941 cgaaaggccc agtctttcga ctgagccttt cgttttattt gatgcctcaa gctagagagt
3001 cattacccca ggcgtttaag ggcaccaata actgccttaa aaaaattacg ccccgccctg
3061 ccactcatcg cagtactgtt gtaattcatt aagcattctg ccgacatgga agccatcaca
3121 aacggcatga tgaacctgaa tcgccagcgg catcagcacc ttgtcgcctt gcgtataata
3181 tttgcccatg gtgaaaacgg gggcgaagaa gttgtccata ttggccacgt ttaaatcaaa
3241 actggtgaaa ctcacccagg gattggctga gacgaaaaac atattctcaa taaacccttt
3301 agggaaatag gccaggtttt caccgtaaca cgccacatct tgcgaatata tgtgtagaaa
3361 ctgccggaaa tcgtcgtggt attcactcca gagcgatgaa aacgtttcag tttgctcatg
3421 gaaaacggtg taacaagggt gaacactatc ccatatcacc agctcaccgt ctttcattgc
3481 catacgaaat tccggatgag cattcatcag gcgggcaaga atgtgaataa aggccggata
3541 aaacttgtgc ttatttttct ttacggtctt taaaaaggcc gtaatatcca gctgaacggt
3601 ctggttatag gtacattgag caactgactg aaatgcctca aaatgttctt tacgatgcca
3661 ttgggatata tcaacggtgg tatatccagt gatttttttc tccattttag cttccttagc
3721 tcctgaaaat ctcgccaagc tagcttggat tctcaccaat aaaaaacgcc cggcggcaac
3781 cgagcgttct gaacaaatcc agatggagtt ctgaggtcat tactggatct atcaacagga
3841 gtccaattac ctgggtgaca ggggcgcggc tggtagttcg gccttgcgtc gcaggcgacc
3901 gaggtcgttg atcaggatcc ggtcgttgcg gcagctgatc agcccttcgc gtttccatat
3961 ttgcagttgc gcgttgagct tcggccggct ggcgttgagc atcgctgcta ggatgctctg
4021 gttggtcggt agcgtcgcgc aggctccctc gggttgttcg ctcgccaggc tggccaggag
4081 atggcgcgcc aggcgcgact ccaggcgatg caggcagacc ctctcgacga aaccggaggc
4141 ttcgcgcagg cgcgaacaga ggagggccag cgcgcggttg aggaattccg tgtcggcgat
4201 caacggggcg aagtgacgtc ggtggaggat cagcaccgac gccgcgccgg aggcgtaggc
4261 agaggtctcg cggcggttcg gctggatcag cgcggtttcc ccgatcactt cgccaggacg
4321 gatatcattg acgatcagct cccgcccgtc cggcccgtag agcacggtat agaggtgtcc
4381 gccgaggacc aggccgatga agtcttcagg gtcgtgcttg aggaacagga cgtccttgtc
4441 gcgcaaccgg cgctccgacg catgggtcgc cacgtgccgc agcagatcgg gcagcccgtg
4501 cagcagcggg caccgttgca gcagggtgta gtggcgtgaa cagcggtctt ccgttgactc
4561 gttcgagtgc atgcttaatt tctcctcttt aattctacga ccagtctaaa aagcgcctga
4621 attcgcgacc ttctcgttac tgacaggaaa atgggccatt ggcaaccagg gaaagatgaa
4681 cgtgatgatg ttcacaattt gctgaattgt ggtgatgtga tgctcaccgc atttcctgaa
4741 aattcacgct gtatcttgaa aaatcgacgt tttttacgtg gttttccgtc gaaaatttaa
4801 ggtaagaacc tgacctcgtg attactattt cgccgtgttg acgacatcag gaggccagta
4861 tgaccgtatt acatagtgtg gatttttttc cgtctggtaa cgcgtccgtg gcgatagaac
4921 cccggctccc gcaggcggat ttcctgaac atcatcatga ttttcatgaa attgtgattg
4981 tcgaacatgg cacgggtatt catgtgttta atgggcagcc ctataccatc accggtggca
5041 cggtctgttt cgtacgcgat catgatcggc atctgtatga acataccgat aatctgtgtc
5101 tgaccaatgt gctgtatcgc tcgccggatc gatttcagtt tctcgccggg ctgaatcagt
5161 tgctgccaca agagctggat gggcagtatc cgtctcactg gcgcgttaac cacagcgtat
5221 tgcagcaggt gcgacagctg gttgcacaga tggaacagca ggaaggggaa aatgatttac
5281 cctcgaccgc cagtcgcgag atcttgttta tgcaattact gctcttgctg cgtaaaagca
5341 gtttgcagga gaacctggaa aacagcgcat cacgtctcaa cttgcttctg gcctggctgg
5401 aggaccattt tgccgatgag gtgaattggg atgccgtggc ggatcaattt tctctttcac
5461 tgcgtacgct acatcggcag cttaagcagc aaacgggact gacgcctcag cgatacctga
5521 accgcctgcg actgatgaaa gcccgacatc tgctacgcca cagcgaggcc agcgttactg

5581 acatcgccta tcgctgtgga ttcagcgaca gtaaccactt ttcgacgctt tttcgccgag
5641 agtttaactg gtcaccgcgt gatattcgcc agggacggga tggctttctg caataacgcg
5701 aatcttctca acgtatttgt acgccatatt gcgaataatc aacttcgttc tctggccgag
5761 gtagccacgg tggcgcatca gttaaaactt ctcaaagatg atttttttgc cagcgaccag
5821 caggcagtcg ctgtggctga ccgttatccg caagatgtct ttgctgaaca tacacatgat
5881 ttttgtgagc tggtgattgt ctggcgcggt aatggcctgc atgtactcaa cgatcgccct
5941 tatcgcatta cccgtggcga tctcttttac attcatgctg acgataaaca ctcctacgct
6001 tccgttaacg atctggtttt gcagaatatt atttattgcc cggagcgtct gaagctgaat
6061 cttgactggc agggggcgat tccgggattt aacgccagcg cagggcaacc acactggcgc
6121 ttaggtagca tggggatggc gcaggcgcgg caggttatcg gtcagcttga gcatgaaagt
6181 agtcagcatg tgccgtttgc taacgaaatg gctgagttgc tgttcgggca gttggtgatg
6241 ttgctgaatc gccatcgtta caccagtgat tcgttgccgc caacatccag cgaaacgttg
6301 ctggataagc tgattacccg gctggcggct agcctgaaaa gtccctttgc gctggataaa
6361 ttttgtgatg aggcatcgtg cagtgagcgc gttttgcgtc agcaatttcg ccagcagact
6421 ggaatgacca tcaatcaata tctgcgacag gtcagagtgt gtcatgcgca atatcttctc
6481 cagcatagcc gcctgttaat cagtgatatt tcgaccgaat gtggctttga agatagtaac
6541 tatttttcgg tggtgtttac ccgggaaacc gggatgacgc ccagccagtg gcgtcatctc
6601 aattcgcaga aagattaact cgaggtgaag acgaaagggc ctcgtgatac gcctattttt
6661 ataggttaat gtcatgataa taatggtttc ttagacgtca ggtggcactt ttcggggaaa
6721 tgtgcgcgga acccctattt gtttattttt ctaaatacat tcaaatatgt atccgctcat
6781 gagacaataa ccctgataaa tggtaccgag ctcgaattcg cgcggccgcg gtaccgggcc
6841 cgtcgggatc cggtgattga ttgagcaagc tttatgcttg taaaccgttt tgtgaaaaaa
6901 tttttaaaat aaaaaagggg acctctaggg tccccaatta gaattggccg cggcgttgtg
6961 acaatttacc gaacaactcc gcggccggga agccgatctc ggcttgaacg aattgttagg
7021 tggcggtact tgggtcgata tcaaagtgca tcacttcttc ccgtatgccc aactttgtat
7081 agagagccac tgcgggatcg tcaccgtaat ctgcttgcac gtagatcaca taagcaccaa
7141 gcgcgttggc ctcatgcttg aggagattga tgagcgcggt ggcaatgccc tgcctccggt
7201 gctcgccgga gactgcgaga tcatagatat agatctcact acgcggctgc tcaaacttgg
7261 gcagaacgta agccgcgaga gcgccaacaa ccgcttcttg gtcgaaggca gcaagcgcga
7321 tgaatgtctt actacggagc aagttcccga ggtaatcgga gtccggctga tgttgggagt
7381 aggtggctac gtctccgaac tcacgaccga aaagatcaag agcagcccgc atggatttga
7441 cttggtcagg gccgagccta catgtgcgaa tgatgcccat acttgagcca cctaactttg
7501 ttttagggcg actgccctgc tgcgtaacat cgttgctgct gcgtaacatc gttgctgctc
7561 cataacatca aacatcgacc cacggcgtaa cgcgcttgct gcttggatgc ccgaggcata
7621 gactgtacaa aaaaacagtc ataacaagcc atgaaaaccg ccactgcgcc gttaccaccg
7681 ctgcgttcgg tcaaggttct ggaccagttg cgtgagcgca tacgctactt gcattacagt
7741 ttacgaaccg aacaggctta tgtcaattcg taattgggga ccctagaggt cccctttttt
7801 attttaaaaa ttttttcaca aaacggttta caagcataaa gcttgctcaa tcaatcaccg
7861 gatccccgac tctagtcgac ctgcaggcca accagataag tgaaatctag ttccaaacta
7921 ttttgtcatt tttaattttc gtattagctt acgacgctac acccagttcc catctatttt
7981 gtcactcttc cctaaataat ccttaaaaac tccatttcca cccctcccag ttcccaacta
8041 ttttgtccgc ccacagcggg gcatttttct tcctgttatg tttgggcgag ctcgaattca
8101 ctggccgtcg ttttacaacg tcgtgactgg gaaaaccctg gcgttaccca acttaatcgc
8161 cttgcagcac atcccccttt cgccagctgg cgtaatagcg aagaggcccg caccgatcgc
8221 ccttcccaac agttgcgcag cctgaatggc gaatggcgcc tgatgcggta ttttctcctt
8281 acgcatctgt gcggtatttc acaccgcata tggtgcactc tcagtacaat ctgctctgat
8341 gccgcatagt taagccagcc ccgacacccg ccaacacccg ctgacgcgcc ctgacgggct
8401 tgtctgctcc cggcatccgc ttacagacaa gctgtgaccg tctccgggag ctgcatgtgt
8461 cagaggtttt caccgtcatc accgaaacgc gcgagacgaa agggcctcgt gatacgccta
8521 tttttatagg ttaatgtcat gataataatg gtttcttaga cgtcaggtgg cacttttcgg
8581 ggaaatgtgc gcggaacccc tatttgttta tttttctaaa tacattcaaa tatgtatccg
8641 ctcatgagac aataaccctg ataaatgctt caataatatt gaaaaaggaa gagtatgagt
8701 attcaacatt tccgtgtcgc ccttattccc ttttttgcgg cattttgcct tcctgttttt
8761 gctcacccag aaacgctggt gaaagtaaaa gatgctgaag atcagttggg tgcacgagtg
8821 ggttacatcg aactggatct caacagcggt aagatccttg agagttttcg ccccgaagaa
8881 cgttttccaa tgatgagcac ttttaaagtt ctgctatgtg gcgcggtatt atcccgtatt
8941 gacgccgggc aagagcaact cggtcgccgc atacactatt ctcagaatga cttggttgag
9001 tactcaccag tcacagaaaa gcatcttacg gatggcatga cagtaagaga attatgcagt
9061 gctgccataa ccatgagtga taacactgcg gccaacttac ttctgacaac gatcggagga
9121 ccgaaggagc taaccgcttt tttgcacaac atgggggatc atgtaactcg ccttgatcgt
9181 tgggaaccgg agctgaatga agccatacca aacgacgagc gtgacaccac gatgcctgta
9241 gcaatggcaa caacgttgcg caaactatta actggcgaac tacttactct agcttcccgg
9301 caacaattaa tagactggat ggaggcggat aaagttgcag gaccacttct gcgctcggcc
9361 cttccggctg gctggtttat tgctgataaa tctggagccg gtgagcgtgg gtctcgcggt
9421 atcattgcag cactggggcc agatggtaag ccctcccgta tcgtagttat ctacacgacg
9481 gggagtcagg caactatgga tgaacgaaat agacagatcg ctgagatagg tgcctcactg
9541 attaagcatt ggtaactgtc agaccaagtt tactcatata tactttagat tgatttaaaa

9601 cttcattttt aatttaaaag gatctaggtg aagatccttt ttgataatct catgaccaaa
9661 atcccttaac gtgagttttc gttccactga gcgtcagacc ccgtagaaaa gatcaaagga
9721 tcttc

#### 13. pAKN69-rcgA

**Description** pAKN69-based constitutive expression construct for rcgA (PA14_27700) under the synthetic PA1/04/03 promoter (Gm(R), Cm(R)).

**Length** 7,798 bp, circular

**Source** This paper

##### Features

| Location | Type | Feature |
|---|---|---|
| 3188..3694 | gene | aacC1 (gentamicin resistance) |
| 4091..4169 | promoter | PA1/04/03 promoter |
| 4189..4194 | misc_feature | ribosome binding site |
| 4203..4928 | CDS | rcgA (PA14_27700), 241 aa |
| 4932..5030 | terminator | terminator |
| 5070..5729 | gene | cat (chloramphenicol resistance) |
| 5788..5884 | terminator | terminator |

##### Sequence (7,798 bp)

```
   1 accgcgccct cgctgacctc tgcgcgacgc tcgaacggga ctacgggctt gagcgtgaca
  61 atcacgaaac gcggcagcgc gtttccgaga accgcgcgaa cgacatggag cggcacgcgg
 121 gcgtggaaag cctggtcggc tggatcctct agagtcctgc attaatgaat cggccaacgc
 181 gcggggagag gcggtttgcg tattgggcgc tcttccgctt cctcgctcac tgactcgctg
 241 cgctcggtcg ttcggctgcg gcgagcggta tcagctcact caaaggcggt aatacggtta
 301 tccacagaat caggggataa cgcaggaaag aacatgtgag caaaaggcca gcaaaaggcc
 361 aggaaccgta aaaaggccgc gttgctggcg tttttccata ggctccgccc ccctgacgag
 421 catcacaaaa atcgacgctc aagtcagagg tggcgaaacc cgacaggact ataaagatac
 481 caggcgtttc cccctggaag ctccctcgtg cgctctcctg ttccgaccct gccgcttacc
 541 ggatacctgt ccgcctttct cccttcggga agcgtggcgc tttctcatag ctcacgctgt
 601 aggtatctca gttcggtgta ggtcgttcgc tccaagctgg gctgtgtgca cgaacccccc
 661 gttcagcccg accgctgcgc cttatccggt aactatcgtc ttgagtccaa cccggtaaga
 721 cacgacttat cgccactggc agcagccact ggtaacagga ttagcagagc gaggtatgta
 781 ggcggtgcta cagagttctt gaagtggtgg cctaactacg gctacactag aagaacagta
 841 tttggtatct gcgctctgct gaagccagtt accttcggaa aaagagttgg tagctcttga
 901 tccggcaaac aaaccaccgc tggtagcggt ggtttttttg tttgcaagca gcagattacg
 961 cgcagaaaaa aaggatctca agaagatcct ttgatctttt ctacggggtc tgacgctcag
1021 tggaacgaaa actcacgtta agggattttg gtcatgagat tatcaaaaag gatcttcacc
1081 tagatccttt taaattaaaa atgaagtttt aaatcaatct aaagtatata tgagtaaact
1141 tggtctgaca gttaccaatg cttaatcagt gaggcaccta tctcagcgat ctgtctattt
1201 cgttcatcca tagttgcctg actccccgtc gtgtagataa ctacgatacg ggagggctta
1261 ccatctggcc ccagtgctgc aatgataccg cgagacccac gctcaccggc tccagattta
1321 tcagcaataa accagccagc cggaagggcc gagcgcagaa gtggtcctgc aactttatcc
1381 gcctccatcc agtctattaa ttgttgccgg gaagctagag taagtagttc gccagttaat
1441 agtttgcgca acgttgttgc cattgctaca ggcatcgtgg tgtcacgctc gtcgtttggt
1501 atggcttcat tcagctccgg ttcccaacga tcaaggcgag ttacatgatc ccccatgttg
1561 tgcaaaaaag cggttagctc nttcggtcct ccgatcgttg tcagaagtaa gttggccgca
1621 gtgttatcac tcatggttat ggcagcactg cataattctc ttactgtcat gccatccgta
1681 agatgctttt ctgtgactgg tgagtactca accaagtcat tctgagaata gtgtatgcgg
1741 cgaccgagtt gctcttgccc ggcgtcaata cgggataata ccgcgccaca tagcagaact
1801 ttaaaagtgc tcatcattgg aaaacgttct tcggggcgaa aactctcaag gatcttaccg
1861 ctgttgagat ccagttcgat gtaacccact cgtgcaccca actgatcttc agcatctttt
1921 actttcacca gcgtttctgg gtgagcaaaa acaggaaggc aaaatgccgc aaaaaaggga
1981 ataagggcga cacggaaatg ttgaatactc atactcttcc tttttcaata ttattgaagc
2041 atttatcagg gttattgtct catgagcgga tacatatttg aatgtattta gaaaaataaa
2101 caaatagggg ttccgcgcac atttccccga aaagtgccac ctgacgtcta agaaaccatt
2161 attatcatga cattaaccta taaaaatagg cgtatcacga ggccctttcg tctcgcgcgt
2221 ttcggtgatg acggtgaaaa cctctgacac atgcagctcc cggagacggt cacagcttgt
2281 ctgtaagcgg atgccgggag cagacaagcc cgtcagggcg cgtcagcggg tgttggcggg
```

2341 tgtcggggct ggcttaacta tgcggcatca gagcagattg tactgagagt gcaccatatg
2401 cggtgtgaaa taccgcacag atgcgtaagg agaaaatacc gcatcaggcg ccattcgcca
2461 ttcaggctgc gcaactgttg ggaagggcga tcggtgcggg cctcttcgct attacgccag
2521 ctggcgaaag ggggatgtgc tgcaaggcga ttaagttggg taacgccagg gttttcccag
2581 tcacgacgtt gtaaaacgac ggccagtgaa ttcgagctcg cccaaacata acaggaagaa
2641 aaatgccccg ctgtgggcgg acaaaatagt tgggaactgg gaggggtgga aatggagttt
2701 ttaaggatta tttagggaag agtgacaaaa tagatgggaa ctgggtgtag cgtcgtaagc
2761 taatacgaaa attaaaaatg acaaaatagt ttggaactag atttcactta tctggttggc
2821 ctgcaggtcg actagagtcg gggatccggt gattgattga gcaagcttta tgcttgtaaa
2881 ccgttttgtg aaaaaatttt taaaataaaa aaggggacct ctagggtccc caattacgaa
2941 ttgacataag cctgttcggt tcgtaaactg taatgcaagt agcgtatgcg ctcacgcaac
3001 tggtccagaa ccttgaccga acgcagcggt ggtaacggcg cagtggcggt tttcatggct
3061 tgttatgact gtttttttgt acagtctatg cctcgggcat ccaagcagca agcgcgttac
3121 gccgtgggtc gatgtttgat gttatggagc agcaacgatg ttacgcagca gcaacgatgt
3181 tacgcagcag ggcagtcgcc ctaaaacaaa ggtaggtggt ccaagtatgg gccatcattc
3241 gcacatgtag gctcggccct gaccaagtca aatccatgcg ggctgctctt gatcatttcg
3301 gtcgtgagtt cggagacgta gccacctact cccaacatca gccgggactc cgataccctc
3361 gggaacatgc tccgtagtaa gacattcatc gcgcttgact gccttcgacc aagaagcggt
3421 tgttggcgct ctcgcggctt acgttctgcc caagtttgag cagccgcgta gtgagatcta
3481 tatctatgat ctcgcagtct ccggcgagca ccggaggcag ggcattgcca ccgcgctcat
3541 caatctcctc aagcatgagg ccaacgcgct tggtgcttat gtgatgtacg tgcaagcaga
3601 ttacggtgac gatcccgcag tggctctcta tacaaagttg ggcatacggg aagaagtgat
3661 gcactttgat atcgacccaa gtaccgccac ctaacaattc gttcaagccg agatcggctt
3721 cccggccgcg gagttgttcg gtaaattgtc acaacgccgc ggccaattct aattggggac
3781 cctagaggtc cccttttttta ttttaaaaat tttttcacaa aacggtttac aagcataaag
3841 cttgctcaat caatcaccgg atcccgacgg gcccggtacc gcggccgcgc gaattcgagc
3901 tcggtaccat ttatcagggt tattgtctca tgagcggata catatttgaa tgtatttaga
3961 aaaataaaca aataggggtt ccgcgcacat ttccccgaaa agtgccacct gacgtctaag
4021 aaaccattat tatcatgaca ttaacctata aaaataggcg tatcacgagg ccctttcgtc
4081 ttcacctcga gaaaatttat caaaaagagt gttgacttgt gagcggataa caatgatact
4141 tagattcaat tgtgagcgga taacaatttc acacatctag aattaaagag gagaaattaa
4201 gcatgcactc gaacgagtca acggaagacc gctgttcacg ccactacacc ctgctgcaac
4261 ggtgcccgct gctgcacggg ctgcccgatc tgctgcggca cgtggcgacc catgcgtcgg
4321 agcgccggtt gcgcgacaag gacgtcctgt tcctcaagca cgaccctgaa gacttcatcg
4381 gcctggtcct cggcggacac ctctataccg tgctctacgg gccggacggg cgggagctga
4441 tcgtcaatga tatccgtcct ggcgaagtga tcggggaaac cgcgctgatc cagccgaacc
4501 gccgcgagac ctctgcctac gcctccggcg cggcgtcggt gctgatcctc caccgacgtc
4561 acttcgcccc gttgatcgcc gacacggaat tcctcaaccg cgcgctggcc ctcctctgtt
4621 cgcgcctgcg cgaagcctcc ggtttcgtcg agagggtctg cctgcatcgc ctggagtcgc
4681 gcctggcgcg ccatctcctg gccagcctgg cgagcgaaca acccgaggga gcctgcgcga
4741 cgctaccgac caaccagagc atcctagcag cgatgctcaa cgccagccgg ccgaagctca
4801 acgcgcaact gcaaatatgg aaacgcgaag ggctgatcag ctgccgcaac gaccggatcc
4861 tgatcaacga cctcggtcgc ctgcgacgca aggccgaact accagccgcg cccctgtcac
4921 ccaggtaatt ggactcctgt tgatagatcc agtaatgacc tcagaactcc atctggattt
4981 gttcagaacg ctcggttgcc gccgggcgtt ttttattggt gagaatccaa gctagcttgg
5041 cgagattttc aggagctaag gaagctaaaa tggagaaaaa aatcactgga tataccaccg
5101 ttgatatatc ccaatggcat cgtaaagaac attttgaggc atttcagtca gttgctcaat
5161 gtacctataa ccagaccgtt cagctggata ttacggcctt tttaaagacc gtaaagaaaa
5221 ataagcacaa gttttatccg gcctttattc acattcttgc ccgcctgatg aatgctcatc
5281 cggaatttcg tatggcaatg aaagacggtg agctggtgat atgggatagt gttcaccctt
5341 gttacaccgt tttccatgag caaactgaaa cgttttcatc gctctggagt gaataccacg
5401 acgatttccg gcagtttcta cacatatatt cgcaagatgt ggcgtgttac ggtgaaaacc
5461 tggcctattt ccctaaaggg tttattgaga atatgttttt cgtctcagcc aatccctggg
5521 tgagtttcac cagttttgat ttaaacgtgg ccaatatgga caacttcttc gcccccgttt
5581 tcaccatggg caaatattat acgcaaggcg acaaggtgct gatgccgctg gcgattcagg
5641 ttcatcatgc cgtttgtgat ggcttccatg tcggcagaat gcttaatgaa ttacaacagt
5701 actgcgatga gtggcagggc ggggcgtaat ttttttaagg cagttattgg tgcccttaaa
5761 cgcctggggt aatgactctc tagcttgagg catcaataaa acgaaaggct cagtcgaaag
5821 actgggcctt tcgttttatc tgttgtttgt cggtgaacgc tctcctgagt aggacaaatc
5881 cgccgctagg agcttgcggc cgccttaaga cgcgtggatc cccgggaatt tcgaccccac
5941 gcccctcttt aatacgacgg gcaatttgca cttcagaaaa tgaagagttt gctttagcca
6001 taacaaaagt ccagtatgct ttttcacagc ataactggac tgatttcagt ttacaactat
6061 tctgtctagt ttaagacttt attgtcatac tttagatcta ttttgttcag tttaagactt
6121 tattgtccgc ccacacccgc ttacgcaggg catccattta ttactcaacc gtaaccgatt
6181 ttgccaggtt acgcggctgg tcctctagag gatccttttt gtccggtgtt gggttgaagg
6241 tgaagccggt cggggccgca gcgggggccc ggcttttcag ccttgccccc ctgcttcggc
6301 cgccgtggct ccggcgtctt gggtgccggc gcgggttccg cagccttggc ctgcggtgcg

6361 ggcacatcgg cgggcttggc cttgatgtgc cgcctggcgt gcgagcggaa cgtctcgtag
6421 gagaacttga ccttccccgt ttcccgcatg tgctcccaaa tggtgacgag cgcatagccg
6481 gacgctaacg ccgcctcgac atccgccctc accgccagga acgcaaccgc agcctcatca
6541 cgccggcgct tcttggccgc gcgggattca acccactcgg ccagctcgtc ggtgtagctc
6601 tttggcatcg tctctcgcct gtcccctcag ttcagtaatt tcctgcattt gcctgtttcc
6661 agtcggtaga tattccacaa aacagcaggg aagcagcgct tttccgctgc ataaccctgc
6721 ttcggggtca ttatagcgat tttttcggta tatccatcct ttttcgcacg atatacagga
6781 ttttgccaaa gggttcgtgt agactttcct tggtgtatcc aacggcgtca gccgggcagg
6841 ataggtgaag taggcccacc cgcgagcggg tgttccttct tcactgtccc ttattcgcac
6901 ctggcggtgc tcaacgggaa tcctgctctg cgaggctggc cggctaccgc cggcgtaaca
6961 gatgagggca agcggatggc tgatgaaacc aagccaacca ggaagggcag cccacctatc
7021 aaggtgtact gccttccaga cgaacgaaga gcgattgagg aaaaggcggc ggcggccggc
7081 atgagcctgt cggcctacct gctggccgtc ggccagggct acaaaatcac gggcgtcgtg
7141 gactatgagc acgtccgcga gctggcccgc atcaatggcg acctgggccg cctgggcggc
7201 ctgctgaaac tctggctcac cgacgacccg cgcacggcgc ggttcggtga tgccacgatc
7261 ctcgccctgc tggcgaagat cgaagagaag caggacgagc ttggcaaggt catgatgggc
7321 gtggtccgcc cgagggcaga gccatgactt ttttagccgc taaaacggcc ggggggtgcg
7381 cgtgattgcc aagcacgtcc ccatgcgctc catcaagaag agcgacttcg cggagctggt
7441 gaagtacatc accgacgagc aaggcaagac cgagcgcctg ggtcacgtgc gcgtcacgaa
7501 ctgcgaggca aacaccctgc ccgctgtcat ggccgaggtg atggcgaccc agcacggcaa
7561 cacccgttcc gaggccgaca agacctatca cctgctggtt agcttccgcg cgggagagaa
7621 gcccgacgcg gagacgttgc gcgcgattga ggaccgcatc tgcgctgggc ttggcttcgc
7681 cgagcatcag cgcgtcagtg ccgtgcatca cgacaccgac aacctgcaca tccatatcgc
7741 catcaacaag attcacccga cccgaaacac catccatgag ccgtatcggg cctaccgc
